# Evolution of tissue-specific expression of ancestral genes across vertebrates and insects

**DOI:** 10.1101/2022.11.14.516384

**Authors:** Federica Mantica, Luis P. Iñiguez, Yamile Marquez, Jon Permanyer, Antonio Torres-Mendez, Josefa Cruz, Xavi Franch-Marro, Frank Tulenko, Demian Burguera, Stephanie Bertrand, Toby Doyle, Marcela Nouzova, Peter Currie, Fernando G. Noriega, Hector Escriva, Maria Ina Arnone, Caroline B Albertin, Karl R Wotton, Isabel Almudi, David Martin, Manuel Irimia

## Abstract

Regulation of gene expression is arguably the main mechanism contributing to tissue phenotypic diversity within and between species. Here, we assembled an extensive transcriptomic dataset covering twenty bilaterian species and eight tissues, selecting a specular phylogeny that allowed both the combined and parallel investigation of gene expression evolution between vertebrates and insects. We specifically focused on widely conserved ancestral genes, identifying strong cores of pan-bilaterian tissue-specific genes and even larger groups that diverged to define vertebrate and insect tissues. Systematic inferences of tissue-specificity gains and losses show that nearly half of all ancestral genes have been recruited into tissue-specific transcriptomes. This occurred during both ancient and, especially, recent bilaterian evolution, with several gains being associated with the emergence of unique phenotypes. Such pervasive evolution of tissue-specificity was linked to gene duplication coupled with specialization, including an unappreciated prolonged effect of whole genome duplications during recent vertebrate evolution.

## Introduction

Fossil records reconstruct the image of the last common ancestor (LCA) of all bilaterian animals as a small, marine creature ^1^ crawling on the seafloor approximately 700 million years ago (MYA) ^1^. Despite its apparent simplicity, this ancestral organism introduced remarkable biological novelties, including an original body plan, defined by two perpendicular symmetry axes, and a third embryonic germ layer ^2^. In addition, it already possessed an ancestral form of the main tissue types that are homologous across extant bilaterian species, including a nervous system, skeletal muscle, female and male gonads, through guts and an excretory system ^3^. How did this ancient organism specify such a great variety of biological structures? Since all its cells shared the same genome, gene expression regulation was likely key for the generation of unique transcriptomes across these ancestral tissue types, and consequently for the emergence of their distinctive biological functions.

The bilaterian ancestor gave rise to the vast majority of extant animals, where the original body plan and tissues have been greatly diversified and modified. Determinants of animal evolution include changes in gene complements (i.e. gene gains/losses and gene duplications), divergence of protein-coding sequences ^4–6^ and regulatory changes in gene expression ^7^. In fact, as the generation of tissue-specific transcriptomes is key for defining distinct tissue types (i.e., intra-species diversity), their evolutionary remodeling is arguably the most crucial determinant of phenotypic variation among species (i.e. inter-species diversity) ^8,9^. Importantly, major transcriptome remodeling often involves conserved genes ^10^. Examples of such occurrences underlie important phenotypic novelties in key bilaterian lineages: the vertebrate endocrine pancreas emerged following the recruitment of ancestrally neural-specific genes ^11^, and it has been suggested that insect wings evolved upon co-option of expression of ancient genes originally involved in gill specification and proximal leg segments ^12–14^. Still, even if remarkable cases linked to biological novelties have been identified, the specific role that the evolution of expression of ancestral genes played in shaping homologous, yet often highly divergent, tissues between distant bilaterian lineages has never been thoroughly assessed.

Comparative transcriptomic data offer valuable insights into ancestral molecular states (e.g., ancestral expression profiles) and their subsequent history. Here, we studied the evolution of tissue-specific transcriptomes based on an extensive RNA sequencing (RNA-seq) dataset covering eight tissue types from twenty bilaterian species, including novel data for fifteen of them. Compared to previous studies of transcriptome evolution, mostly focused on mammals, this dataset extends the phylogenetic coverage beyond vertebrates and provides the first comparative framework of this scale for insects. Vertebrates and insects were selected as focus clades because they include highly accessible organisms with a relatively similar body plan compared to other lineages (e.g., most lophotrochozoans). However, they also reached some opposite evolutionary solutions in terms of structural organization (e.g., dorsal/ventral positioning of the spinal/nerve cords in vertebrates and insects, respectively ^15^), and show remarkably different molecular and genomic evolutionary rates ^16,17^. We selected a symmetric phylogeny for both species groups, which allowed us to perform sound ancestral inferences as well as to uncover parallel, convergent and divergent evolutionary trajectories in comparable evolutionary/phylogenetic nodes of these early diverging bilaterian lineages. First, we characterized global gene expression patterns and reconstructed ancestral bilaterian tissue-specific modules that are still widely conserved across extant species. Second, we systematically inferred gains and losses of tissue-specific expression throughout our selected bilaterian phylogeny. Lastly, we characterized these inferred tissue-specificity gains from the mechanistic and functional perspective. Overall, our work sheds light on the highly plastic nature of deeply conserved genes in terms of tissue-specific expression patterns, which we find to be tightly linked to gene duplication, specialization and the emergence of unique tissue-related phenotypes.

## Results

### Characterization of global patterns of gene expression across bilaterian tissues

In order to reliably investigate the evolution of tissue-specific transcriptomes in two key bilaterian lineages, we selected twenty representative species (eight gnathostome vertebrates, eight insects and two pairs of relative outgroups) evenly divided into two monophyletic branches with specular phylogenetic structures (**Fig. 1a**). After correcting for broken/chimeric genes and enriching the annotations of the majority of the species (see **Supplementary Methods Extended Data Fig. 1a-d** and **Supplementary Table 1**), we derived gene orthology relationships among all of them and isolated 7,178 bilaterian-conserved gene orthogroups, which were unambiguously present in the bilaterian LCA (i.e., conserved in at least 12/20 species. See **Supplementary Methods**; **Extended Data Fig. 1e,f** and **Supplementary Table 2)**. In addition, as several comparisons required the selection of one representative ortholog per species, we generated the most suitable one-to-one orthogroups for each set of analyses (i.e., best-ancestral and best-TS orthogroups; see **Extended Data Fig. 2** and **Methods**) by applying distinct filters to the original bilaterian-conserved orthogroups. We then assembled an extensive bulk RNA-seq dataset covering up to eight tissues in all species (**Fig. 1a** and **Supplementary Table 3**). Notably, while some of the included tissue types (neural, testis, ovaries, muscle and excretory system) had been considered in previous studies of gene expression evolution among bony vertebrate species ^18–21^, others (digestive tract, epidermis and adipose) have never been analyzed in such a context. In total, we generated 89 RNA-seq samples across 15 species, which we combined with publicly available data into a final dataset of 346 RNA-seq meta-samples (see **Methods** for meta-samples definition), including up to three meta-samples for each tissue and species (**Fig. 1a** and **Supplementary Fig. 1**).

**Fig. 1:**
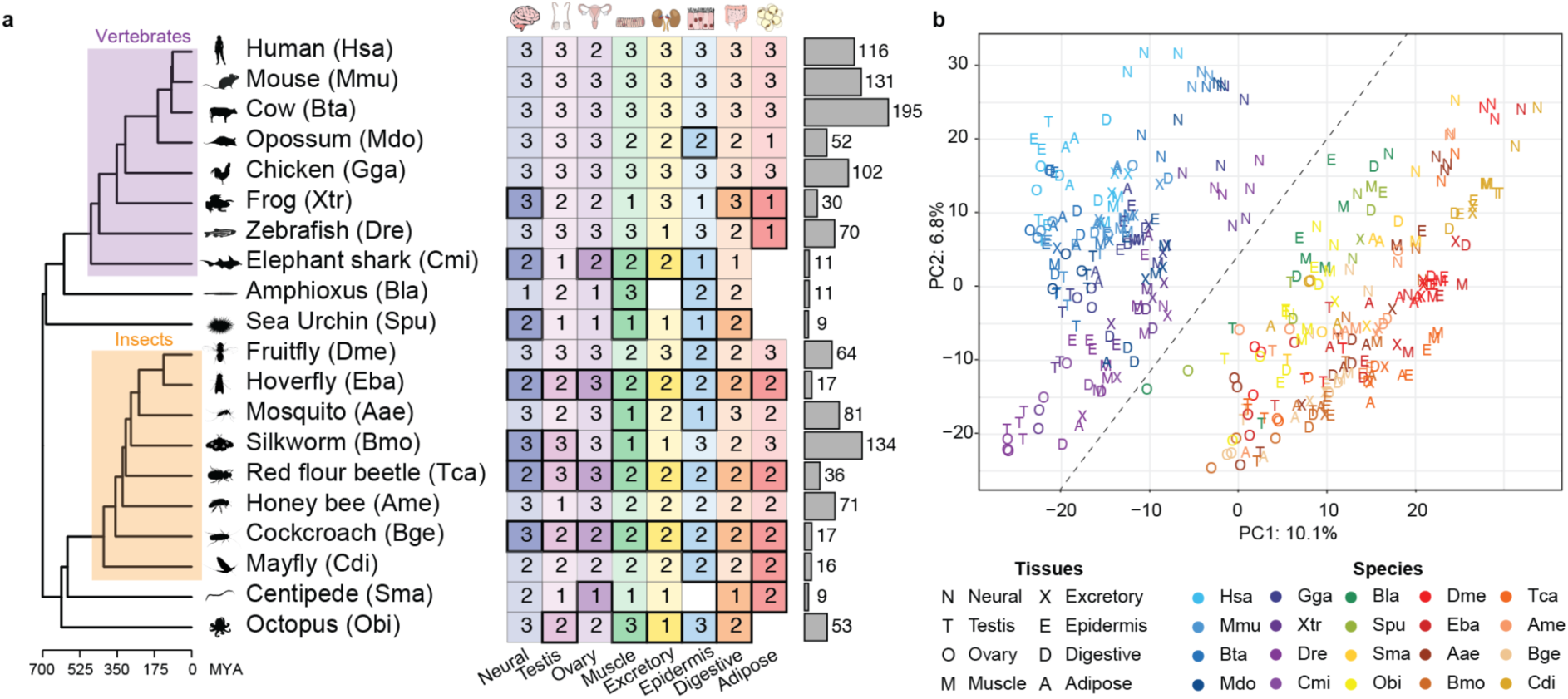
Dataset overview and global patterns of gene expression across bilaterian tissues. **a.** RNA-seq dataset overview. Left: phylogenetic tree including the common names and scientific acronyms of the 20 bilaterian species considered in this study. Evolutionary distances were derived from ^81^ (MYA: million years ago) and animal silhouettes downloaded from http://phylopic.org/ (see **Acknowledgements** for credits to Phylopic icons). Center: scheme of RNA-seq meta-samples. The number of meta-samples for each species (rows) and tissue (columns) is reported. The cell color corresponds to the tissue, while its intensity and border thickness distinguishes between cases where at least one RNA-seq sample has been generated for this project (full color; thick borders) from cases where all the included samples are publicly available (transparent color; thin borders). Right: barplot with the total number of processed RNA sequencing (RNA-seq) samples per species. **b.** Coordinates of the first (PC1; x axis) and second (PC2; y axis) principal components from a principal component analysis (PCA) performed on the best-ancestral orthogroups normalized expression matrix (see **Methods**). Only the 2,436 best-ancestral orthogroups conserved in all species were considered. Tissue identity is represented by letters and species by colors. The percentage of variance explained by each PC is reported on the relative axis.

In our dataset, the first two components of a principal component analysis (PCA) showed a clear distinction between meta-samples from vertebrates and non-vertebrates (including all outgroups), independently from their tissue identity (**Fig. 1b**; see **Methods** for normalization procedure). The same pattern is observed when controlling for potential batch effects (**Supplementary Fig. 2a-f**) or using alternative strategies for the selection and expression quantification of the best-ancestral orthogroups (**Supplementary Fig. 2g-j**). Thus, this separation probably emerged because of some intrinsic differences between vertebrates and insects, which we further investigated by performing clade-specific PCAs (see **Methods**). The first two components of the vertebrate PCA outlined groups of meta-samples of the same tissue origin (specially for neural and testis; **Supplementary Fig. 3a,b**), confirming the pattern characterized by previous studies ^18,21,22^ and validating the idea of a conserved, tissue-related transcriptomic signature that prevails over the species identity. On the other hand, the first two components of the insect PCA were dominated by species clustering (**Supplementary Fig. 3c**), with successive components separating species according to their evolutionary distances (**Supplementary Fig. 3d**). Altogether, these results thus suggest faster evolutionary rates within and between insects compared to vertebrates, as reflected by the vertebrate/non-vertebrate meta-samples separation in the first two components of the bilaterian PCA (**Fig. 1b**). Nevertheless, some subsequent components of this PCA significantly separated groups of meta-samples based on their tissue of origin, starting with neural and testis (**Extended Data Fig. 3a-c**). Thus, we performed a z-score transformation of gene expression values within species (see **Methods**) in order to compare their relative expression profiles across tissues. Upon this transformation, we obtained clusters largely corresponding to tissue groups (**Extended Data Fig. 3d**), which suggested at least partial conservation of ancestral bilaterian tissue-specific expression modules.

### Reconstruction of ancestral bilaterian tissue-specific expression modules

In order to characterize ancestral tissue-specific expression modules that are still widely conserved across extant bilaterians, we implemented a strategy based on a sparse partial least square discriminant analysis (sPLS-DA) ^23^, which allowed us to isolate the orthogroups with the most distinctive expression profiles in each tissue (compared to the others) across all species (see **Methods**, **Fig. 2a** and **Extended Data Fig. 4a-f**). The sPLS-DA approach was set up to produce eight sets (one per tissue type) of ancestral orthogroups with conserved tissue-specific expression, which overall comprised 506 (∼7%) of all bilaterian-conserved orthogroups (**Fig. 2b** and **Supplementary Table 4)**. A PCA performed on these ancestral orthogroups (see **Methods** and **Supplementary Fig. 4a**), or subsets of these orthogroups of the same size across tissues (**Supplementary Fig. 4b**), showed aggregation by tissue type, with a clear separation between all neural and non-neural meta-samples along the first principal component.

**Fig. 2:**
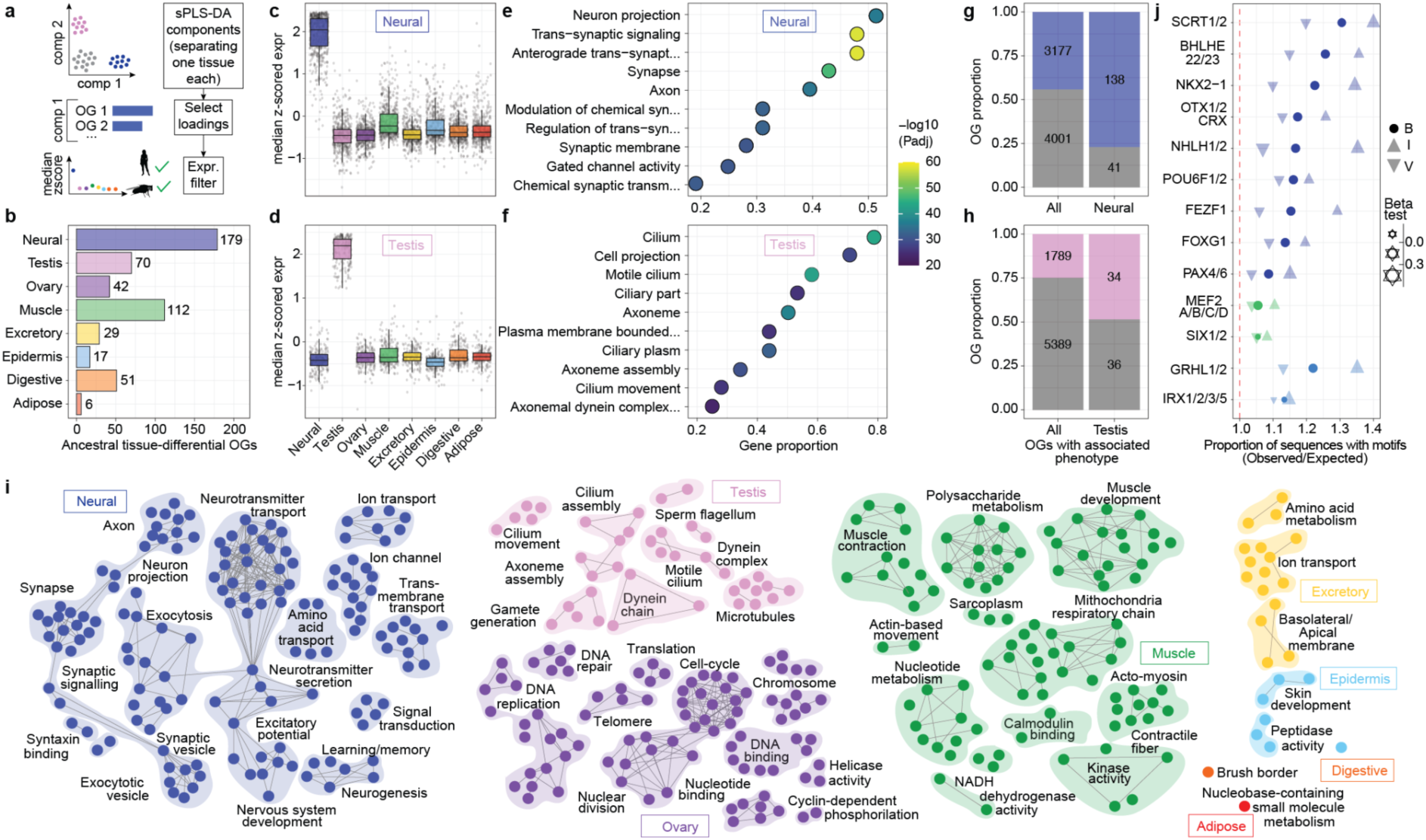
Reconstruction of ancestral bilaterian tissue-specific expression modules. **a.** Scheme depicting the procedure for the definition of the ancestral tissue-specific modules. We performed a sparse partial least square discriminant analysis (sPLS-DA) on the expression of the best-ancestral orthogroups, where each component specifically separated the meta-samples of one tissue group from all the others. We selected loadings of each component, which corresponded to orthogroups with the most distinctive expression profile in the isolated tissue compared to the rest. Finally, we required these orthogroups to show the highest median expression (after z-scoring expression within each species) in that particular tissue among both vertebrates and insects. **b.** Number of best-ancestral orthogroups (OGs) included in each ancestral tissue-specific module. **c,d.** Expression profiles across tissues of best-ancestral orthogroups in the ancestral neural-(c) and testis-(d) specific modules. Expression values were first z-scored by species, and each dot represents the median expression among vertebrates, insects or outgroups. **e,f.** Top 20 most significantly enriched GO categories in the ancestral neural-(e) and testis-(f) specific modules. Only GOs containing at least 5 orthogroups in the tested set were considered. **g,h**: Proportion of all bilaterian-conserved orthogroups (left) and ancestral neural-(g) or testis-specific (h) modules (right) associated with experimentally validated phenotypes (in the respective tissue) in mammals and/or fruit fly. **i.** Representation of GO networks of significantly enriched categories for all ancestral tissue-specific modules, where only categories containing at least 5 orthogroups in the tested set were considered (see **Methods**). Each node represents a GO category. **j.** TFs included within each tissue-specific module whose known binding motifs are significantly over-represented in the regulatory regions of the genes in the corresponding module (see **Methods**). Each TF was tested (Fisher’s exact and regression tests) on all sequences (B: bilaterian), only vertebrate (V) or only insect (I) sequences within the module. TFs in each tissue are ordered by the ratio of the proportion of sequences with at least one predicted binding site in the tested module (observed) compared to the proportion in all other bilaterian-conserved genes (expected). The size of each dot reflects the beta from the regression test in the corresponding group, and tissue colors refer to panel b.

We next investigated in detail the orthogroups belonging to the neural and testis modules, whose expression profiles are shown in **Fig. 2c,d** (see **Extended Data Fig. 4g-l** for other modules). The neural module presented strong over-representation of gene ontology (GO) categories related to synaptic transmission, neuronal morphology and other associated terms (**Fig. 2e**), reflecting the high expression conservation of the specialized neuronal gene complement across eumetazoan nervous systems ^24,25^. The testis module showed significant enrichments for cilium and cytoskeleton-related functions (**Fig. 2f**), probably determined by the axoneme, a highly conserved microtubule-based structure located at the core of most bilaterian spermatozoa flagella and indispensable for their mobility ^26^. The relevant role that these ancestral orthogroups likely play in the respective tissue is supported by their significantly greater association with validated neural-or testis-related phenotypes either in mammals or fly, compared to all bilaterian-conserved orthogroups (**Fig. 2g,h** and **Supplementary Table 5**; p-value < 1e-05 for both tissues, Fisher’s exact tests).

In addition to the neural and testis modules, all other sets exhibited GO enrichments coherent with the deep-rooted functions of each tissue, and comparable between the human (**Fig. 2i** and **Supplementary Table 6**) and fruit fly based GO annotations (**Supplementary table 7**). For instance, genes in the ancestral ovary module comprised several key meiotic genes and were enriched in cell cycle and DNA-replication/repair functions (**Supplementary Tables 4,6,7**). Some examples included *CCNB2*, a cyclin necessary for timely oocyte maturation and correct metaphase-to-anaphase transition in mice ^27,28^, *MOS*, a serine-threonine kinase which mediates metaphase II arrest during meiosis and whose deletion causes human female infertility ^29^, and *CPEB*, a protein involved in regulation of translation prior to fertilization ^30^. As another example, the most significant GO categories for the excretory system module, mainly ion transport and amino acid metabolism, reflected the basic shared functions of ultrafiltration-based excretory systems ^31^. Moreover, even in those tissues in which the homology status of the specific cellular components is more ambiguous/complex (e.g., epidermis, digestive system, adipose), we still obtained a few significant enrichments linked to core molecular programs underlying fundamental functions of each tissue type (**Fig. 2i**).

Finally, we investigated which transcription factors (TFs) might have regulated these ancestral modules since the bilaterian LCA. We specifically tested if the TFs included in each module presented a significant over-representation of predicted binding sites in the regulatory regions of all the other genes in the same ancestral set (see **Methods**). We obtained significant results for multiple TFs, comprising several known master regulators of the respective tissues such as PAX4/6 ^32^ or FEZF1 ^33^ in neural, MEF2A-D ^34^ in muscle and GRHL1/2 ^35^ in epidermis. Importantly, over-representation of their binding sites was observed both at the module level (**Fig. 2j**) and within each studied species separately (**Supplementary Fig. 4c**).

In summary, we reconstructed eight confident modules of ancestral orthogroups with highly conserved tissue-specific expression profiles, offering a high-resolution snapshot of the tissue-specific transcriptomes in the bilaterian LCA.

### Pervasive evolution of tissue-specificity of ancestral genes impacts both ancient and recent bilaterian history

In order to study the evolution of tissue-specificity throughout our entire phylogeny, we next used the Tau metric to define genes with tissue-specific expression profiles in all extant species ^36^. The overall proportion of tissue-specific genes (Tau ≥ 0.75) in each species was lower for bilaterian-conserved genes (i.e., belonging to the 7,178 bilaterian-conserved orthogroups) compared to all genes (**Fig. 3a** and **Supplementary Fig. 5**). This was partially expected, as highly conserved genes are usually associated with greater pleiotropic roles. We assigned each bilaterian-conserved, tissue-specific gene to the tissue(s) with the highest relative expression (see **Methods**, **Fig. 3b**, **Extended Data Fig. 2c**, and **Supplementary Fig. 6**), providing a comprehensive characterization of their tissue-specificity across all species and tissues (**Fig. 3c**). Neural-and testis-specific genes were the most abundant throughout our phylogeny, followed by genes with restricted expression in two different tissues (**Fig. 3d**). Overall, the number of tissue-specific genes was significantly higher among vertebrate species (**Fig. 3e**; p-value = 2e-04; Wilcoxon rank-sum test), likely as a consequence of the two whole genome duplications (WGDs) at the base of vertebrates (see below).

**Fig. 3:**
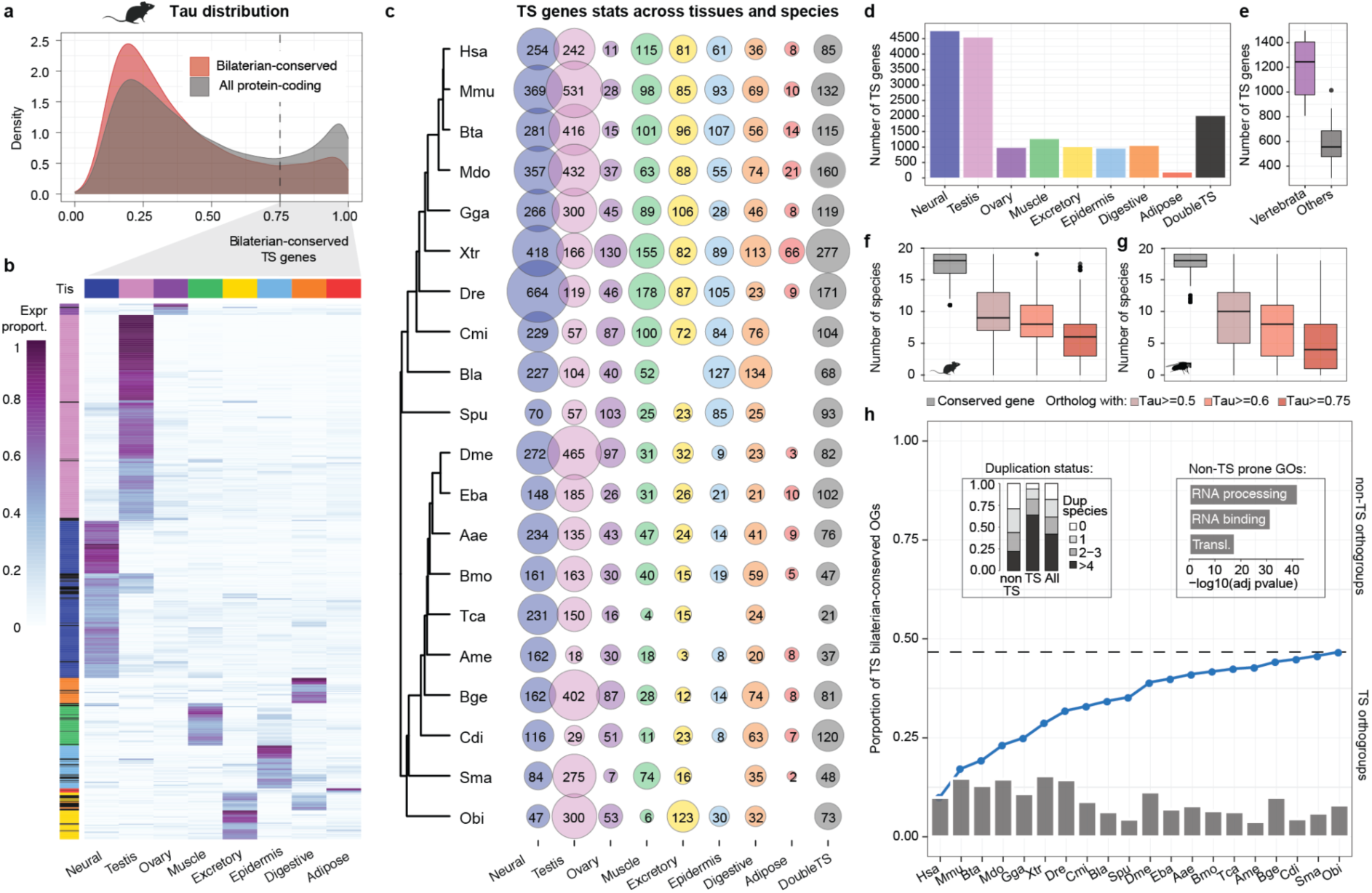
Tissue-specificity patterns across species reveal low conservation of tissue-specific expression profiles. **a.** Tau distributions of all (gray) or bilaterian-conserved (red) mouse protein-coding genes passing the expression cutoff. **b.** Heatmap showing the clustering of mouse bilaterian-conserved, tissue-specific genes (rows) based on their expression proportion (tissue_expr / all_tissue_expr) across tissues (columns). The heatmap was generated by *pheatmap* in R with default parameters, and the complete dendrogram is shown in **Supplementary Fig. 6**. Black indicates double-tissue-specificity. **c,d.** Number of bilaterian-conserved, tissue-specific genes across all species (rows) and tissues (columns) (c) and collapsed by tissue (d). **e.** Distribution of the number of bilaterian-conserved, tissue-specific genes in vertebrates versus all other species (p-value = 2e-04, Wilcoxon rank-sum test). **f,g.** Distribution of the number of species in which mouse (f) or fruit fly (g) bilaterian-conserved, tissue-specific genes have at least one ortholog (gray) and this ortholog(s) has a Tau value higher than a specific cutoff (0.5, 0.6 and 0.75; other shades). **h.** Barplot: proportion of bilaterian-conserved orthogroups including at least one tissue-specific gene in each given species. Line plot: cumulative distribution of the proportion of unique bilaterian-conserved orthogroups containing at least one tissue-specific gene across species. NB: All genes included in the bilaterian-conserved orthogroups were considered for this analysis. The dashed line marks the total proportion. The two boxes include information on the duplication status of the non-tissue-specific (non-TS) and tissue-specific (TS) orthogroups (left) and the top GO enrichments for the non-TS orthogroups (right).

We then set out to investigate the conservation of the identified tissue-specific profiles. Remarkably, we found that these profiles were overall poorly conserved. For instance, for the orthogroups that have at least one gene that is tissue-specific in mouse, only a median of 6 out of the other 19 species had at least one ortholog with Tau ≥ 0.75 (with any associated tissue), and this number merely increased to 9 for Tau ≥ 0.5 (**Fig. 3f**). This pattern was consistent across all species (**Fig. 3g** and **Supplementary Fig. 7**), suggesting that tissue-specificity is highly dynamic and that a high proportion of these tissue-specific expression profiles may have a recent evolutionary origin. In fact, when considering individual species alone, only between 4% and 15% of bilaterian-conserved orthogroups are tissue-specific in each of them (barplot in **Fig. 3h**); however, when all species are taken together, 47% of the bilaterian-conserved orthogroups contain at least one tissue-specific gene (line plot in **Fig. 3h**). In other words, the orthogroups containing tissue-specific genes are widely non-overlapping among species. As expected, orthogroups that are tissue-specific in at least one studied species presented a significantly higher proportion of gene duplications compared to orthogroups that are never tissue-specific (p-value = 2e-04, Fisher’s exact test), which in turn were strongly enriched for housekeeping functions such as RNA processing/binding and translation (extra boxes in **Fig. 3h** and **Supplementary Table 8**).

### Systematic inferences of gains and losses of tissue-specific expression throughout the phylogeny

Next, to investigate dynamics of tissue-specificity evolution in finer detail throughout the entire phylogeny, we adopted a parsimony-based approach to perform a systematic phylogenetic inference of tissue-specificity gains and losses in each tissue (see **Methods**, **Extended Data Fig. 5** and **Supplementary Table 9;** for the rationale behind our chosen procedure, the comparison with other inference methods and their limitations for our dataset, see **Supplementary Discussion**). By definition, given the lack of non-bilaterian outgroups in our phylogeny, we could only infer tissue-specificity gains and losses posterior to the last bilaterian ancestor. Thus, we focused only on the patterns of gain/loss on the phylogenetically equivalent nodes along the two main branches and the tips leading to the extant species (**Fig. 4a**). These inferences, as well as the underlying Tau values, were highly robust to the use of alternative combinations of the original RNA-seq samples (see **Methods**), computed either after averaging the expression of all available samples for each tissue (**Supplementary Fig. 8**) or after randomizing the samples of each tissue across the relative meta-samples (**Supplementary Fig. 9**). Moreover, since we observed that gene expression divergence is subjected to stabilizing selection in all tissues along both phylogenetic branches, and thus mainly evolves following an Ornstein-Uhlenbeck (OU) curve (**Supplementary Fig. 10**), we used an OU-based approach to orthogonally validate our inferred tissue-specificity gains and losses, finding a good overall agreement (see **Extended Data Fig. 6**, **Methods** and **Supplementary Discussion**).

**Fig. 4:**
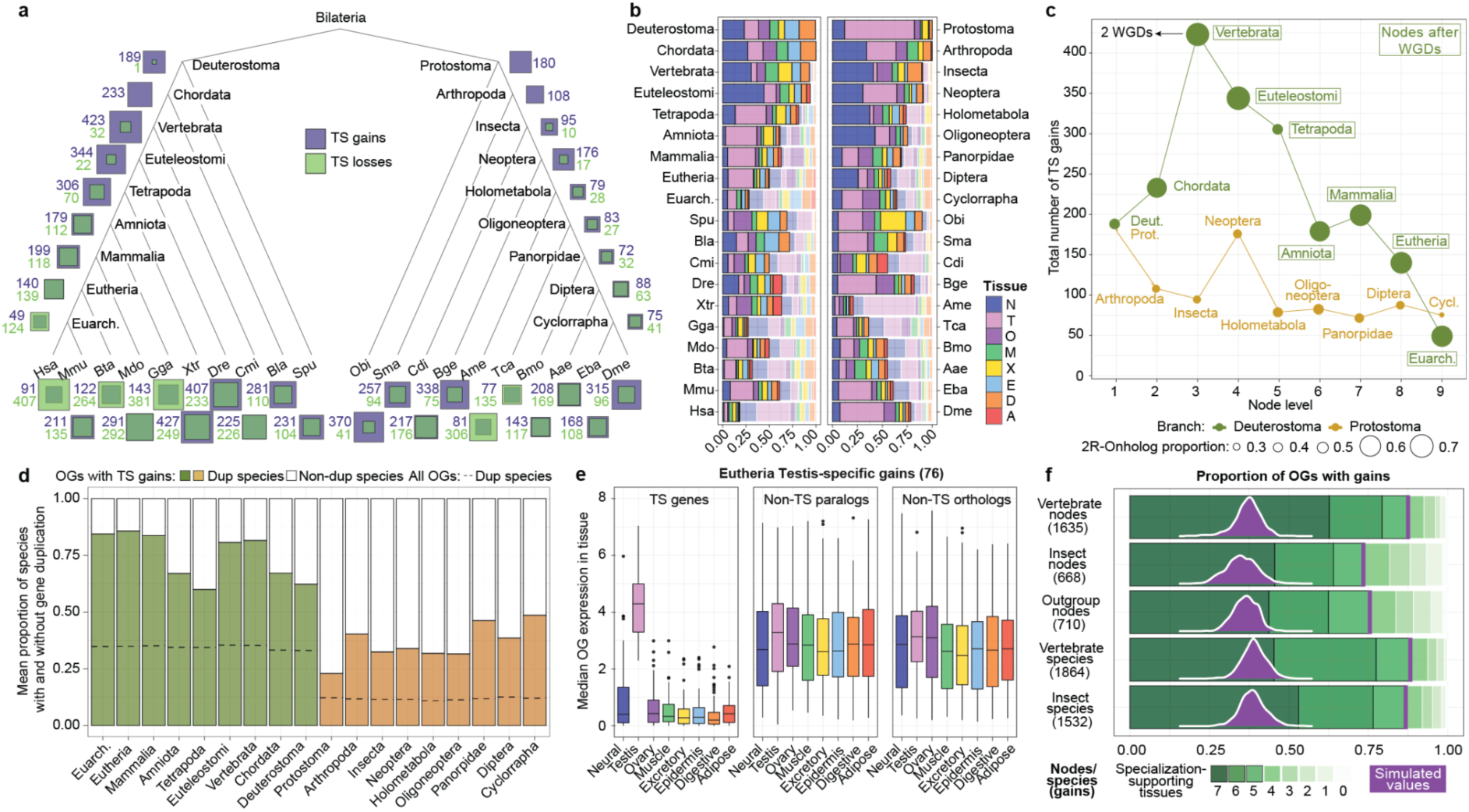
Tissue-specificity gains are associated with gene duplication and specialization. **a.** Total numbers of tissue-specificity gains and losses across all nodes and species. **b.** Relative proportions of tissue-specificity gains and losses across tissues within each node and species. Full/transparent shades of tissue colors represent gains/losses, respectively. **c.** Total number of tissue-specificity gains across nodes on each phylogenetic branch. The size of the dots represents the proportion of orthogroups including 2R-onhologs (i.e., paralogs originated by the two rounds of vertebrate WGDs) in each gain group. **d.** Average proportions of duplicated and non-duplicated species among the species with tissue-specific expression in the orthogroups that gain tissue-specificity in each node. The background line represents the expected proportion based on all bilaterian-conserved orthogroups for the same sets of species (i.e., descendant species for that node). **e.** Median gene expression across tissues for bilaterian-conserved orthogroups with testis-specific gains in Eutheria (76 orthogroups). Left: testis best-TS orthologs in eutherians (3 species). Middle: eutherian non-testis-specific paralogs. Right: testis best-TS orthologs in non-eutherian species (17 species). **f.** For each set of tissue-specificity gains, distribution of the number of tissues (in which the gene is not tissue-specific) where the median expression of the species without tissue-specificity is higher than in the set of species with tissue-specificity (from 0 to 7, “specialization-supporting tissues’’). The purple distribution represents the proportion of gains with specialization-supporting tissues ≥ 5 coming from 100 randomizations of the tissue-specificity labels within the respective best-TS orthogroups (see **Extended Data Fig. 7d,e** for full data). Abbreviations: N: neural, T: testis, O: ovary, M: muscle, X: excretory, E: epidermis, D: digestive, A: adipose, Euarch: Euarchontoglires. Cycl: Cyclorrapha. Deut: Deuterostoma. Prot: Protostoma.

According to our inference methodology, the Vertebrata ancestor presents the highest overall number of inferred tissue-specificity gains among all ancestral nodes (**Fig. 4a**), even if some of the most important gain waves were observed for individual species (e.g., in fruit fly testis and frog ovary, with 198 and 109 gains, respectively; **Fig. 4a** and **Extended Data Fig. 7a**). In this regard, we found that some genomic features were positively correlated with the number of species-specific, tissue-specific gains, but none of them reached statistical significance (**Supplementary Fig. 11a-e**).

Comparison of the proportions of tissue-specificity gains and losses within each node and species shows that testis had the highest turnover, as it presented the greatest proportion of gains and/or losses in 30/39 (77%) of nodes/species (**Fig. 4b**). On the contrary, neural gains are mainly prevalent in the most ancestral nodes on both branches (i.e., Euteleostomi/Neoptera or older), but while they seem to have little impact in later vertebrate evolution, they still dominate the gain landscape in several more recent insect nodes (e.g., Holometabola, Oligoneoptera and Diptera). Finally, as opposed to the most ancestral nodes, the tissue-specific transcriptome of few recent nodes and of the majority of single species is predominantly shaped by losses of tissue-specificity rather than by gains (i.e., we observed an average of 11% of losses on the total number of inferences for Tetrapoda/Holometabola and more ancient nodes, compared to 42% for more recent nodes and single species). While this result is expected, as by definition we infer losses of tissue-specificity only after a tissue-specificity gain has been identified in more ancestral nodes, it is still relevant to highlight the large plasticity of the expression profiles of ancestral genes during recent vertebrate and insect evolution.

We then evaluated enrichments of orthogroups with tissue-specificity gains across gene families, obtaining similar results upon definition of gene families based either on the human (**Supplementary Fig. 12a**) or fruit fly (**Supplementary Fig. 12b**) annotation (See **Methods**). On one hand, we found significant over-representation of tissue-specificity gains for many families of membrane proteins, including several types of transporters and receptors (e.g., ion channels, EGF transporters, G protein-coupled receptors, etc) (**Supplementary Fig. 12a,b**). On the other hand, families of house-keeping genes (e.g., rRNA, histone proteins) or enzymes devoted to basic cellular metabolic processes (e.g., tRNA synthetases, acetyl-transferases) were significantly depleted for tissue-specificity gains, in line with the results reported in **Fig. 3h** and **Supplementary Table 8.**

### Evolution of tissue-specificity is tightly linked to gene duplication and specialization

We then compared the total numbers of tissue-specificity gains between phylogenetically equivalent nodes on the two branches (**Fig. 4c**). The gain signal reached the overall maximum in the Vertebrata ancestor, consistent with a strong impact of the two rounds of WGD at the origin of this group. Strikingly, although this effect progressively decreased, the fraction of gains remained high throughout all subsequent vertebrate nodes, in clear contrast with the relative flat signal observed on the insect side (with the exception of the Neoptera ancestor, in line with a burst of gene duplication events ^37^). Moreover, gains in both the Vertebrata and subsequent nodes, as well as in the species-specific branches, showed a much higher proportion of orthogroups involving paralogs derived from the vertebrate WGDs (2R-ohnologs) compared to phylogenetically equivalent Insecta nodes and species (**Fig. 4c** and **Extended Data Fig. 7b**). Altogether, this suggests the existence of a previously unappreciated prolonged evolutionary impact of vertebrate WGDs on the rewiring of tissue-specific transcriptomes.

Remarkably, the association between the gain of tissue-specificity and gene duplication extended beyond the vertebrate WGDs. For all nodes and extant species, we found that orthogroups with inferred tissue-specificity had a higher proportion of duplicates compared to the corresponding background (**Fig. 4d** and **Extended Data Fig. 7c**). Importantly, while the chances of having a tissue-specific gene are expected to increase with the number of paralogs in an orthogroup, randomizations showed that this effect cannot explain the observed association between gene duplication and tissue-specificity (**Extended Data Fig. 8a**).

Moreover, we found evidence that the acquisition of tissue-specific expression occurred to a large extent through the process of specialization ^38^, where the specialized paralog reduces its expression in most tissues, while (i) the other paralog(s) and (ii) their orthologs in other species conserve the ancestral, broader expression pattern (see scheme in **Extended Data Fig. 8b**). For example, under this model, the Eutheria testis-specific genes are expected to have lower expression across the other tissues (i.e., all tissues but testis) compared to (i) their non testis-specific paralogs in eutherians and (ii) their orthologs in non-eutherians, as readily observed in our dataset (**Fig. 4e**). Both these patterns were consistently observed across all nodes and species (**Extended Data Fig. 8c,d** and **Supplementary Dataset**), which prompted us to systematically evaluate the incidence of specialization events. We used as a metric the number of “specialization-supporting” tissues, corresponding to all those tissues (excluding the tissue with tissue-specificity) where the expression of a given ortholog is lower in the set of species with tissue-specificity compared to those without. We plotted the total number of specialization-supporting tissues (from 0 to 7) for tissue-specificity gains across different groups of nodes and species (**Fig. 4f**), showing that, in all cases, specialization events (i.e., number of specialization-supporting tissues ≥ 5) occurred more extensively than expected by chance (**Fig. 4f** and **Extended Data Fig. 7d,e**).

### Tissue-specificity gains are associated with emergence of unique tissue-related phenotypes

Exploiting the symmetric structure of our phylogeny, we also identified 156 bilaterian-conserved orthogroups that acquired unique but distinct tissue-specificity on the vertebrate and insect sides, thus fulfilling their functional potential in divergent contexts **(Extended Data Fig. 9a**). The most frequent pairs of tissues among which these parallel tissue-specificity gains occurred were neural and testis together with testis and ovary (**Extended Data Fig. 9b**), in agreement with the compartments among which expression shifts are more likely to occur also within vertebrates ^21^. In addition to these divergent/parallel tissue-specificity gains, we also characterized independent convergent acquisitions of the same tissue-specific expression profiles in both the vertebrate and the insect sides (**Fig. 5a**). Such convergent gains were most abundant in testis, probably as a consequence of the faster turnover of tissue-specificity this tissue experienced both in vertebrates and in insects (**Fig. 4b**). One exemplary case is represented by *TESMIN* and *tomb* (**Fig. 5b**). These are paralogs of the ancestral *LIN54/mip120* gene that independently originated in the vertebrate and insect lineages and convergently acquired testis-specific expression in amniotes and the fruit fly, respectively, and whose importance for testis development and function is proven by spermatogenesis disruption upon gene perturbation both in mouse ^39^ and fruit fly ^40^.

**Fig. 5:**
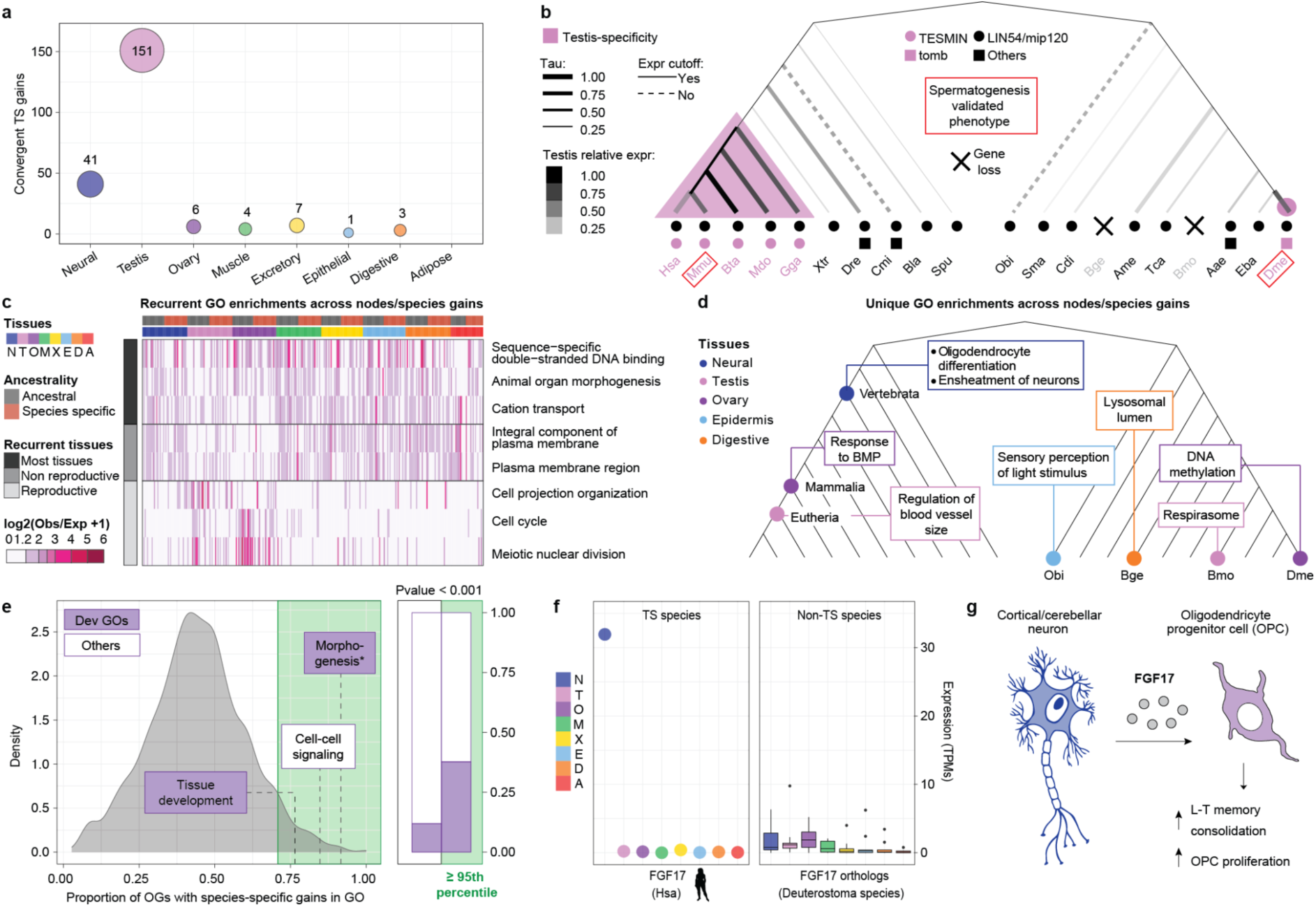
Tissue-specific gains are associated with the emergence of unique phenotypes. **a.** Number of convergent tissue-specificity gains (on the deuterostome and protostome branches) in each tissue. **b.** Example of a convergent testis-specific gain: *TESMIN*/*tomb*. **c.** Heatmap representing GO categories either (i) significantly enriched in the gains of at least 15 nodes/species across all tissues (most/non reproductive labels) or (ii) significantly enriched in the gains of at least 8 nodes/species in one tissue exclusively (reproductive label, which indicates ovary and testis combined). The plotted values (log2(observed/expected+1)) were computed starting from the proportion of gains in each node/species belonging to the tested category (observed) and the proportion of all bilaterian-conserved orthogroups with a functional annotation belonging to the same category (expected). **d.** Examples of GO categories significantly enriched exclusively among the gains of one node/species. **e.** Left: Distribution of the proportion of orthogroups in each GO category with at least one tissue-specific, species-specific gain. The green area represents categories in the 95th percentile or above. Only GO categories including at least ten bilaterian-conserved orthogroups are shown. Right: Proportions of GO terms below or above the 95th percentile representing developmental functions. The reported p-value is computed out of the proportions of developmental functions in the 95th percentile coming from 1000 randomizations of the GO labels (**Extended Data Fig. 9c**). Morphogenesis* stands for “anatomical structure formation involved in morphogenesis”. See Methods for definition of developmental categories. **f.** Expression across tissues for human *FGF17* (left) and its deuterostome orthologs (right). **g.** Schematic summary of FGF17’s function in the brain (based on ^43^). Abbreviations: N: neural, T: testis, O: ovary, M: muscle, X: excretory, E: epidermis, D: digestive, A: adipose, BMP: bone morphogenic protein, TS: tissue-specific, L-T: long-term.

Next, we aimed to functionally characterize the tissue-specificity gains in each node and species (**Supplementary Table 10**). We found that a few gene functions were significantly and repeatedly enriched in multiple nodes/species on both phylogenetic branches (**Fig. 5c**). GO categories such as double-strand DNA binding, cation transmembrane transport and animal organ morphogenesis were over-represented throughout all tissue types, but we also identified a few functions specifically enriched across somatic organ tissues (e.g., plasma membrane region, consistent with the gene family analyses (**Supplementary Fig. 11**) or reproductive ones (mainly related to meiotic division). Moreover, GO enrichments performed with insect-specific GO annotations (see **Methods**) showed repeated enrichments for orthogroups involved in flight and cuticle formation, respectively, in muscle-specific and epidermis-specific gains throughout insect evolution (**Supplementary Table 11,12**). In contrast, each tissue presented categories exclusively enriched across gains in a single node/species (**Fig. 5d** and **Supplementary Table 13)**, several of which could be linked to the concurrent emergence of novel phenotypic traits. For instance, only vertebrate neural-specific gains were significantly enriched in categories related to oligodendrocyte differentiation and ensheathment of neurons (i.e., the myelination of neuronal axons operated by the oligodendrocytes), consistent with the origin of these glial cells in the gnathostome vertebrate ancestor ^41^. In another example, we detected a distinctive enrichment in sensory perception of light stimulus in the octopus’ skin, likely reflecting the unique presence of light-activated chromatophores organs all over the cephalopod’s body surface ^42^.

Finally, we focused on the functions of species-specific, tissue-specific gains, which represent 59% of all our inferred gains. In order to identify functional categories potentially over-represented among these species-specific inferences, we plotted a distribution of GOs based on the proportion of their bilaterian-conserved orthogroups experiencing at least one of such species-specific gains (**Fig. 5e**). Strikingly, the top 5% of this distribution includes cell-cell signaling, tissue development and several other morphogenesis-related or developmental categories, which are significantly over-represented in the upper tail compared to the lower percentiles (**Fig. 5e** and **Extended Data Fig. 9c,d**). One remarkable example of a developmental gene included in these species-specific gains is *FGF17*, which is neural-specific only in human (**Fig. 5f**). *FGF17* is a fibroblast growth factor broadly expressed during the embryonic and postnatal brain development of multiple species, but which was co-opted in the adult brain only in human (**Extended Data Fig. 10**; data from ^19^). Remarkably, a recent study ^43^ showed that the *Fgf17* contained in the cerebrospinal fluid of young mice activates a transcriptional program leading to proliferation of oligodendrocyte progenitors and, when injected into aged mice, slows down brain aging and improves memory functions (**Fig. 5g**). Thus, even if Fgf17’s potential to induce oligodendrocyte proliferation seems to be ancestral, this gene became part of the adult neural-specific transcriptome only during recent human evolution, where it might contribute to the preservation of cognitive abilities in old age.

## Discussion

In this study, we have assembled an extensive dataset of RNA-seq samples spanning twenty bilaterian species and eight tissues, with the goal of tracing the evolution of gene expression in homologous tissues within and between vertebrates and insects. Therefore, in terms of phylogenetic range, our study represents a step forward compared to previous works where a similar framework of tissue transcriptional evolution has been applied ^18–21^, as it extends the investigation range to a large panel of vertebrate and non-vertebrate species, including organisms which diverged ∼700 MYA. Indeed, our principal component analysis highlighted consistent transcriptome variation between vertebrate and non-vertebrate species, as perhaps expected given their distinct genomic features (e.g., larger genomes and gene numbers, as a consequence of the two WGDs events) and evolutionary traits (e.g., longer generation times). This variation manifests into characteristic transcriptomic signatures across tissue types, which are more homogenous in vertebrates and relatively more fast-evolving in the other species. Moreover, we designed our study around a symmetric phylogeny for the vertebrate and insect branches. This allowed us to identify not only ancestral features, but also parallel, convergent and divergent evolutionary trajectories of ancestral genes between and within these two major bilaterian lineages. Using this phylogenetic framework, we performed an analysis of the evolutionary dynamics of tissue-specific expression among ancestral bilaterian genes. Strikingly, we found that nearly half of the ancestral bilaterian gene complement has acquired tissue-specific expression in at least one of the studied species, revealing a surprising plasticity for this transcriptomic trait. We thus investigated the timings and mechanisms behind the pervasive evolution of these tissue-specificity patterns, as well as their functional impact.

Before discussing these aspects, however, we acknowledge that a major limitation of our study is the use of bulk RNA-seq data, which merges the signals originating from the different cell types present in each tissue. This is a relevant issue, given that we are analyzing distantly related species with divergent tissue histologies. Thus, differences in cell type composition might be a confounding factor in our analyses of gene expression dynamics, especially those involving quantitative comparisons among species across the entire tissue panel (e.g., correlations, PCAs, etc.). Notwithstanding, we aimed at minimizing these considerations by explicitly studying tissue-specific patterns of expression, which are largely qualitative in nature (i.e., presence/absence), and thus are more robust to quantitative variations in cell type composition. Indeed, detected changes in tissue-specific transcriptomes can provide information about evolutionary events such as the origin of novel cell types. For instance, we detected enrichment for oligodendrocyte differentiation exclusively in the neural-specific genes acquired in the vertebrate node, concomitantly with the emergence of this cell type ^41^.

With regards to evolutionary timings, our phylogenetic inference revealed that most ancestral genes acquired tissue-specific expression during late bilaterian evolution. Despite this, we found that at least ∼7% of all ancestral orthogroups have been expressed in a tissue-specific manner since the bilaterian LCA, a number that might be higher if we had investigated more slow-evolving species, such as annelids or other lophotrochozoans. Importantly, all the ancestral tissue-specific modules we identified are linked to core functions within each tissue type, even in those tissues more divergent at the histological and cell type level (e.g., digestive system ^44^) or mainly originated by convergent morphological trajectories (e.g., fat-rich tissues ^45^). Moreover, we found that validated phenotypic effects both in mammals and in fly supported their extant, wide physiological relevance. Neural, muscle and reproductive organ transcriptomes present the largest ancestral tissue-specific modules; this suggests they have more distinctive and conserved transcriptomic signatures compared to other bilaterian tissues, likely related to the high complexity and specialization of the main cell types that form them (neurons, myocytes and meiotic cells, respectively).

At the mechanistic level, we showed that tissue-specific gains have a strong association with gene duplication across the entire bilaterian phylogeny, as previously reported for more restricted lineages ^38,46^, and demonstrated that this association is not simply due to multi-copy orthogroups having more chances of including tissue-specific members. Furthermore, we investigated how often evolution of tissue-specificity occurred through specialization, by which the specialized paralog in a given species reduced its expression in the tissues without tissue-specificity compared to the broadly expressed ancestral patterns, which are preserved in the non-tissue-specific paralogs of that species as well as the orthologs in the other species. This pattern had previously been identified for more restricted groups, including paralogs originated from vertebrate ^38^ and salmon ^47^ WGDs, together with gene duplicates specific in the pea aphid ^48^, primates and rodents ^49^; here, we expanded the search space and provided evidence that specialization is associated with tissue-specificity gains throughout the entire bilaterian phylogeny. Another remarkable finding in the context of gene duplication is the seemingly prolonged effect of the vertebrate WGDs on the amount of tissue-specificity gains throughout recent vertebrate evolution. Specifically, the Vertebrata node showed the highest level of tissue-specificity gains in the phylogeny, particularly among paralogs retained from the WGDs (ohnologs), as expected from a direct causal effect of these events. However, subsequent ancestral nodes and extant species within the vertebrate lineage also exhibited higher number of gains, as well as a higher fraction of affected ohnologs, compared to other phylogenetically equivalent non-vertebrate nodes and species. While this could be partly due to loss of tissue-specificity in early branching vertebrate species, we suggest that this pattern reflects the increased likelihood of orthogroups with retained ohnologs in vertebrates to evolve tissue-specificity even millions of years after the WGDs that generated the genetic redundancy. If so, this unexpected finding implies that the evolutionary impact of WGDs on phenotypic diversification may go beyond immediate subsequent effects, providing an additional potential explanation for the lag observed between the timing of WGDs and their purported consequences in multiple lineages ^50–52^.

Finally, we assessed the functional impact of the rewiring of tissue-specific transcriptomes. Almost 60% of our inferred tissue-specificity gains occurred in specific species and were often associated with the emergence of unique phenotypes, highlighting the potential of novel tissue-specific expression patterns to underlie organismal novelties ^53^. For instance, we detected a distinctive enrichment in sensory perception of light stimulus in the octopus’ skin, consistent with the unique presence of light-activated chromatophores organs all over the cephalopod’s body surface ^42^. We also uncovered a significant tendency for developmental genes to retain adult tissue-specific expression in a species-specific manner. Cases like *FGF17* in humans that we reported here point to a potential widespread, functional co-option of ancestral developmental genes within distinct tissue-specific transcriptomes throughout the most recent bilaterian evolution. Future research should elucidate the functional significance of the adult tissue-specific expression of these developmental genes as well as of the myriad of other tissue-specific genes we identified, and how they ultimately contribute to animal evolution.

## Methods

### Genome annotation and sequence files

For eight species, we downloaded the GTF and genome FASTA files from Ensembl [https://www.ensembl.org/], selecting the following assemblies and versions: human (*Homo sapiens, Hsa*: hg38, v88), mouse (*Mus musculus, Mmu:* mm10, v88), cow (*Bos taurus, Bta*: bosTau9, v99), opossum (*Monodelphis domestica, Mdo*: monDom5, v86), chicken (*Gallus gallus, Gga*: galGal6, v99), tropical clawed frog (*Xenopus tropicalis, Xtr*: XenTro9, v101), zebrafish (*Danio rerio, Dre*: danRer10, v80), elephant shark (*Callorhinchus milii, Cmi:* 6.1.3, v99). For other eight species, we downloaded GTF and genome FASTA files from Ensembl Metazoa [https://metazoa.ensembl.org/]: sea urchin (*Strongylocentrotus purpuratus:* Spu: Spur_5.0 v51), fruit fly (*Drosophila melanogaster, Dme*: dm6, v26), yellow fever mosquito (*Aedes aegypti, Aae*: AaegL5, v46), domestic silk moth (*Bombyx mori, Bmo*: ASM15162v1, v45), red flour beetle (*Tribolium castaneum, Tca*: Tcas5.2, v45), honey bee (*Apis mellifera, Ame*: Amel_4.5, v35), centipede (*Strigamia maritima*, Sma: Smar1, v26), California two-spot octopus (*Octopus bimaculoides, Obi*: ASM119413v2). For other three species, we used the GTF and genome FASTA files from the relative publications: amphioxus (*Branchiostoma lanceolatum*, Bla: ^54^), marmalade hoverfly (*Episyrphus balteatus*, Eba: ^55^), mayfly (*Cloeon dipterum*, Cdi: ^12^). For the cockroach (*Blattella germanica*, Bge) we downloaded the GFF3 (blager_OGSv1.2.1.gff3) and the genome FASTA (GCA_000762945.2_Bger_2.0_genomic.fna.gz) from https://i5k.nal.usda.gov/content/data-downloads. We then converted the GFF3 to GTF with gffread and we modified it to match the Ensembl format using custom scripts. Finally, for Bmo and Bge, we enriched the annotations using the RNA-seq data we generated as described in the **Supplementary Methods**.

### Gene orthology calls

We used *Broccoli (v1.2*) ^56^ to infer gene orthogroups among all protein-coding genes from the 20 species. In order to avoid redundant gene homology calls, we selected one representative protein isoform for each gene in each species (i.e., the isoform with the longest coding sequence). See **Extended Data Fig. 1** and **Supplementary Table 2** for gene orthogroup statistics and **Supplementary Dataset** for gene orthogroup files.

### Genome annotation refinements

We corrected broken genes (a single gene annotated as two or more separated entities) and chimeric genes (independent genes annotated as a fused element) from almost all genome annotations (excluding human, mouse and fruit fly), since they can respectively result in erroneous gene duplication inferences and incomplete ortholog detection. For this purpose, we used the information provided by the gene orthology call as well as pairwise alignments, as described in detail in the **Supplementary Methods**. We finally ran *BUSCO* ^57^ (options *-m proteins-l metazoa_odb10*) to assess the quality of these refined genome annotations using as proxy 954 metazoan single-copy orthologs (**Extended Data Fig. 1d**).

### Gene Ontology (GO) annotations and transfers

Comparative analyses relying on functional GO annotations risk being biased by the different GO annotation qualities existing between species. In order to avoid such biases, we generated a unified annotation for each of the species starting from the assumption that orthologous genes likely share functional properties. First, we built comprehensive GO annotations both for human and fruit fly. We downloaded the GO annotations (GeneID-GO correspondence) from *Ensembl (v106)* ^58^ for the two species, and we combined the human annotations with those from *clueGO v2.5.5 level 5* ^59^. Then, to build a human-based GO annotation file, we assigned the GO annotations of each human gene to all the genes from the other species belonging to the same orthogroup whenever the number of human genes within the orthogroup with that GO annotation was ≥ 1/4 of the total human genes in that orthogroup. A similar procedure was followed to build a fruit fly-based GO annotation file derived from the fruit fly GO annotation. Then, we selected only the GO categories with a number of genes included between 3 and 1,500 for the human-based annotation and 3 and 500 for the fruit fly one. The annotation files resulting from these “ontology transfers” (see **Supplementary Dataset**) were used for downstream GO analyses.

To get additional insights into potentially novel functions that emerged in vertebrate or insect evolution, we also generated GO transfers based on GO annotations unique in either vertebrates or insects. Apart from the human and fruit fly GO annotations (see above), we downloaded annotations for all the vertebrate and insect species in our dataset available in Ensembl: mouse, cow, chicken, frog, shark (Ensembl*, v106*), opossum (Ensembl*, v97*), zebrafish (Ensembl*, v80*), yellow fever mosquito, (Ensembl Metazoa*, v57*), domestic silk moth, red flour beetle (Ensembl Metazoa*, v45*) and honey bee (Ensembl Metazoa*, v37*). We combined the annotations of vertebrate and insect species separately, and selected all the genes belonging to GO categories exclusively annotated in either of the two clades. We then used these vertebrate and insect-specific annotations to generate GO transfers with the same procedure described above, filtering for only those GO categories represented in at least two bilaterian-conserved orthogroups. The resulting GO transfer files are available in the **Supplementary Dataset**.

## RNA-seq sample dataset

We downloaded in total 1,136 individual RNA-seq samples across 18 species. All downloaded samples and relative information can be found in **Supplementary Table 3**. Moreover, we generated 89 RNA-seq samples for 15 species covering different tissues that were missing in public resources. See **Supplementary Table 3** for more details (all the samples generated for this project report “in_house” in the Sample_origin field). All these samples were dissected from adult animals and the RNA extracted using the most suited protocol for the organism and tissue, namely TRIzol™ Reagent (Thermo Fisher) or Qiagen RNeasy kit (QIAGEN) (see GEO series GSE205498 for more details on sample extraction and processing protocols). These RNA samples were used to construct standard Illumina RNA-seq libraries at the CRG Genomics Unit, and an average of ∼78 million 125-nucleotide paired-end reads were generated for each of them in a HiSeq2500. In the case of octopus, the sequencing was performed at the University of Chicago, with NovaSeq. In total, we generated ∼7.6 billion individual reads. FastQC reports for all in-house generated samples are available in the **Supplementary Dataset**. Extra metadata information for both in-house generated and publicly available samples is provided in **Supplementary Table 3**.

## RNA-seq quantification

We quantified expression using *Kallisto quant* ^60^, setting parameter *-b 100--single-l 190-s 20* for single end RNA-seq samples and *-b 100* for paired-end RNA-seq samples (with-b=number of bootstrap samples;-l=estimated average fragment length;-s=estimated standard deviation of fragment length). *Kallisto* logs for all samples are provided in the **Supplementary Dataset**. For each species, we quantified gene expression for each sample by summing the raw counts of all its corresponding annotated transcripts. We next normalized the expression with *DESeq2* ^61^ and used the effective lengths returned by Kallisto to compute the Transcript Per Million (TPMs). For all analyses, log2(TPM+1) was used as the final expression measure for each sample.

## Meta-samples and tissue expression measures

When multiple datasets were available for a given tissue and species, we grouped the RNA-seq samples into a maximum of three meta-samples. This was done to: (i) increase read depth per meta-sample, (ii) dilute potential batch effects from publicly available samples, (iii) facilitate downstream analyses and comparisons by having a comparable number of replicates across tissues and species. Meta-sample groups are detailed in the column “Metasample” in **Supplementary Table 3**. In particular, we followed the approach that we previously described for human, mouse, cow, zebrafish and fruit flyin *VastDB* [https://vastdb.crg.eu/] ^62^, where samples from comparable experiments based on clustering approaches are pooled. For this study, we first computed the median expression across all the samples included in each meta-sample, which we used as its representative measure. Then, we calculated the average of these measures across all the meta-samples belonging to each tissue, generating representative expression values at the tissue level. The same expression quantification procedure was adopted for both in-house generated and publicly available samples. Importantly, the use of alternative combinations of RNA-seq samples to define meta-samples, computed either after averaging the expression of all available samples for each tissue or after randomizing the samples of each tissue across the relative meta-samples, yielded similar results (see below).

## Definition of best-ancestral orthogroups

Most comparative analyses need matrices with a single gene per species as input, which often restricts the studies to single 1-to-1 orthologs, which are known to have different evolutionary and expression biases ^38^. To expand our gene sets beyond standard 1-to-1 orthologies, we defined a best representative ortholog per species in each of the 7,178 bilaterian-conserved orthogroup (see scheme in **Extended Data Fig. 2a**), aiming at selecting those orthologs in each species that most likely conserved the ancestral protein sequence and expression profile (i.e., best-ancestral). We reasoned that such orthologs would be the most suitable to reconstruct ancestral transcriptional patterns, even if we acknowledge that the least divergent paralog is not always the one conserving the original expression profile ^63^. For this purpose, we first calculated the pairwise protein sequence similarity for all pairs of genes from different species within an orthogroup based on BLOSUM62, using *mafft* ^64^ with default parameters and comparing the best representative isoform per gene. Next, for any species with multiple paralogs in a given orthogroup, we selected as best-ancestral the gene with the highest average sequence similarity (with respect to the genes from all other species in the orthogroup) if this similarity value was at least 0.2 higher than the sequence similarity of all the other paralogs from the same species. If this requirement was not fulfilled, we discarded the genes with ≥0.2 average sequence similarity difference from the gene with the top average sequence similarity, and we selected as best-ancestral among the remaining genes the one with the highest expression similarity across tissues (or a random one in case of equal expression similarity). This expression similarity was defined as the average of all pairwise Pearson’s expression correlations across tissues between the target gene and each gene from all other species in the orthogroup (i.e., considering all genes where there are multiple copies), where expression in each tissue was defined as described above. Finally, it should be noted that, for some analyses, only the 2,436 best-ancestral orthogroups conserved in all species were considered.

To test if, as we hypothesized, the best-ancestral orthologs were the most suitable to reconstruct ancestral transcriptomic patterns, we compared them to alternative strategies for paralog selection or orthogroup expression quantification. We created three control sets, considering as expression measure for each species in each orthogroup (i) the average expression among all its paralogs, (ii) the summed expression among all its paralogs and (iii) the expression of a randomly selected paralog. For all the control sets, we generated z-scored expression matrices as described in “Best-ancestral orthogroups normalized and z-scored expression matrices”, and from these matrices, we computed all intra-tissue, species-pairwise Pearson’s expression correlations across orthogroups. Comparisons of these correlations between the best-ancestral orthogroups and the control sets are available in **Extended Data Fig. 2b**.

## Best-ancestral orthogroups normalized and z-scored expression matrices

For the normalized expression matrix, we first created a matrix with the expression values of the best-ancestral orthologs across the relative orthogroup (rows) and species’ meta-samples (columns). We then quantile normalized the initial matrix using *limma* in R ^65^ only the 2,436 best-ancestral orthogroups with at least one ortholog in each species.

For the z-scored expression matrix, we first created a matrix with the expression values of the best-ancestral orthologs across the relative orthogroup (rows) and species’ tissues (columns). We quantile normalized this initial matrix using *limma* in R ^65^ only the 2,436 best-ancestral orthogroups with at least one ortholog in each species. Finally, we applied a z-score transformation across tissues of the same species within each orthogroup, which should minimize the inter-species variability.

## Principal component analysis (PCA) and clustering analysis

To investigate the interrelation among our meta-samples, we performed a PCA on the best-ancestral orthogroups normalized expression matrix (see previous section) using the *prcomp* function in R (center=TRUE, scale=TRUE). To assess the biological nature of each principal component, we performed one-sided ANOVA tests between species and tissue groups employing the *aov* function in R (**Extended Data Fig. 3c**; shown p-values were Bonferroni corrected). The heatmap in **Extended Data Fig. 3d** was generated by the *pheatmap* R package with *ward.D2* clustering method on the best-ancestral orthogroups z-scored expression matrix (see previous section).

In order to prove the robustness of the pattern observed in the PCA in **Fig. 1b**, we also tested if any batch effects were affecting the clustering of vertebrate vs non-vertebrate meta-samples along the first principal component. First, we selected meta-sample related variables (sequencing strategy, sample origin, sequencing technology, read depth, read length and total number of samples) from the metadata information in **Supplementary Table 3**. Then, we tested if any of them was introducing a potential batch effect by assessing if the same separation between vertebrate and non-vertebrate meta-samples was observed within batches. For the categorical variables, we selected all the meta-samples from the most abundant category: paired end sequencing for the sequencing strategy, publicly available samples for the sample origin, and Illumina HiSeq 2500 for the sequencing technology (**Supplementary Fig. 2a-c**, left panels). For the continuous variables, we selected only the meta-samples falling within a restricted interval around the mean or median variable value: mean read length between 140 and 160, total number of reads between 80 and 100 millions, and total number of samples equal to 1 (**Supplementary Fig. 2d-f**, left panels). We then repeated the PCA with each meta-sample selection (**Supplementary Fig. 2a-f**, right panels). Moreover, In order to show that our definition of best-ancestral orthogroups was not responsible for the observed pattern, we compared it to alternative strategies for paralog selection or orthogroup expression quantification. We created three control sets, considering as expression measure for each species in each orthogroup (i) the average expression among all its paralogs, (ii) the summed expression among all its paralogs and (iii) the expression of a randomly selected paralog (see ‘Definition of best-ancestral orthogroups“). For all these sets, we generated normalized expression matrices as described in “Best-ancestral orthogroups normalized and z-scored expression matrices”, and repeated the PCA (**Supplementary Fig. 2g-j**).

## Vertebrate and insect PCAs

We first performed two new *Broccoli (v1.2*) ^56^ runs to derive separate orthologies for the vertebrate and insect species. From each orthology, we selected only the single-copy orthogroups conserved across all species (5,398 for vertebrates and 1,766 for insects). We then generated normalized gene expression matrices for these orthogroups following the procedure described for the best-ancestral orthogroups, and performed a PCA for each lineage separately (**Supplementary Fig. 3a,c**). Significant separation of species groups across principal components (**Supplementary Fig. 3d**) were computed through one-sided ANOVA tests between species groups employing the *aov* function in R (see above: “Principal component analysis (PCA) and clustering analysis”).

## Definition of ancestral bilaterian tissue-specific modules

As summarized in **Fig. 2a**, we first performed a sparse partial least square discriminant analysis (sPLS-DA) with the *splsda* function from the *mixOmics* package ^66^ in R, using as input the best-ancestral orthogroups normalized expression matrix (but where all bilaterian-conserved orthogroups are included). We specifically compared all tissue groups versus each other, selecting the optimal number of components and loadings per component by running the *tune.splsda* function on the same expression table with the following parameters: *ncomp = 10, validation = ‘Mfold’, folds = 4, dist = ‘max.dist’, measure = “BER”, test.keepX = c(1:10, seq(20, 300, 10)), nrepeat=10*. Since each of the resulting components specifically separated the meta-samples of each tissue group (Extended Data Fig. 4a-f), we used the corresponding loadings (which represent orthogroups with the most distinctive expression profiles in the isolated tissue compared to the others) to define the respective ancestral bilaterian tissue-specific modules. Importantly, contrary to a PCA, the proportion of variance explained by consecutive components does not necessarily decrease, as the aim is not to maximize the variance. As an extra filter, we further selected only those best-ancestral orthogroups that had the highest median expression in the isolated tissue both among vertebrates and insects. In order to be able to pool tissue expression values across species, we considered the z-scored expression matrix described above (see “Best-ancestral orthogroups normalized and z-scored expression matrices”), but where all bilaterian-ancestral orthogroups are included. The values plotted in **Fig. 2c,d** and **Extended Data Fig. 4g-l** correspond to the median of these expression measures among all vertebrates, all insects or all outgroups (i.e., only three values instead of 20 are plotted per orthogroup and tissue).

## Characterization of ancestral bilaterian tissue-specific modules

GO enrichment analyses were performed with the *gprofiler2* ^67^ R package, using either the human or the fruit fly ontology transfers as GO annotation and all bilaterian-conserved, best-ancestral orthogroups as background. All p-values were FDR corrected. Results obtained with both GO annotations are provided in **Supplementary Table 6** (human) and **Supplementary Table 7** (fruit fly), but only GO enrichments from the human annotations were discussed in the relative Results section and represented in **Fig. 2e,f,i**. For the representation of GO networks of significantly enriched categories (adjusted p-value ≤ 0.05) in **Fig. 2i**, only significant categories containing at least 5 genes in the tested set were considered. The networks were obtained from *Revigo* (http://revigo.irb.hr/) ^68^, selecting large output lists (90% of the input list; option 0.9) for all modules except the neural-differential (for which 0.4 [40%] was selected). To characterize the phenotypic impact of these genes, we downloaded all validated gene-phenotype associations from Ensembl (^58^; v105) for human and mouse and from FlyBase ^69^ updated in January 2020. Neural phenotypes were defined as anything matching “neuro”, “behavior”, “brain”, “glia”, “CNS” (case insensitive), while testis phenotypes (which we reasoned should include also broader reproduction-related phenotypes) were defined as anything matching “sperm”, “infert”, “sterile”, “testis” (case insensitive). Orthogroups with positive matches in either species were considered for the plots shown in **Fig. 2g,h**. In this analysis, no distinctions were made between genes lacking a neural/testis phenotype and genes without phenotypic characterization. Neural and testis phenotypes associated with the respective ancestral tissue-specific module are reported in **Supplementary Table 5**, while all phenotypic associations mapped to the respective bilaterian-conserved orthogroup are available in the **Supplementary Dataset**. Finally, the PCAs in **Supplementary Fig. 4a,b** were performed on the best-ancestral orthogroups normalized expression matrix (see above) but after filtering either for all the orthogroups belonging to the ancestral bilaterian tissue-specific modules (**Supplementary Fig. 4a**) or for a matched number of orthogroups from each tissue’s modules (maximum of 20 random orthogroups per module; **Supplementary Fig. 4b**).

## Enrichment of TF binding motifs in ancestral bilaterian tissue-specific modules

To investigate which TFs might be behind ancestral bilaterian tissue-specific modules, we first built a database of Positional Weight Matrices (PWMs) combining motifs from 56 vertebrate and non-vertebrate species, for a total of 8016 original motifs with direct evidence in CIS-BP ^70^. We clustered similar motifs into consensus motifs by running *gimme_cluster* ^71^ with parameter *-t 0.9999* as described in ^38^, obtaining a final set of 1406 PWMs of length ≥ 5. We then mapped each of these consensus motifs to the respective bilaterian-conserved orthogroup(s) based on the genes (in all species) from which the original motifs in the group were derived. The database containing the orthogroup-motif cluster associations is provided in the **Supplementary Dataset**.

We then checked if the TFs included in the ancestral bilaterian tissue-specific modules presented binding sites enriched in the regulatory regions of the other genes in the same modules (results shown in **Fig. 2j** and **Supplementary Fig. 4c**). For this, we considered as regulatory region the 3kb upstream of each gene’s annotated transcription start site, or the intergenic distance from the closest upstream gene (last poly-adenilation site) if this was lower than 3kb. For all bilaterian-conserved genes in each species, we then calculated the di-nucleotide frequency with *rsat oligo analysis-2str-noov* ^72^. We scanned all the sequences in each species, using the di-nucleotide frequencies as background and running *rsat matrix-scan* with the following parameter: *-quick-pseudo 1-decimals 1-2str-log_pseudo 0.01-uth pval 0.01-n score*. From the returned hits, we only selected the highly confident matches in each sequence (p-value ≤ 1e-05). Then, for all ancestral tissue-specific TFs, we performed two tests on the corresponding module, using as background all other bilaterian-conserved genes: (i) a Fisher’s exact test, to check if the proportion of sequences with at least one hit in the tested module was significantly higher compared to the background proportion; and (ii) a negative binomial regression test, to check if the number of hits per sequence was increased among the genes in the tested module compared to the background. We performed each test three times, selecting in turns all, only vertebrate or only insect species. In **Fig. 2j** we represent all TFs which have: (i) either p-value ≤ 0.05 and positive beta in the regression test or p-value ≤ 0.05 in the Fisher’s exact test performed on all species; and (ii) p-value ≤ 0.05 for both vertebrates and insect species separately either in the regression or Fisher’s exact test. Finally, in order to assess whether the observed enrichments were driven by a shared signal across the entire phylogeny or derived from only a few species, we repeated the same tests separately in each species, and plotted the observed vs. expected values for each TF and tissue set (**Supplementary Fig. 4c**).

## Tissue-specificity calls

To perform the tissue-specificity calls, we first computed the Tau ^36^ for all genes separately in each species. Tau is a measure of tissue-specificity ranging from 0 (ubiquitous genes) to 1 (highly tissue-specific genes). For each species, we employed as input a quantile-normalized expression matrix of log2(TPMs+1) values averaged by tissue (i.e., one value per tissue). We defined as tissue-specific in each species all genes with Tau ≥ 0.75 and maximum expression ≥ log2(5). The Tau threshold was chosen by looking at the general Tau distributions across species (**Fig. 3a** and **Supplementary Fig. 5**), many of which show a bimodal trend where a Tau cutoff of 0.75 would select the majority of the upper tail (i.e., highly tissue-specific genes); moreover, it was similar to what implemented as a tissue-specificity threshold in previous publications ^73–75^. To associate these tissue-specific genes with one or two tissues (“Associated tissue(s)” in **Fig. 3b**, **Supplementary Fig. 6** and **Extended Data Fig. 2c**), we evaluated the expression proportion per tissue (tissue_expr / all_tissue_expr), where “tissue_expr” is the average normalized log2(TPMs+1) expression of the gene in the target tissue and “all_tissue_expr” the sum of the average normalized log2(TPMs+1) expression values across all tissues. Specifically, we applied the following steps for each gene in each species (**Extended Data Fig. 2c**): (i) if the difference in expression proportion between two most-highly expressed tissues was ≥ 0.10 and their ratio ≥ 1.7, we associated the gene only with the top tissue. (ii) If the above conditions are not fulfilled, but the difference in expression proportion between the second and third most highly-expressed tissues was ≥ 0.15, we associated the gene with the two top tissues (double-tissue-specificity). (iii) Else, the gene was not considered as tissue-specific and not associated with any tissue. In addition, for the gain/loss inferences (see next section), we more loosely defined the “Top tissue(s)”, corresponding to the “Associated tissue(s)”, when available, or simply to the two tissues with the highest expression (**Extended Data Fig. 2c**, last example).

## Definition of best-TS orthogroups

We generated eight sets (one per tissue) of one-to-one “best-TS” orthogroups, selecting the orthologs in each species that showed the most prominent tissue-specific expression profile in the query tissue. In particular, we selected the paralog with the strongest association with the query tissue according to the following prioritization (**Extended Data Fig. 2d**): (i) a paralog called as tissue-specific in that tissue as defined above; if there were multiple tissue-specific paralogs, we selected the one with the highest Tau. (ii) Else, a paralog in which the target tissue is in the Top tissue(s), as defined above; if there were multiple such paralogs, we selected the one with the highest Tau, giving priority to those passing the expression cutoff (max expression ≥ log2(5). (iii) Else, the paralog with the highest Tau, giving priority to those passing the expression cutoff.

## Phylogenetic inference of tissue-specificity gains

We performed the phylogenetic inferences of tissue-specificity gains and losses for each tissue separately, considering all the orthogroups presenting at least one tissue-specific call in that tissue (see “tissue-specificity calls”). We implemented two subsequent ad-hoc, parsimony-based inference approaches independently for each major branch (deuterostome and protostome), which we developed due to the limitations of other inference methods with respect to our dataset and our specific scientific aim (see **Supplementary Discussion**). First, we applied a “strict approach”, inferring a maximum of one tissue-specificity gain for each major branch. Here, we inferred a gain in a node if (**Extended Data Fig. 5a**, left): (i) the first-branching species in the node was tissue-specific in the query tissue (as defined in the previous section); (ii) at least 50% of the node’s descendant species with an ortholog had Tau ≥ 0.60 and were associated with the query tissue; and (iii) none of the outgroup species to that node on the same branch that passed the expression cutoff had Tau ≥ 0.60 and were associated with the query tissue. Exceptionally, in the case of the most internal nodes (i.e. Euarchontoglires: human and mouse, Cyclorrhapha: fruit fly and hoverfly) we required Tau ≥ 0.6 and association with the query tissue in both species, and a tissue-specific call in that tissue for at least one of them.

Second, for all the orthogroups that could not be classified with the first strict approach for a given branch, we inferred gains with less stringent requirements (“relaxed approach”; **Extended Data Fig. 5a**, right). Here, we inferred gains in the last common ancestor of all species with Tau ≥ 0.60 that are associated with the query tissue as long as at least one tissue-specific gene is present. However, the relaxed approach inferred multiple gains on each branch if the minimum distance between two species or nodes respecting those tissue-specificity cutoffs was higher than 3 nodes (e.g. in human and in chicken, or in Eutheria and in zebrafish). Also, if no inference of gain in an ancestral node could be done by either approach, tissue-specific genes (as defined in the previous section) were considered species-specific gains. Finally, from the combined output of both approaches, we inferred an ancestral bilaterian (or earlier) tissue-specificity whenever a “gain” was identified in both Deuterostoma/Chordata/Vertebrata and Protostoma/Arthropoda/Insecta with either strict or relaxed criteria (**Extended Data Fig. 5b**; “merged” label in **Supplementary Table 9**). As an exception, since shark testis samples showed poor correlation with other testis samples, we also inferred ancestral bilaterian tissue-specificity for testis in case of gain inferences in Euteleostomi and Protostoma.

## Phylogenetic inference of tissue-specificity losses

We then inferred tissue-specificity losses exclusively starting from the nodes in which gains were inferred for each tissue (**Extended Data Fig. 5c**). In case of ancestral bilaterian tissue-specificity, the inferences were conducted separately on the deuterostome and protostome branches. We considered as potential losses all species (internal to the node with the inferred gain) where either: (i) Tau ≤ 0.45; or (ii) the query tissue is not among the top tissue(s), as defined above (**Extended Data Fig. 2c**); or (iii) the difference in expression proportions between the query tissue and the third highest tissue is ≤ 0.1. Then, starting from the innermost species with a potential loss, if there were two or more consecutive such species, we inferred a loss in the node corresponding to their LCA and a novel gain in the node of the LCA of their consecutive inner species if: (i) all these species are tissue-specific as described above; (ii) the ancestral loss is separated by at least one node from the most ancestral gain, and (iii) the total number of these new inferences (including single losses in all the species excluded from the ancestral loss inference) is lower than the number of original inferences (i.e. independent losses for each potential loss species). Otherwise, separated losses for each single species are inferred.

## Validation of the inferred tissue-specificity gains

We performed two kinds of validations for our inferred tissue-specificity gains, based on the comparison with the inferences derived (1) upon Tau quantification based on alternative combinations of the initial samples or (2) by an independent methodology.

For the first validation approach, we first computed Tau values across species (i) after averaging the expression of all available samples for each tissue and (ii) after randomizing the samples of each tissue across the relative meta-samples. Of note, these “alternative” Taus could only be computed for the 13/20 species for which at least one of the tissues contained meta-samples composed by more than one sample. We then repeated the tissue-specificity call and the inferences of tissue-specificity gains and losses using these alternative Tau quantifications. Comparisons of the original Taus and inferences with the ones returned by these approaches are available in **Supplementary Figs. 8,9**.

For the second validation approach, in each tissue separately, we selected all the best-TS orthogroups with inferred tissue-specificity gains which: (1) presented orthologs in all species, (2) for which all orthologs passed the same expression cut-off imposed in our tissue-specificity call (i.e., maximum expr ≥ 5 TPMs), and (3) were not inferred to be present before the last common ancestor of Chordata and/or Arthropoda. In particular, this last filtering was applied so that there would be at least one species with and without tissue-specificity, for which the existence of putatively distinct expression optima could be tested. Adipose-specific gains were excluded from this validation, as both branches present species with missing samples and consequently lack gene expression quantification in at least one of them. We then used the *OUwie* function from the homonymous R package ^76^ to fit a double-optima (OUM) and a single-optimum (OU1) OU models to the expression proportions in the query tissue. The double-optima OU model specifically assumed differences between the species with and without tissue-specificity, which have to be specified a priori, and where the latter group also included all species with inferred losses. To determine which model better fit the data, we compared the Akaike information criterion (AIC) weights ^77^; in case of equal AIC weights between the two models, we declared that no optimal model was found (**Extended Data Fig. 6**, left panels). Then, for those cases where the double-OU model was the best fit, we checked if expression in the query tissue was higher for the species with tissue-specificity compared to the species without, expecting positive deltas for all cases representing supported tissue-specificity gains (**Extended Data Fig. 6**, right panels). Finally, we generated two control groups for each of the test tissue sets, selecting a matching number of best-ancestral orthogroups (either starting from all orthogroups or only from those orthogroups without tissue-specific gains) to which we randomly assigned the tissue-specificity gain labels from the relative test set. The same analyses were performed as for the test sets, and results shown side by side on **Extended Data Fig. 6**.

## Characterization of non-tissue-specific orthogroups

The GO enrichments for the non-tissue-specific orthogroups shown in **Fig. 3h** and **Supplementary Table 8** were performed as described in “Characterization of ancestral bilaterian tissue-specific modules”. The duplication status of these orthogroups compared to the tissue-specific and all orthogroups was evaluated by counting the number of species in each of them that presented at least two paralogs (also shown in **Fig. 3h**). The difference between the TS and non-TS orthogroups in terms of duplications was assessed with Fisher’s exact test (*fisher.test* function in R, with *simulate.p.value = TRUE*).

## Enrichments of tissue-specificity gains and losses across gene families

We defined gene families based on PFAM clans, which represent groups of related protein domains ^78^, and with an approach analogous to the GO transfers. First, we collected the PFAM clan annotation (release 36.0), containing the association between PFAM clans and relative PFAM protein domains. Then, we mapped the human proteome to the relative PFAM domains through the provided *pfam_scan.pl* script (parameters: *-e_dom* 0.1 *-e_seq* 0.1; release 27.0), generating a human PFAM annotation (gene-PFAM domain correspondences). We combined these annotations and assigned each human gene to the relative PFAM clans, preserving the PFAM domain annotation in case of domains that did not belong to any clan. We then associated the bilaterian-conserved orthogroups with the relative PFAM clans using the approach implemented for the GO transfers (see above), and we selected only those PFAM clans (gene families hereafter) containing at least 20 orthogroups. We repeated the same procedure with fruit fly PFAM annotation, and performed all downstream analyses with both sets of gene families (**Supplementary Fig. 12a,b**).

First, we evaluated enrichments/depletions of orthogroups with any kind of tissue-specificity gains across the selected gene families. Enrichments were computed with a binomial test (*binom.test* function from the R *stats* package) with the following parameters: *x*=number of orthogroups in the gene family with tissue-specificity gains; *n*=total number of orthogroups in the gene family; *p*=proportion of orthogroups with tissue-specificity gains over all bilaterian-conserved orthogroups; *alternative*=two sided.

Second, we evaluated enrichments/depletions of orthogroups undergoing tissue-specificity gains in multiple tissues (multi-TS) across the same gene families. The binomial test was run with the following parameters: *x*=number of orthogroups in the gene family with multi-TS gains; *n*=total number of orthogroups in the gene family with tissue-specificity gains; *p*=proportion of orthogroups with multi-TS gains over all bilaterian-conserved orthogroups with tissue-specificity gains; *alternative*=two sided.

## Duplication and specialization patterns of tissue-specificity gains

Each orthogroup’s duplicated proportion was defined as the number of species with at least two paralogs over the total number of considered species (which depends on the tested node). The mean duplicated proportion for the orthogroups with gains in each node compared to the relative background (i.e. all orthogroups in that node) is shown in **Fig. 4d**. The proportion of orthogroups with gains including 2R-onhologs (**Fig. 4c**) was based on the list of 2R-ohnologs provided by ^79^. The ten randomized bilaterian-conserved gene orthogroups used in **Extended Data Fig. 8a** were obtained by shuffling genes within each species while preserving the original paralogy structures (i.e., each randomized orthogroup conserved the original number of paralogs from each species, but the actual orthologous genes no longer corresponded across species).

We then checked how each tissue-specific gain fitted the specialization hypothesis. We started from the same expression matrices used for the tissue-specificity call (see above), comparing the median expression in each tissue between species with tissue-specificity and species without tissue-specificity (including species with inferred tissue-specificity losses). For each gain, we counted for how many tissues (excluding the tissue with tissue-specificity) this median expression was higher in the species without tissue-specificity (specialization-supporting tissues, ranging 0-7; relative proportions across nodes and species in **Fig. 4f** and **Extended Data Fig. 7d,e**). For the gains in each node and species, we performed 100 randomizations of the tissue-specificity labels among all species in the relative orthogroup. For each of these randomization rounds, we counted the proportion of gains in which the number of specialization-supporting tissues was ≥ 5. We plotted in purple the distributions of these proportions for all randomizations overlaying the relative observed distributions in **Extended Data Fig. 7d,e** or their collapsed distributions in **Fig. 4f**.

## Functional characterization of tissue-specificity gains

Parallel and convergent gains of tissue-specificity (**Extended Data Fig. 9a** and **Fig. 5a**) were evaluated exclusively among those best-TS orthogroups that present tissue-specificity gains in only one tissue on each of the main branches (deuterostome or protostome). The GO enrichment analysis on the orthogroups with gains in each node/species reported in **Fig. 5c,d** and **Supplementary Tables 10,13** were performed as described in “Characterization of ancestral bilaterian tissue-specific modules” and using the GO transfers derived from the human annotation. The same enrichments were also repeated using the vertebrate-specific and insect-specific GO transfers (**Supplementary Tables 11,12**) For the heatmap in **Fig. 5c**, we exclusively considered GO categories that were either (i) significantly enriched in the gains of at least 15 nodes/species across all tissues or (ii) significantly enriched in the gains of at least 8 nodes/species in one tissue exclusively; in this last analysis, ovary and testis were grouped in order to catch a combined signature from the reproductive organs. The plotted values (log2(observed/expected+1)) were computed starting from the proportion of gains in each node/species belonging to the tested category (observed) and the proportion of all bilaterian-conserved orthogroups with a functional annotation belonging to the same category (expected). Highly redundant categories were manually removed. For **Fig. 5d** and **Supplementary Table 13**, we only considered the GO categories that were exclusively enriched in one node or species. Then, we moved to the characterization of species-specific gains, where we evaluated if developmental GOs were more represented in these recent gains compared to ancestral ones. Developmental GO categories were defined starting from the human transferred GO annotation (see above) as any term matching “develop”, “differentiation”, “determination”, “morphogen”, “commitment”, “specification”, “regionalization”, “formation”, “genesis”. For the plot shown in **Fig. 5e**, only the GO categories including at least 10 bilaterian-conserved orthogroups were considered. The GSEA analysis in **Extended Data Fig. 9c** was performed with the *fgsea* package in R ^80^, and distribution shown in **Extended Data Fig. 9d** resulted from 1000 randomizations of the GO categories labels across the proportions of orthogroups in each category that included at least one species-specific gain.

## Authors’ contribution

FM performed most analyses and generated most figures and tables. LPI built the motif dataset, designed and performed all motif-related analysis, and contributed with intellectual discussion. YM and ATM performed additional analyses and contributed with intellectual discussion. JP, ATM, JC, XFM, FT, DB, SB, TD, MN, PC, FN, HE, MIA, CA, KW, IA, DMC contributed RNA and/or tissue samples. FM and MI wrote the manuscript.

## Acknowledgements

We thank Queralt Tolosa Ramon and Niccoló Arecco for their original drawing of tissue and cell-type icons. We also thank Niccoló Arecco, Nuno Barbosa Morais, Arnau Sebé-Pedrós and Donate Weghorn for their critical feedback on the manuscript. We thank the CRG Genomics Unit for the RNA sequencing. Phylopic icons credits (**Fig. 1a**): to Sarah Werning for the opossum, Soledad Miranda-Rottman for the frog, Tony Ayling and Milton Tan for the elephant shark, Harold N Eyster for the sea urchin, Gareth Monger for the hoverfly, Jamie Whitehouse for the silkworm, George Starr for the mayfly: https://creativecommons.org/licenses/by-nc-sa/3.0/, and Birgit Lang for the centipede)

## Data and code availability

Code availability: All the code used for analysis and figure generation is available on github at https://github.com/fedemantica/bilaterian_GE.

Data availability: The FASTQ and processed files of the RNA-seq samples generated for this project are available at GEO under series GSE205498. The Supplementary Dataset is available at https://data.mendeley.com/datasets/22m3dwhzk6/2

---

**Extended Data Fig. 1:**
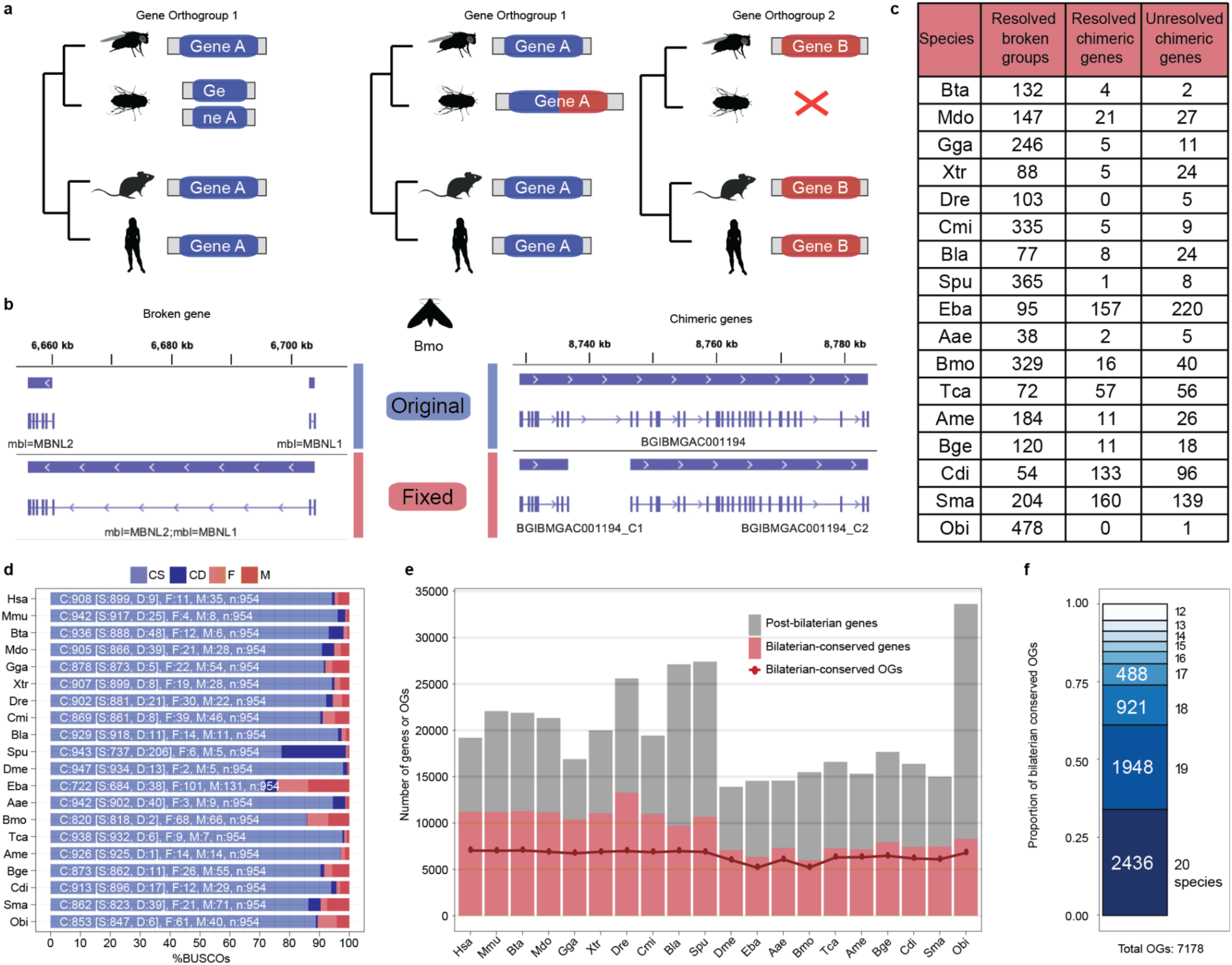
**a.** Schematic representation of broken (left) and chimeric (right) genes and how they potentially influence gene orthology inferences. **b.** Examples of a broken (left) and chimeric (right) genes corrected in the silkworm gene annotation. **c.** Statistics of corrected and unresolved broken and chimeric genes across all species. **d.** Results from a *BUSCO* run (options *-m proteins-l metazoa_odb10*) assessing the status of 954 metazoa single-copy orthologs in the proteomes of all the species. CS: complete and single-copy, CD: complete and duplicated, F: fragmented, M: missing. **e.** Barplot representing the number of bilaterian-conserved (red) or more recent (gray) protein-coding genes across all species. The line plot represents the number of bilaterian-conserved orthogroups (OGs; i.e., orthogroups conserved in at least 12 species) in which genes from each species are represented. **f.** Proportions of bilaterian-conserved orthogroups based on the number of species in which they are conserved.

**Extended Data Fig. 2:**
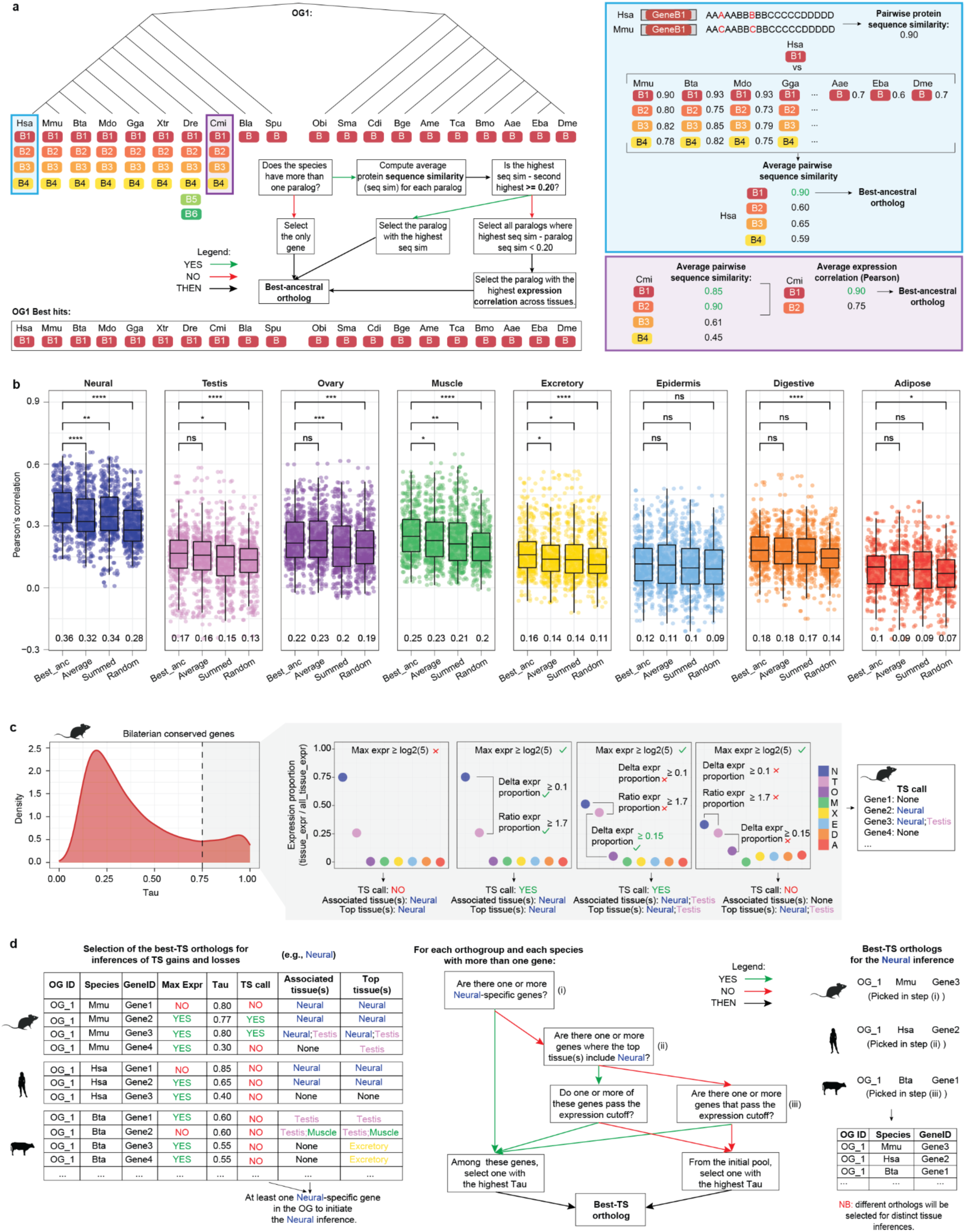
**a.** Schematic and relative example for the selection of bilaterian-conserved, best-ancestral orthologs in each species (see **Methods**). **b.** Distributions of Pearson’s correlation coefficients from all intra-tissue, species pairwise comparisons of gene expression upon distinct procedures for paralog selection and gene expression quantification. The expression measure for each species in each orthogroup corresponds to the expression of its best-ancestral ortholog (Best_anc), the average expression among all its paralogs (Average), the summed expression among all its paralogs (Summed) and the expression of a randomly selected paralog (Random). Significance levels of Wilcoxon rank-sum tests comparing the Best_anc distribution to each of the others are reported at the top, while the median value of each distribution is printed at the bottom. Correlations are performed on z-scored expression matrices (see **Methods**). Only the 2,436 gene orthogroups conserved in all species were considered. c. Schematic of the procedure adopted to associate all tissue-specific genes in each species (Tau ≥ 0.75) with the tissue(s) with tissue-specificity. This association (which we also evaluated for non-tissue-specific genes) will be considered for the inference of tissue-specificity gains (**Extended Data Fig. 5**). Additionally, we identified the top tissue(s) (i.e., the tissue(s) with the highest expression) for all bilaterian-conserved genes, which will be considered for the selection of the best-TS orthogroups and the inference of tissue-specificity losses in each tissue (panel d and **Extended Data Fig. 5c**, respectively). **d.** Schematic and relative example for the selection of the best-TS ortholog in each species (see **Methods**).

**Extended Data Fig. 3:**
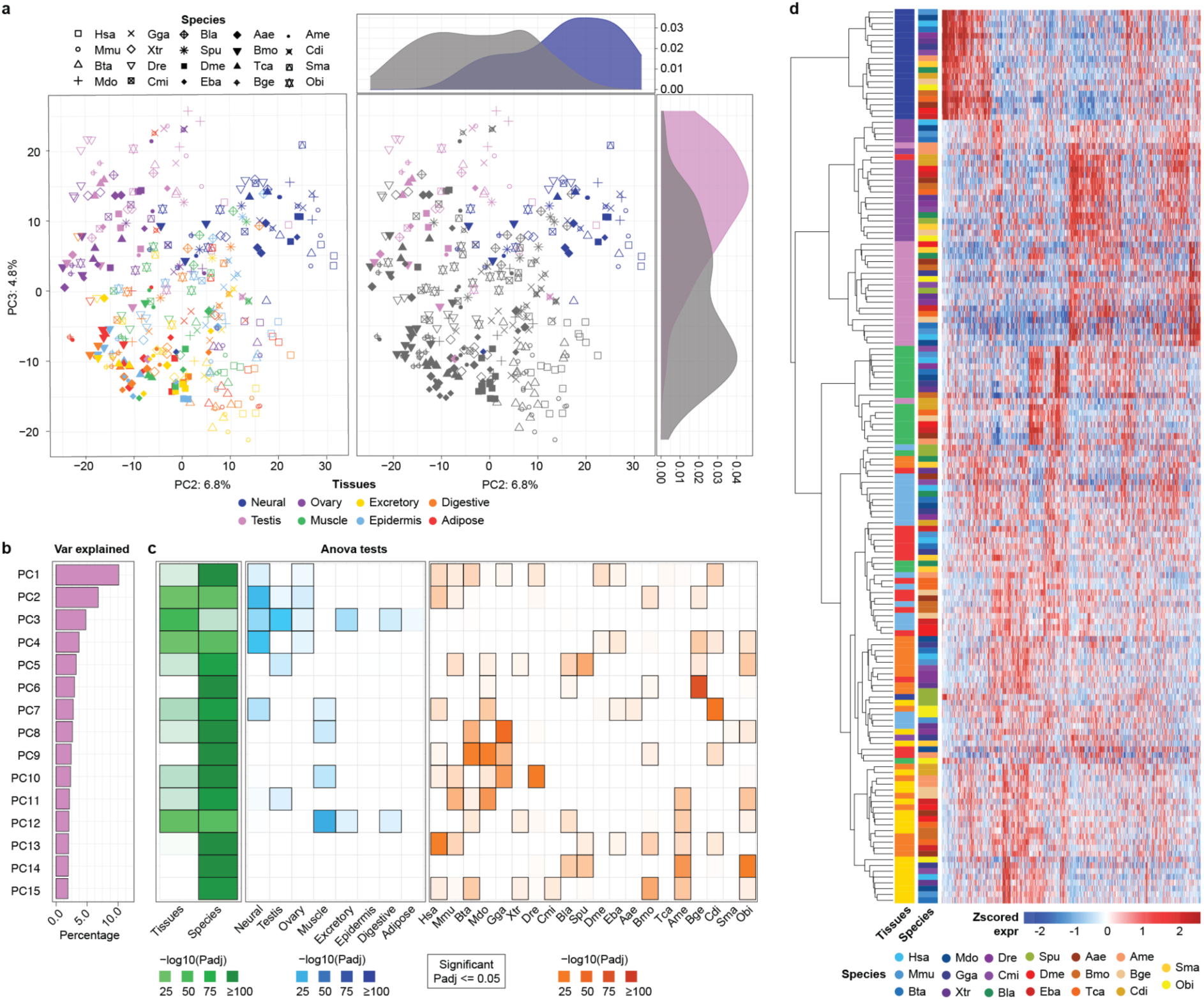
**a.** Coordinates of the second (PC2; x axis) and third (PC3; y axis) components of a PCA performed on the best-ancestral orthogroups normalized gene expression matrix. Only the 2,436 best-ancestral orthogroups conserved in all species were considered. Tissue identity is represented by colors and species by shape. The left panel shows all tissues, while the right panel highlights neural and testis samples compared to all others. Coordinate distributions of these three groups of meta-samples are shown on the side of the relative component. The percentage of variance explained by each PC is reported on the relative axis. **b.** Percentage of variance explained by the first 15 principal components from the PCA described in a. **c.**-log10(p-value) of ANOVA tests performed among the coordinates of the specified groups on each component. For the left panel (green) we tested if there was a significant difference between tissues or species groups. For the center and right panel (blue and orange) we tested if there was a significant difference between any query group (i.e., column) versus all other collapsed groups. All tests were performed with the *aov* function in R, and p-values were Bonferroni corrected. **d.** Heatmap showing the clustering of tissues and species (rows) based on the expression across tissues of best-ancestral bilaterian-conserved orthogroups (columns). Expression values were z-scored across tissues of the same species in order to minimize the inter-species variability (see Methods for the definition of the best-ancestral orthogroups z-scored expression matrix). Only the 2,436 best-ancestral orthogroups conserved in all species were considered. The heatmap was generated by the *pheatmap* function in R with *ward.D2* clustering method. Tissue colors refer to panel a.

**Extended Data Fig. 4:**
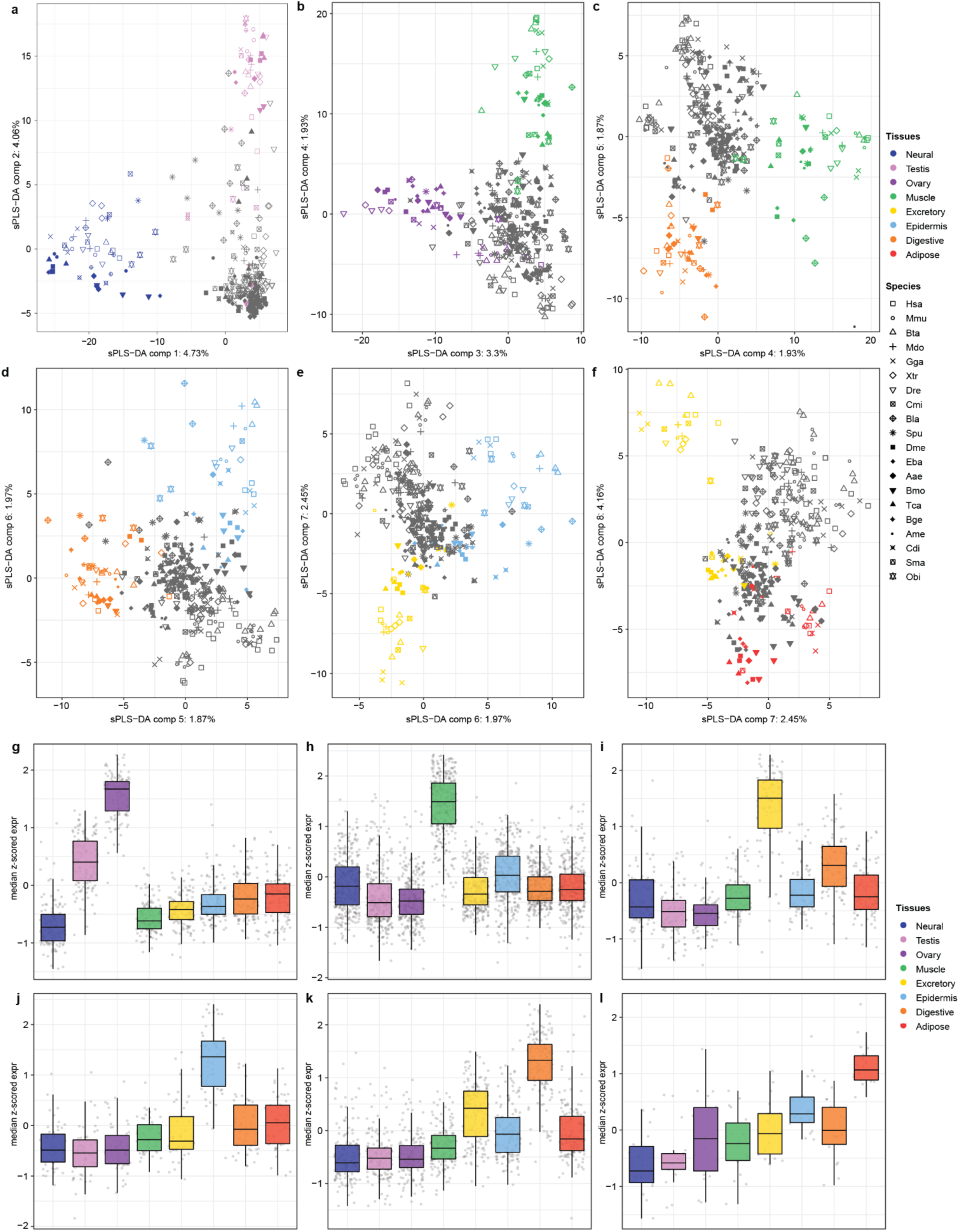
**a-f.** Coordinates of components returned by a sparse partial least square discriminant analysis (sPLS-DA) run separating the meta-samples of each tissue group (depicted with the relative colors) from all the others (gray). All 7,178 best-ancestral orthogroups were considered. The loadings of these components will be used to define the ancestral bilaterian tissue-specific modules (see **Fig. 2a,b**). The percentage of variance explained by each component is reported on the relative axis. **g-l**: Expression profiles across tissues of best-ancestral orthogroups in the ancestral tissue-specific modules (see **Fig. 2c,d** for neural and testis modules). Expression values were first z-scored by species, and each dot represents the median expression among vertebrates, insects or outgroups.

**Extended Data Fig. 5:**
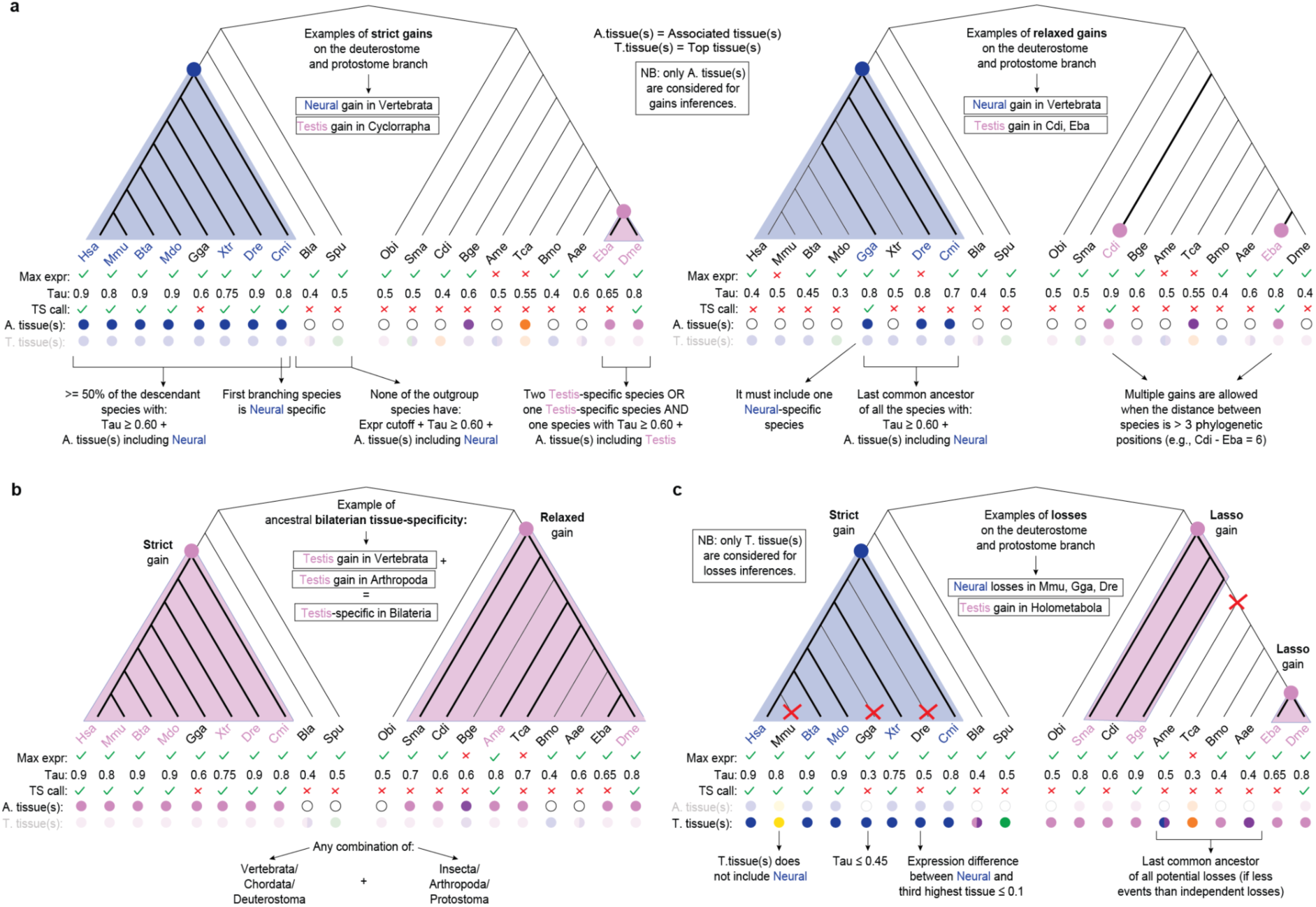
**a.** Examples and criteria for the inference of tissue-specificity gains on either the deuterostome or protostome branches with the strict approach (left panel) and the relaxed approach (right panel). **b.** Example and criteria for the inference of ancestral bilaterian tissue-specificity. **c.** Examples and criteria for the inference of tissue-specificity losses. NB: the best-TS orthogroups are the ones considered for all inferences of tissue-specificity gains and losses (see **Methods** and **Extended Data Fig. 2c,d**).

**Extended Data Fig. 6:**
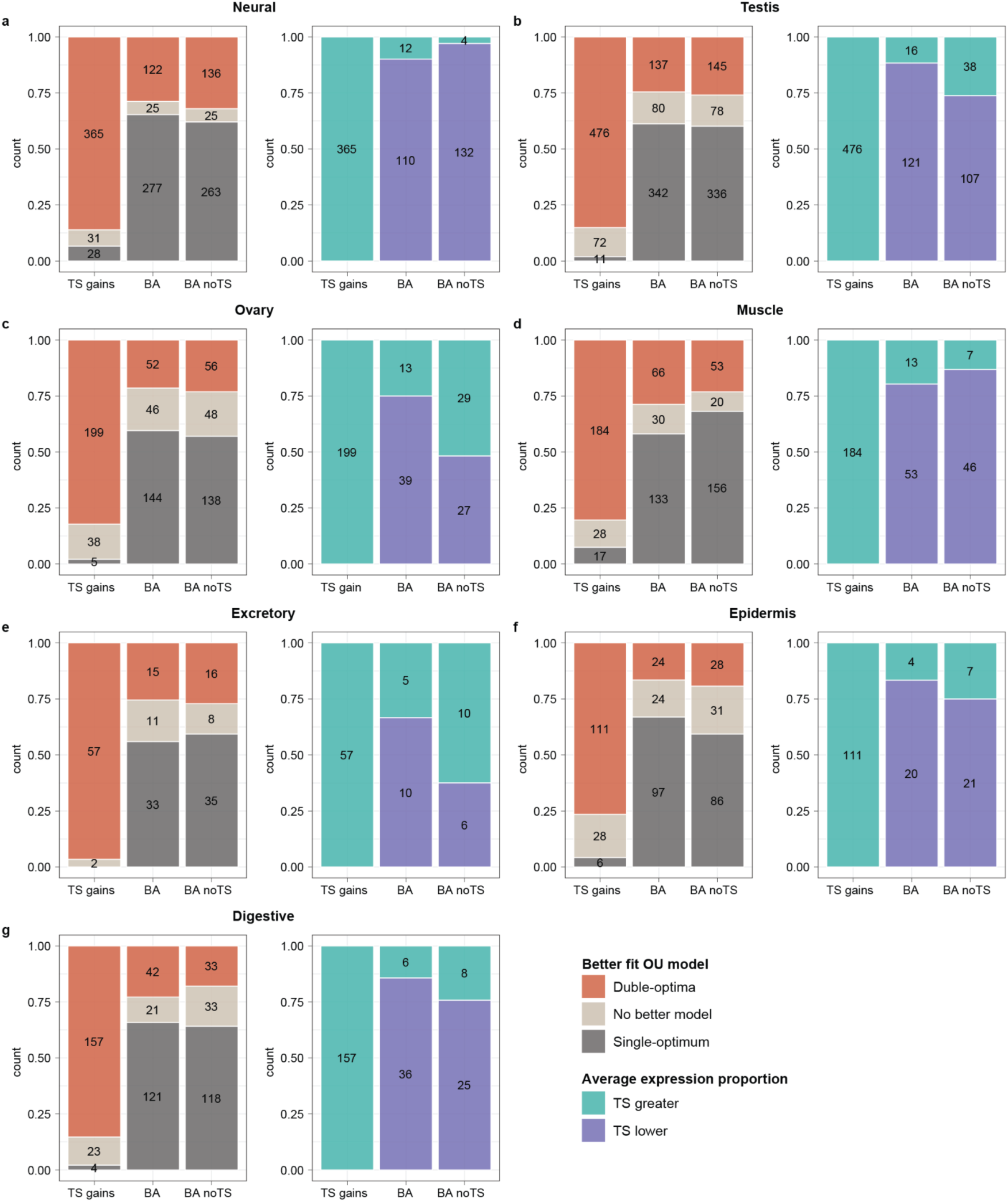
**a-g.** Orthogonal validation of all the inferred tissue-specificity gains in each tissue for which we could implement an OUs comparison method (see **Methods** and **Supplementary Discussion**). The first bar always corresponds to the selected tissue-specificity gains (TS gains), while the second and third bars represent control sets (of the same size as the test set) sampled from either all best-ancestral orthogroups (BA) or best-ancestral orthogroups without tissue-specificity gains (BA no TS), to which we randomly assigned the tissue-specificity labels of the corresponding test set (see **Methods**). *Left barplot*: proportions of orthogroups based on the OU model (either a double-optima or a single-optimum) that better fits the relative expression levels. The double-optima OU model postulates different expression optima for the species with and without tissue-specificity, where the latter also include all species with losses. *Right barplot*: proportions of orthogroups better fitting a double-optima OU model (in red on the left barplot) depending on whether the species with tissue-specificity show higher/lower average relative expression compared to species without (TS greater/lower, respectively).

**Extended Data Fig. 7:**
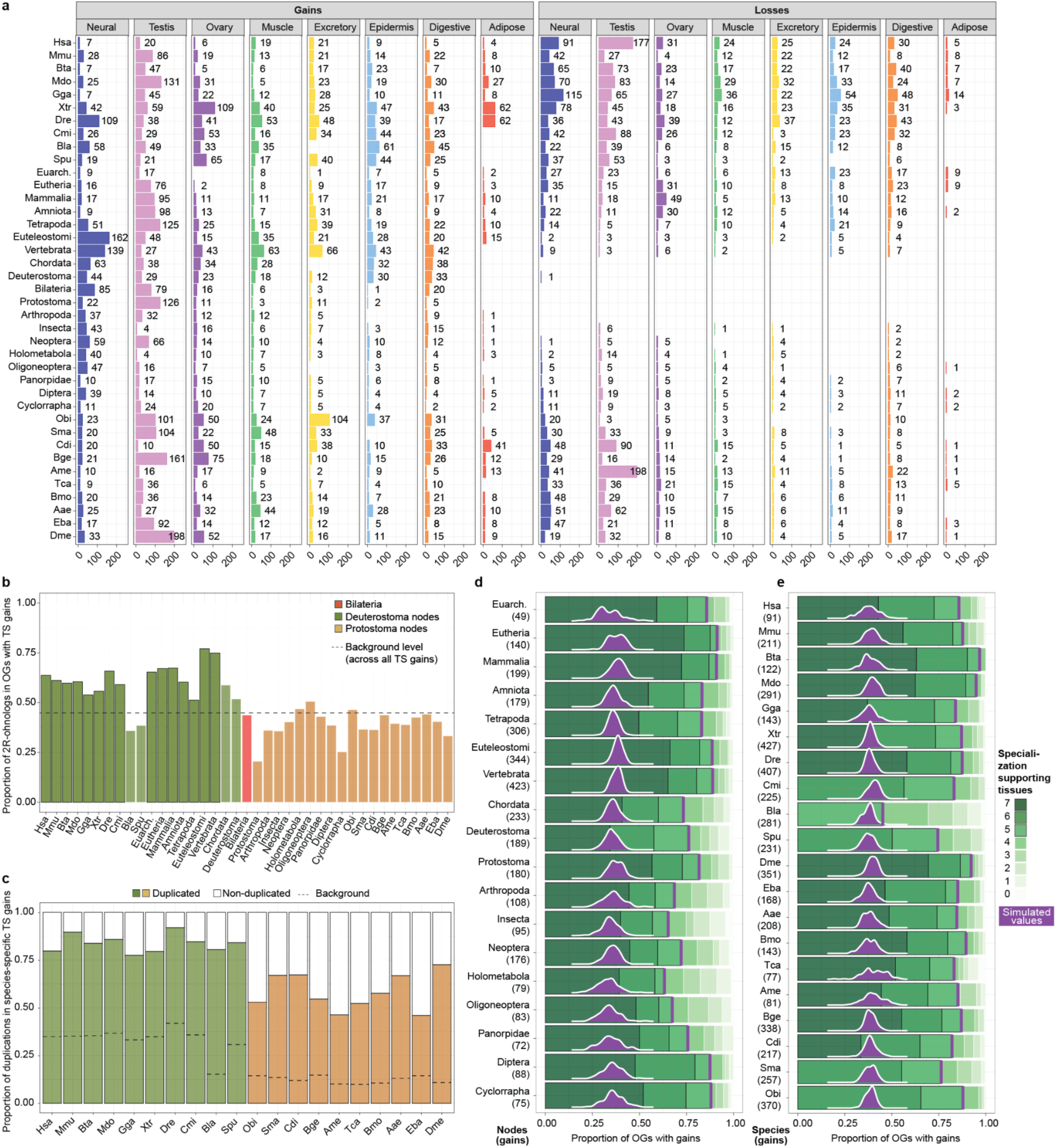
**a.** Barplots representing the number of inferred tissue-specificity gains (left) and losses (right) across all nodes/species (rows) and tissues (columns). Best-TS, bilaterian-conserved orthogroups were considered for these inferences. **b.** Proportion of tissue-specificity gains in each node/species occurring in best-TS orthogroups that include 2R-onhologs. Deuterostome nodes/species are distinguished between those diverging before (transparent color) or after (full color) the two rounds of vertebrate WGDs. The black line represents the proportion of 2R-onhologs across all tissue-specificity gains. **c.** Proportions of duplicated (i.e., with at least one paralog) or non-duplicated (i.e., single-copy) genes with tissue-specific, species-specific gains in all species. The background line represents the overall proportion of duplicated genes in each species. **d,e.** Same data represented in Fig. 4f, but plotted separately across all nodes (d) and species (e). **NB:** Bilaterian “gains” indicate ancestral bilaterian tissue-specificity, which might have been acquired either in the last bilaterian ancestor or previously in evolution. Abbreviations: Euarch: Euarchontoglires.

**Extended Data Fig. 8:**
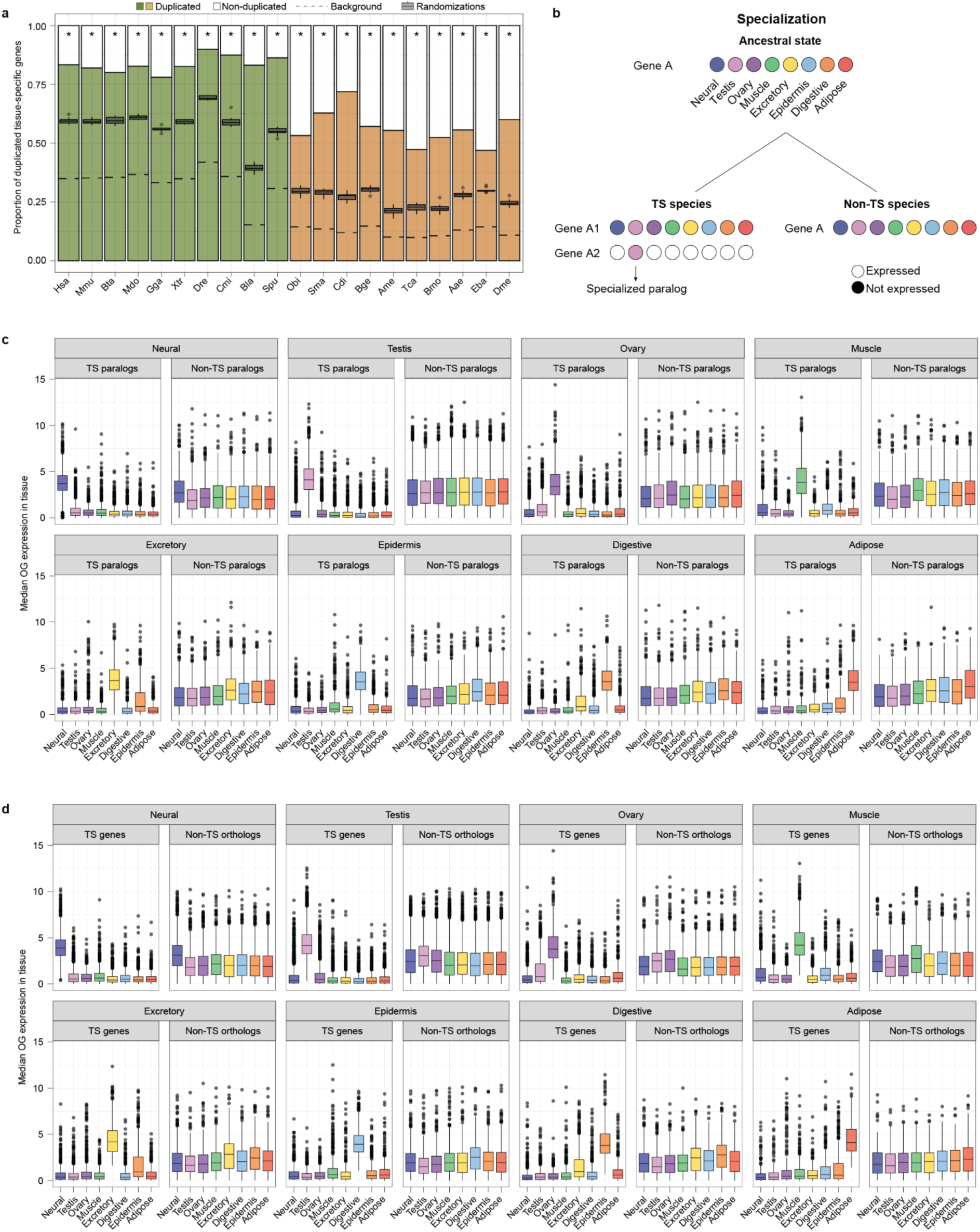
**a.** Barplot: proportions of duplicated (i.e., with at least one paralog) or non-duplicated (i.e., single-copy) tissue-specific genes in each species. Boxplot: proportions of duplicated tissue-specific genes in each species upon the ten randomizations of the original orthogroups (see **Methods**). The asterisks indicate a significant difference (binomial test, p-value ≤ 0.05) between the observed proportion of duplicated tissue-specific genes and the median of such proportions coming from the randomization trials. The background line represents the overall proportion of duplicated genes in each species. **b**. Scheme illustrating how tissue-specific expression can be gained following gene duplication and specialization. Color dots indicate expression in the relative tissue, white dots represent lack of expression. **c.** For each tissue, median gene expression in each bilaterian-conserved orthogroups for species possessing at least one tissue-specific and one non-tissue-specific gene. Expression of tissue-specific genes is plotted on the left, while expression of their non-tissue specific paralogs is shown on the right. Each data point in each tissue’s boxplot is the median of the relative expression in that tissue for all corresponding genes and species. **d.** Median gene expression across tissues for bilaterian-conserved orthogroups with tissue-specific gains in each tissue. Left: best-TS orthologs of the species with tissue-specificity. Right: best-TS orthologs in the other species. Each data point in each tissue’s boxplot is the median of the relative expression in that tissue for all corresponding genes and species. Distributions for gains within single nodes/species are available in the Supplementary Dataset.

**Extended Data Fig. 9:**
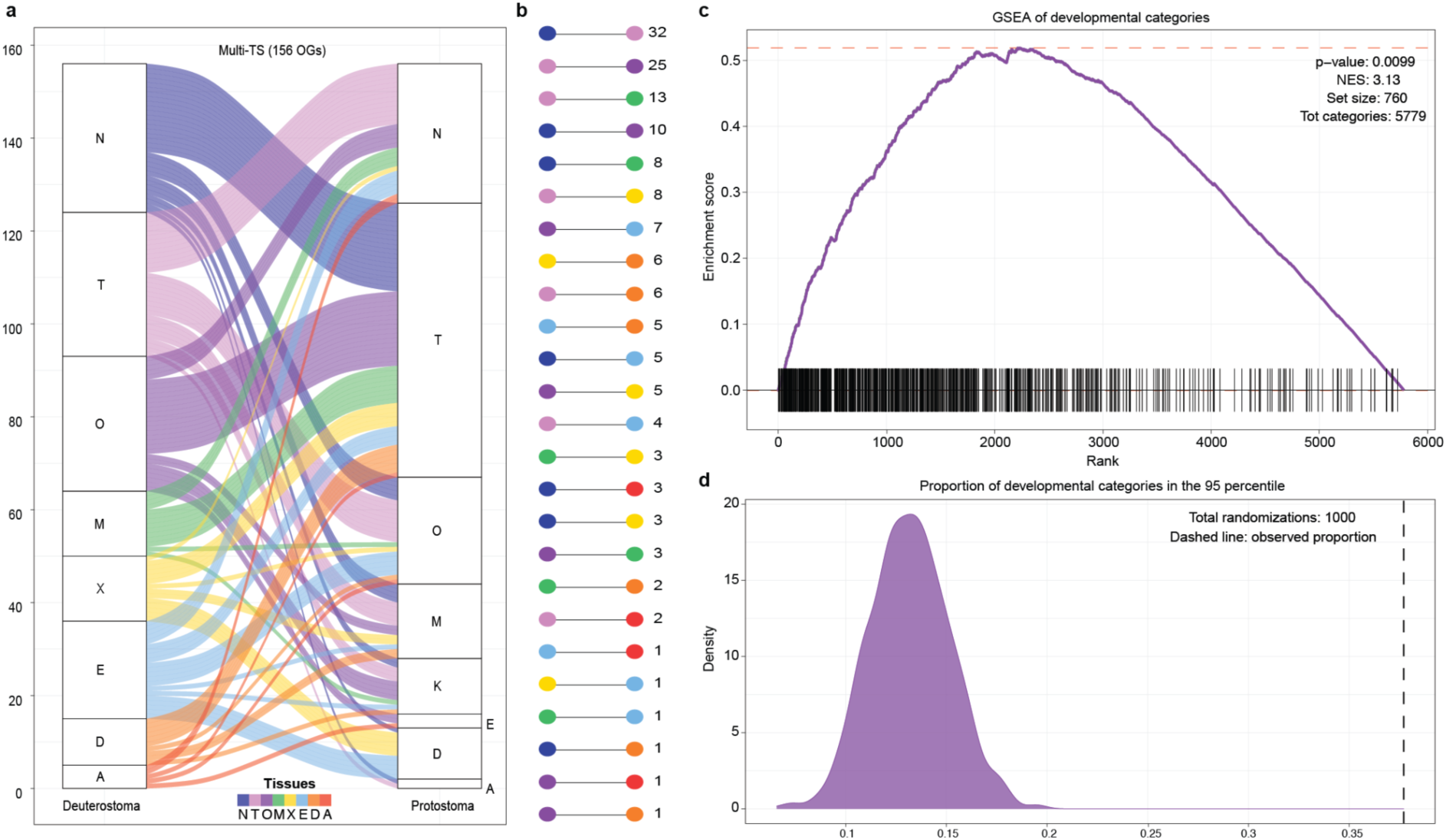
**a.** Alluvia plot representing the best-TS, bilaterian-conserved orthogroups with tissue-specificity gains in distinct tissues between deuterostome (left) or protostome (right) nodes and species. Only orthogroups with gains in exclusively one tissue on each branch were considered. **b**. Number of parallel tissue-specificity gains between the deuterostome and protostome branch for all pairs of tissues represented in panel a. **c.** Plot from a Gene Set Enrichment Analysis (GSEA) testing for over-representation of developmental categories (760 out of 5779) among categories with high proportions of orthogroups that undergo species-specific gains of tissue-specificity. Only categories including at least 10 gene orthogroups were considered. **d.** Proportions of developmental GO categories among the top 5% (i.e., 95th percentile) of all GO categories ranked based on the proportions of their annotated orthogroups that undergo species-specific gains. The plotted values derive from 1000 randomization of the developmental labels among all GO categories, with the vertical dashed line corresponding to the observed proportion.

**Extended Data Fig. 10:**
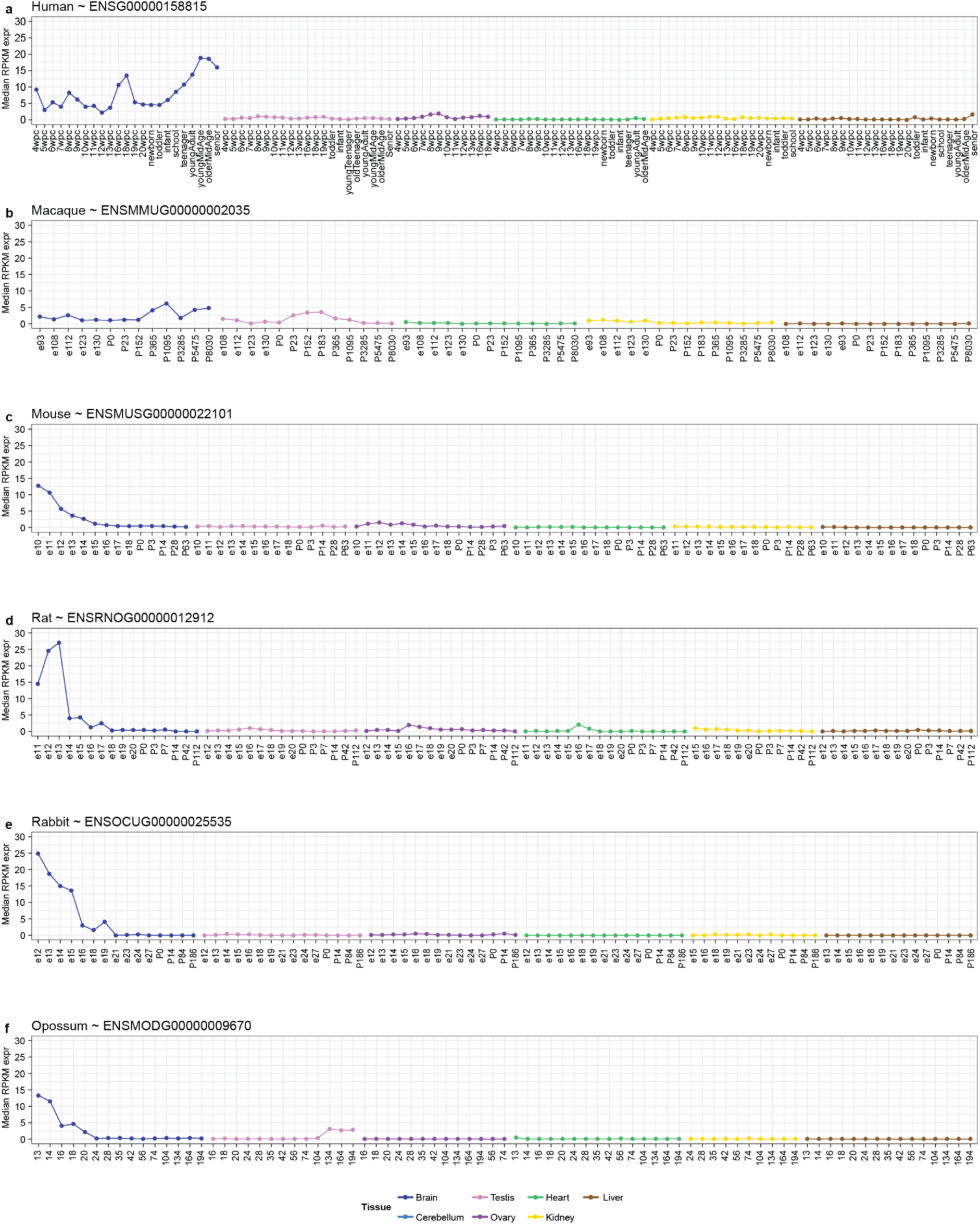
**a-f:** Expression values (RPKMs) for human FGF17 (a) and its orthologs in five mammalian species (b-f) across several developmental and adult timepoints in seven tissues. Data from ^19^.

## References

1. Evans, S. D., Hughes, I. V., Gehling, J. G. & Droser, M. L. Discovery of the oldest bilaterian from the Ediacaran of South Australia. Proc. Natl. Acad. Sci. U. S. A. 117, 7845–7850 (2020).

2. Technau, U. Brachyury, the blastopore and the evolution of the mesoderm. Bioessays 23, 788– 794 (2001).

3. Brusca, Moore & Shuster. Introduction to the Bilateria and the Phylum Xenacoelomorpha Triploblasty and Bilateral. in Invertebrates (ed. Sinauer Associates, Inc) (2016).

4. Paps, J. & Holland, P. W. H. Reconstruction of the ancestral metazoan genome reveals an increase in genomic novelty. Nat. Commun. 9, 1730 (2018).

5. Fernández, R. & Gabaldón, T. Gene gain and loss across the metazoan tree of life. Nat Ecol Evol 4, 524–533 (2020).

6. Lopez-Bigas, N., De, S. & Teichmann, S. A. Functional protein divergence in the evolution of Homo sapiens. Genome Biol. 9, R33 (2008).

7. Davidson, E. H. & Erwin, D. H. Gene regulatory networks and the evolution of animal body plans. Science 311, 796–800 (2006).

8. King, M.-C. & Wilson, A. C. Evolution at Two Levels in Humans and Chimpanzees. Science 188, 107–116 (1975).

9. Carroll, S. B. Evo-devo and an expanding evolutionary synthesis: a genetic theory of morphological evolution. Cell 134, 25–36 (2008).

10. True, J. R. & Carroll, S. B. Gene co-option in physiological and morphological evolution. Annu. Rev. Cell Dev. Biol. 18, 53–80 (2002).

11. Arntfield, M. E. & van der Kooy, D. β-Cell evolution: How the pancreas borrowed from the brain: The shared toolbox of genes expressed by neural and pancreatic endocrine cells may reflect their evolutionary relationship. Bioessays 33, 582–587 (2011).

12. Almudi, I. et al. Genomic adaptations to aquatic and aerial life in mayflies and the origin of insect wings. Nat. Commun. 11, 2631 (2020).

13. Clark-Hachtel, C. M. & Tomoyasu, Y. Two sets of candidate crustacean wing homologues and their implication for the origin of insect wings. Nat Ecol Evol 4, 1694–1702 (2020).

14. Bruce, H. S. & Patel, N. H. Knockout of crustacean leg patterning genes suggests that insect wings and body walls evolved from ancient leg segments. Nat Ecol Evol 4, 1703–1712 (2020).

15. Martín-Durán, J. M. et al. Convergent evolution of bilaterian nerve cords. Nature 553, 45–50 (2018).

16. Thomas, J. A., Welch, J. J., Lanfear, R. & Bromham, L. A Generation Time Effect on the Rate of Molecular Evolution in Invertebrates. Mol. Biol. Evol. 27, 1173–1180 (2010).

17. Wyder, S., Kriventseva, E. V., Schröder, R., Kadowaki, T. & Zdobnov, E. M. Quantification of ortholog losses in insects and vertebrates. Genome Biol. 8, R242 (2007).

18. Brawand, D. et al. The evolution of gene expression levels in mammalian organs. Nature 478, 343– 348 (2011).

19. Cardoso-Moreira, M. et al. Gene expression across mammalian organ development. Nature 571, 505–509 (2019).

20. Chen, J. et al. A quantitative framework for characterizing the evolutionary history of mammalian gene expression. Genome Res. 29, 53–63 (2019).

21. Fukushima, K. & Pollock, D. D. Amalgamated cross-species transcriptomes reveal organ-specific propensity in gene expression evolution. Nat. Commun. 11, 4459 (2020).

22. Barbosa-Morais, N. L. et al. The evolutionary landscape of alternative splicing in vertebrate species. Science 338, 1587–1593 (2012).

23. Lê Cao, K.-A., Boitard, S. & Besse, P. Sparse PLS discriminant analysis: biologically relevant feature selection and graphical displays for multiclass problems. BMC Bioinformatics 12, 253 (2011).

24. Burkhardt, P. & Sprecher, S. G. Evolutionary origin of synapses and neurons-Bridging the gap. Bioessays 39, (2017).

25. Sebé-Pedrós, A. et al. Cnidarian Cell Type Diversity and Regulation Revealed by Whole-Organism Single-Cell RNA-Seq. Cell 173, 1520–1534.e20 (2018).

26. Inaba, K. Sperm flagella: comparative and phylogenetic perspectives of protein components. Mol. Hum. Reprod. 17, 524–538 (2011).

27. Daldello, E. M., Luong, X. G., Yang, C.-R., Kuhn, J. & Conti, M. Cyclin B2 is required for progression through meiosis in mouse oocytes. Development 146, (2019).

28. Li, J., Ouyang, Y.-C., Zhang, C.-H., Qian, W.-P. & Sun, Q.-Y. The cyclin B2/CDK1 complex inhibits separase activity in mouse oocyte meiosis I. Development 146, (2019).

29. Zeng, Y. et al. Bi-allelic mutations in MOS cause female infertility characterized by preimplantation embryonic arrest. Hum. Reprod. 37, 612–620 (2022).

30. Tay, J., Hodgman, R., Sarkissian, M. & Richter, J. D. Regulated CPEB phosphorylation during meiotic progression suggests a mechanism for temporal control of maternal mRNA translation. Genes Dev. 17, 1457–1462 (2003).

31. Gąsiorowski, L. et al. Molecular evidence for a single origin of ultrafiltration-based excretory organs. Curr. Biol. 31, 3629–3638.e2 (2021).

32. Thakurela, S. et al. Mapping gene regulatory circuitry of Pax6 during neurogenesis. Cell Discov 2, 15045 (2016).

33. Eckler, M. J. & Chen, B. Fez family transcription factors: controlling neurogenesis and cell fate in the developing mammalian nervous system. Bioessays 36, 788–797 (2014).

34. Taylor, M. V. & Hughes, S. M. Mef2 and the skeletal muscle differentiation program. Semin. Cell Dev. Biol. 72, 33–44 (2017).

35. Mathiyalagan, N. et al. Meta-Analysis of Grainyhead-Like Dependent Transcriptional Networks: A Roadmap for Identifying Novel Conserved Genetic Pathways. Genes 10, (2019).

36. Yanai, I. et al. Genome-wide midrange transcription profiles reveal expression level relationships in human tissue specification. Bioinformatics 21, 650–659 (2005).

37. Roelofs, D. et al. Multi-faceted analysis provides little evidence for recurrent whole-genome duplications during hexapod evolution. BMC Biol. 18, 57 (2020).

38. Marlétaz, F. et al. Amphioxus functional genomics and the origins of vertebrate gene regulation. Nature 564, 64–70 (2018).

39. Oji, A. et al. Tesmin, Metallothionein-Like 5, is Required for Spermatogenesis in Mice†. Biol. Reprod. 102, 975–983 (2020).

40. Jiang, J., Benson, E., Bausek, N., Doggett, K. & White-Cooper, H. Tombola, a tesmin/TSO1-family protein, regulates transcriptional activation in the Drosophila male germline and physically interacts with always early. Development 134, 1549–1559 (2007).

41. Hines, J. H. Evolutionary Origins of the Oligodendrocyte Cell Type and Adaptive Myelination. Front. Neurosci. 15, 757360 (2021).

42. Ramirez, M. D. & Oakley, T. H. Eye-independent, light-activated chromatophore expansion (LACE) and expression of phototransduction genes in the skin of Octopus bimaculoides. J. Exp. Biol. 218, 1513–1520 (2015).

43. Iram, T. et al. Young CSF restores oligodendrogenesis and memory in aged mice via Fgf17. Nature 605, 509–515 (2022).

44. Hartenstein, V. & Martinez, P. Structure, development and evolution of the digestive system. Cell Tissue Res. 377, 289–292 (2019).

45. Ottaviani, E., Malagoli, D. & Franceschi, C. The evolution of the adipose tissue: a neglected enigma. Gen. Comp. Endocrinol. 174, 1–4 (2011).

46. Kryuchkova-Mostacci, N. & Robinson-Rechavi, M. Tissue-Specificity of Gene Expression Diverges Slowly between Orthologs, and Rapidly between Paralogs. PLoS Comput. Biol. 12, e1005274 (2016).

47. Lien, S. et al. The Atlantic salmon genome provides insights into rediploidization. Nature 533, 200– 205 (2016).

48. Fernández, R. et al. Selection following Gene Duplication Shapes Recent Genome Evolution in the Pea Aphid Acyrthosiphon pisum. Mol. Biol. Evol. 37, 2601–2615 (2020).

49. Farré, D. & Albà, M. M. Heterogeneous patterns of gene-expression diversification in mammalian gene duplicates. Mol. Biol. Evol. 27, 325–335 (2010).

50. Clark, J. W. & Donoghue, P. C. J. Constraining the timing of whole genome duplication in plant evolutionary history. Proc. Biol. Sci. 284, (2017).

51. Macqueen, D. J. & Johnston, I. A. A well-constrained estimate for the timing of the salmonid whole genome duplication reveals major decoupling from species diversification. Proc. Biol. Sci. 281, 20132881 (2014).

52. Donoghue, P. C. J. & Purnell, M. A. Genome duplication, extinction and vertebrate evolution. Trends Ecol. Evol. 20, 312–319 (2005).

53. Almudí, I. & Pascual-Anaya, J. How Do Morphological Novelties Evolve? Novel Approaches to Define Novel Morphologies. in Old Questions and Young Approaches to Animal Evolution (eds. Martín-Durán, J. M. & Vellutini, B. C.) 107–132 (Springer International Publishing, 2019).

54. Brasó-Vives, M. et al. Parallel evolution of amphioxus and vertebrate small-scale gene duplications. bioRxiv 2022.01.18.476203 (2022) doi:10.1101/2022.01.18.476203.

55. Doyle, T. et al. Genome-wide transcriptomic changes reveal the genetic pathways involved in insect migration. Mol. Ecol. 31, 4332–4350 (2022).

56. Derelle, R., Philippe, H. & Colbourne, J. K. Broccoli: Combining Phylogenetic and Network Analyses for Orthology Assignment. Mol. Biol. Evol. 37, 3389–3396 (2020).

57. Manni, M., Berkeley, M. R., Seppey, M., Simão, F. A. & Zdobnov, E. M. BUSCO Update: Novel and Streamlined Workflows along with Broader and Deeper Phylogenetic Coverage for Scoring of Eukaryotic, Prokaryotic, and Viral Genomes. Mol. Biol. Evol. 38, 4647–4654 (2021).

58. Cunningham, F. et al. Ensembl 2022. Nucleic Acids Res. 50, D988–D995 (2022).

59. Bindea, G. et al. ClueGO: a Cytoscape plug-in to decipher functionally grouped gene ontology and pathway annotation networks. Bioinformatics 25, 1091–1093 (2009).

60. Bray, N. L., Pimentel, H., Melsted, P. & Pachter, L. Near-optimal probabilistic RNA-seq quantification. Nat. Biotechnol. 34, 525–527 (2016).

61. Love, M. I., Huber, W. & Anders, S. Moderated estimation of fold change and dispersion for RNA-seq data with DESeq2. Genome Biol. 15, 550 (2014).

62. Tapial, J. et al. An atlas of alternative splicing profiles and functional associations reveals new regulatory programs and genes that simultaneously express multiple major isoforms. Genome Res. 27, 1759–1768 (2017).

63. Briggs, J. A. et al. The dynamics of gene expression in vertebrate embryogenesis at single-cell resolution. Science 360, (2018).

64. Katoh, K. & Standley, D. M. MAFFT multiple sequence alignment software version 7: improvements in performance and usability. Mol. Biol. Evol. 30, 772–780 (2013).

65. Ritchie, M. E. et al. limma powers differential expression analyses for RNA-sequencing and microarray studies. Nucleic Acids Res. 43, e47 (2015).

66. Rohart, F., Gautier, B., Singh, A. & Lê Cao, K.-A. mixOmics: An R package for ‘omics feature selection and multiple data integration. PLoS Comput. Biol. 13, e1005752 (2017).

67. Kolberg, L., Raudvere, U., Kuzmin, I., Vilo, J. & Peterson, H. gprofiler2--an R package for gene list functional enrichment analysis and namespace conversion toolset g:Profiler. F1000Res. 9, (2020).

68. Supek, F., Bošnjak, M., Škunca, N. & Šmuc, T. REVIGO summarizes and visualizes long lists of gene ontology terms. PLoS One 6, e21800 (2011).

69. Gramates, L. S. et al. FlyBase: a guided tour of highlighted features. Genetics 220, (2022).

70. Weirauch, M. T. et al. Determination and inference of eukaryotic transcription factor sequence specificity. Cell 158, 1431–1443 (2014).

71. Bruse, N. & van Heeringen, S. J. GimmeMotifs: an analysis framework for transcription factor motif analysis. bioRxiv 474403 (2018) doi:10.1101/474403.

72. Nguyen, N. T. T. et al. RSAT 2018: regulatory sequence analysis tools 20th anniversary. Nucleic Acids Res. 46, W209–W214 (2018).

73. Jin, L. et al. A pig BodyMap transcriptome reveals diverse tissue physiologies and evolutionary dynamics of transcription. Nat. Commun. 12, 3715 (2021).

74. Wang, Z.-Y. et al. Transcriptome and translatome co-evolution in mammals. Nature 588, 642–647 (2020).

75. Guschanski, K., Warnefors, M. & Kaessmann, H. The evolution of duplicate gene expression in mammalian organs. Genome Res. 27, 1461–1474 (2017).

76. Beaulieu, J. M. & O’Meara, B. OUwie: analysis of evolutionary rates in an OU framework. R package version.

77. Cavanaugh, J. E. & Neath, A. A. The Akaike information criterion: Background, derivation, properties, application, interpretation, and refinements. Wiley Interdiscip. Rev. Comput. Stat. 11, e1460 (2019).

78. Mistry, J. et al. Pfam: The protein families database in 2021. Nucleic Acids Res. 49, D412–D419 (2021).

79. Touceda-Suárez, M. et al. Ancient Genomic Regulatory Blocks Are a Source for Regulatory Gene Deserts in Vertebrates after Whole-Genome Duplications. Mol. Biol. Evol. 37, 2857–2864 (2020).

80. Korotkevich, G. et al. Fast gene set enrichment analysis. bioRxiv 060012 (2021) doi:10.1101/060012.

81. Kumar, S. et al. TimeTree 5: An Expanded Resource for Species Divergence Times. Mol. Biol. Evol. 39, (2022).

