## Supplementary Methods and Figures for "Evolution of tissue-specific expression of ancestral genes across vertebrates and insects"

#### Pre-orthology genome annotation enrichments

For some of the species, we performed genome annotation enrichments which were directly integrated in the gene orthology call (see below). For *Bombyx mori* (Bmo) and *Blattella germanica* (Bge) we used the newly generated RNA-seq samples (see relative Methods section) to infer previously missed gene models by applying Hisat2 [1], StringTie [2] and TransDecoder [3]. We then run *Broccoli* (v1.2) [4] on the selected protein isoforms for the 20 species, and we integrated in the Bmo/Bge genome annotation all newly predicted genes included into gene orthogroups where none of the genes from the original annotations were represented (Bmo: 862 genes; Bge: 1082 genes). For *Strongylocentrotus purpuratus* (Spu), which is prone to erroneously annotated in-tandem duplication, we only selected one gene from each of the gene pairs (49) which were (i) included in the same orthogroups, (ii) sequentially annotated on the same chromosome/strand and (iii) characterized by reciprocal sequence similarity >=0.98.

#### Post-orthology genome annotation and gene orthogroups refinements

We used the information provided by the gene orthology call to correct broken genes (a single gene annotated as two or more separated entities) and chimeric genes (independent genes annotated as a fused element) from the genome annotations, which in the context of orthology analysis can respectively result in erroneous gene duplication inference or incomplete ortholog detection. We defined as broken genes all genes from the same species that **(1)** belonged to the same gene orthogroup, **(2)** were sequentially annotated in the same chromosome/scaffold and in the same strand, **(3)** showed low reciprocal sequence similarity (<30%, calculated based on BLOSUM62 using *mafft* [5] with default parameters), and **(4)** matched different portions of their longest human gene ortholog (overlap cut-off: 25%). All identified broken genes were joined into a unique gene entity with a novel GeneID, which replaced the original entries both in the GTFs and in the gene orthogroups (see **Supplementary Table 1** and **Supplementary Dataset**). With regards to chimeric genes, we defined them as those that were assigned by *Broccoli* to two or more gene orthogroups. We tried to determine the ideal splitting point for each chimeric gene by generating multiple alignments within all the gene orthogroups they belonged to (aligned by *mafft* [5] with parameters *--retree 2 --localpair --maxiterate 1000*), specifically isolating the fragment of the chimeric gene aligning to each orthogroup. Aligned fragment is defined as the portion of the chimeric gene contained between two aminoacids included in continuously aligned sequences of at least 10 amino-acids and aligned with at least 60% of the other genes in the orthogroups, excluding other eventual chimeric genes. We used the aligned fragments to classify the chimeric genes in categories: **(A)** Complete overlap: one aligned fragment spans all the others. The chimeric gene was uniquely assigned to the gene orthogroup with the longest aligned fragment (i.e., unresolved); **(B)** Different exons, continuous: two aligned fragments end up in two adjacent exons, whose boundary represents the splitting point; **(C)** Different exons, not continuous: two aligned fragments end up in not-adjacent exons. The terminal exons of each aligned fragment became the boundaries of the corrected genes, and all the in-between exons were removed from the annotation; **(D)** Different exons, overlap: two aligned fragments share more than one common exon. If only two exons are shared, each of them is assigned to one of the corrected genes. If more exons are shared, the chimeric gene is uniquely assigned to the gene orthogroup with the longest aligned fragment (i.e., unresolved); **(E)** Same exon, not overlap: two aligned fragments align to the same terminal exon, but in different portions; **(F)** Same exon, overlap: two aligned fragments align to overlapping portions of the same terminal exon. Both in **(E)** and **(F)**, the disputed exon is assigned to the aligned fragment with the greatest overlap proportion. While many chimeric genes represent annotation mistakes, we considered as potentially fused genes all those chimeric genes shared between gene orthogroups which contain chimeric genes from at least two species and share exactly the same set of chimeric genes (we identified 75 such genes, divided into 48 gene orthogroups). See **Extended Data Fig. 1** for complete statistics of fixed broken genes, resolved and unresolved chimeric genes, and **Supplementary Table 1** for gene IDs correspondences and other information.

In addition, we used a manually curated dataset of human ohnologs (see **Supplementary Table 2** and [6]) to combine the different orthogroups to which known human ohnologs might have been erroneously assigned. This step reduces the number of gene orthologs from 15,516 to 14,941. Then, to avoid processing large gene families predating the origin of Bilateria, we selected only the 14,713 gene orthogroups either containing less than 80 genes or less than 120 genes where none of the species' genes represent more than 20% of the gene orthogroup. See **Supplementary Table 2** for post-correction gene orthogroups related statistics. Finally, we defined as bilaterian-conserved the 7,178 gene orthogroups conserved in at least 12 species, 2,436 of which resulted to be present in all species. Importantly, only genes with expression equal or higher to 1 TPM in at least one tissue were included in the bilaterian-conserved orthogroups.

**Supplementary Discussion**

**Aim**

This supplementary discussion aims at (1) explaining the rationale behind the choice of an ad-hoc, parsimony-based approach for the inferences of tissue-specificity gains and losses and (2) validating the resulting inferences with an orthogonal method

**Background**

The identification of evolutionary expression shifts is a long-standing challenge of comparative transcriptomic studies. The procedure adopted to identify such shifts can vary, but it relies on the understanding of how gene expression evolves in the clades of interest. In fact, depending on the general patterns of gene expression evolution, different models should be used to predict significant alteration of these patterns from their expected trajectory (and hence, to identify the genes undergoing an evolutionary expression shift). Two approaches have historically been most employed to model gene expression evolution across different clades: (a) brownian models, which assume neutral evolution of gene expression, and (b) Ornstein-Uhlenbeck (OU) models, which postulate the existence of a stabilizing selective pressure that forces the expression divergence between species to reach a plateau, notwithstanding their increasing evolutionary distances. On one hand, gene expression among closely-related species generally evolves neutrally, and better fits a brownian model. For instance, a recent paper [7] released a software based on a bounded brownian model which coherently recapitulated expected evolutionary expression shifts in a specific tomato clade (diverging time: 2.5 MY). On the other hand, gene expression among more distantly-related species tends to evolve following an OU model, as it has been demonstrated both for mammal (diverging time: ~200 MY) [8] and drosophila (diverging time: ~33 MY) [9] species across different tissue types. This knowledge was integrated into previous studies of gene expression evolution in vertebrates [8,10,11], which inferred tissue-specific expression shifts for all genes whose expression profiles better fit an OU model assuming a double expression optima (double-optima OU) compared to a OU model postulating a single expression optimum (single-optimum OU). We will refer to this approach as the “OUs comparison method” throughout the following paragraphs.

**Rationale**

As expected, considering that we work at even larger evolutionary distances compared to the clades investigated in [8,9], the expression of our best-ancestral orthogroups consistently evolves according to an OU model in all tissues and on both branches of our phylogeny (**Supplementary Fig. 10**). However, the standard OUs comparison method presents several limitations that make it unsuitable to infer tissue-specificity gains/losses as per our definition and in this particular phylogenetic scenario:

- The majority of the described instances of the OUs comparison method infer tissue-specific expression shifts exclusively based on the expression level in the queried tissue (usually represented as RPKMs/TPMs) [8,10]. This implies that the expression context in all the other tissues is not taken into account, with two main drawbacks. First, the RPKM/TPM ranges of conserved genes at very large evolutionary distances can vary considerably (e.g. not only due to evolutionary divergence, but also because of differences in gene content or annotation). This could potentially lead to identification of expression shifts even if a gene’s relative expression across tissues remains constant. Second, by exclusively considering the expression in the queried tissue, tissue-specificity gains originated by mechanisms such as specialization (which involves loss of expression in all other tissues) are undetectable by definition. Notably, these limitations could be overcome by implementing an OUs comparison method that considers expression proportions across tissues.
- Some applications of the OUs comparison method can indeed consider multiple traits (e.g., the expression levels across all tissues of interest), as implemented by [11] through the method developed by [12]. While the simultaneous evaluation of expression proportions across all tissues would provide the power to detect wider ranges of expression shifts (i.e., specialization cases),

this particular implementation does not assure the identification of tissue-specificity gains/losses as per our definition: first, it cannot distinguish between changes occuring in one/two tissues from shifts simultaneously involving several of them; second, losses of tissue-specificity would not rely on previously detected gains.

- To the best of our knowledge, the available implementations of the OUs comparison method do not allow missing orthologs. This would prevent us from inferring tissue-specific expression shifts for the majority of our bilaterian-conserved orthogroups, as only ~30% of them are conserved in all species (see **Extended Data Fig. 1f**). However, ~50% of these orthogroups are missing representative orthologs in only 1-3 species, which is highly expected when working at such large evolutionary distances and including recently annotated genomes. Thus, many of these orthogroups represent valid and potentially interesting cases, which could be included in the analysis by a method able to handle few missing values.
- Related to the previous point and the handling of missing values, the OU comparison method cannot adopt a flexible behavior depending on the expression level. In fact, its standard implementation entails an initial filtering that selects only the orthogroups where all the genes pass a pre-determined expression cutoff. However, many of the excluded orthogroups based on this criteria (e.g. because the expression in one/some of the species is lower than the cut-off) from our dataset would represent potentially interesting cases, which could be evaluated if the model contemplated different strategies for genes passing or not a defined expression cutoff.
- Importantly, the OU comparison method requires the a-priori specification of two groups of species with putatively different expression optima, among which the better fit of a double-optima OU model and a single-optimum OU model will be tested. In our framework, testing all clade combinations for all bilaterian-conserved orthogroups across all tissues would be very detrimental in terms of multiple test correction (~2,200,000 tests if testing all bilaterian-conserved orthogroups across all tissues), allowing only a partial reconstruction of the total landscape of tissue-specific expression shifts.

In order to overcome these limitations, we decided to implement a parsimony-based approach that (1) considers each gene's Tau and its relative expression profile across tissues; (2) can handle cases with a few missing orthologs; (3) implements a flexible algorithm that takes into account the expression cutoff and (4) does not require any a-priori hypothesis on the location of the expression shift (see **Methods**). We then attempted to orthogonally validate our approach through the OUs comparison method (see **Extended Data Fig. 6** and **Methods**).

**Validation**

Our inference approach provides defined sets of species with and without tissue-specificity for each inferred tissue-specificity gain, allowing for a direct assessment of whether the expression profiles underlying such gains better fit a double-optima OU model compared to a single-optimum one. Thus, for each tissue separately, we selected all inferred tissue-specificity gains for which a standard OU comparison method could be implemented (8-33% of the total), generating two control groups per test set (see **Methods**). 76-97% of orthogroups with inferred tissue-specificity gains showed evidence for a dual optima in the tested tissue, all of them with shifts in the expected direction (i.e., increased expression in the species with tissue-specificity), as opposed to 16-32% in the relative controls. Moreover, only 2-7% presented a single optimum, in contrast with the 56-68% of the control groups.

### Supplementary Figures

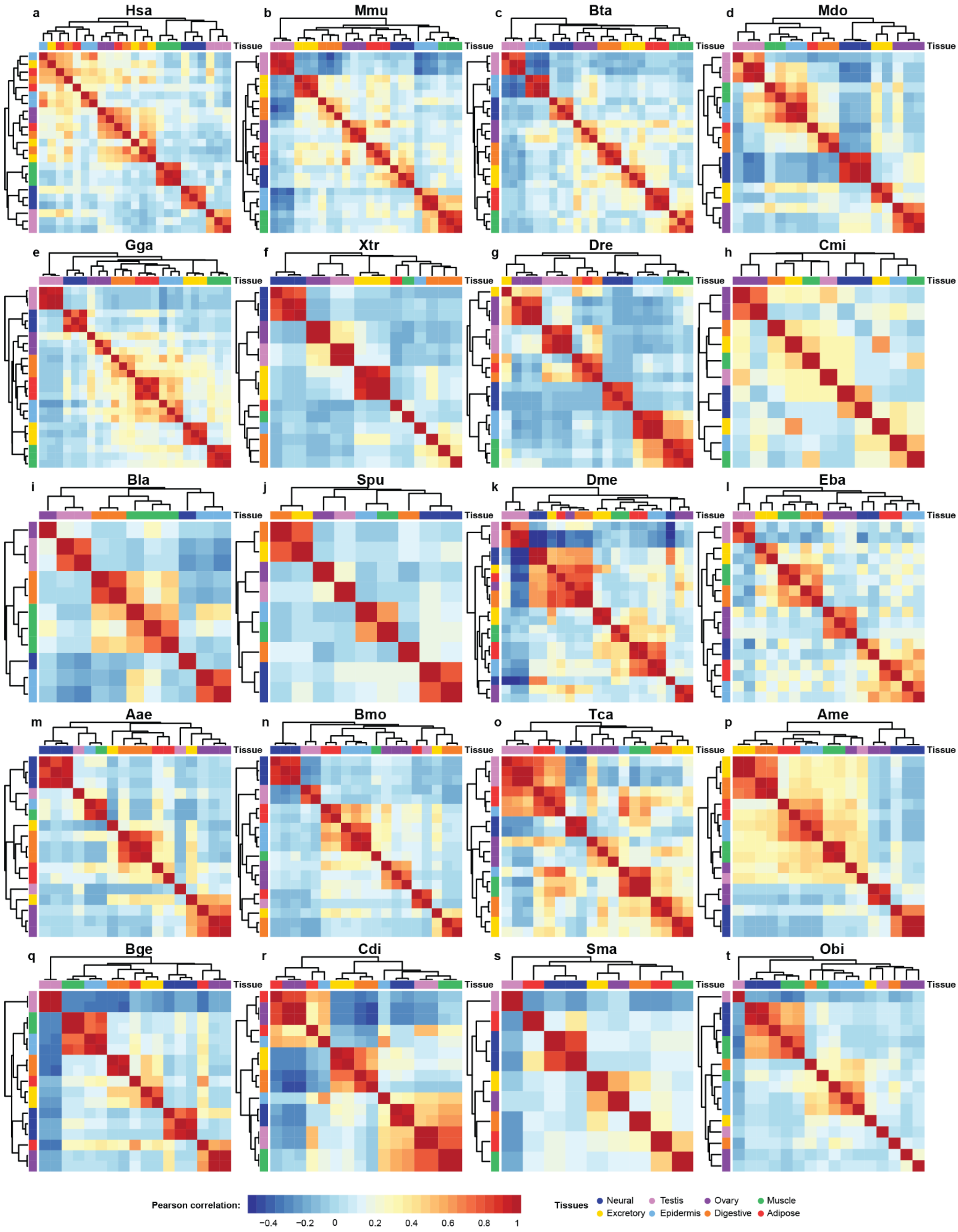


**Supplementary Fig. 1: a-t.** Clustering of each species' meta-samples based on their expression correlation. Expression correlation is represented by Pearson coefficient computed on log2(TPM+1) meta-sample expression values (see **Methods**), where only the 2,500 genes with the highest coefficient of variation were considered in each species. The heatmaps were generated by the *pheatmap* function in R with default clustering parameters.

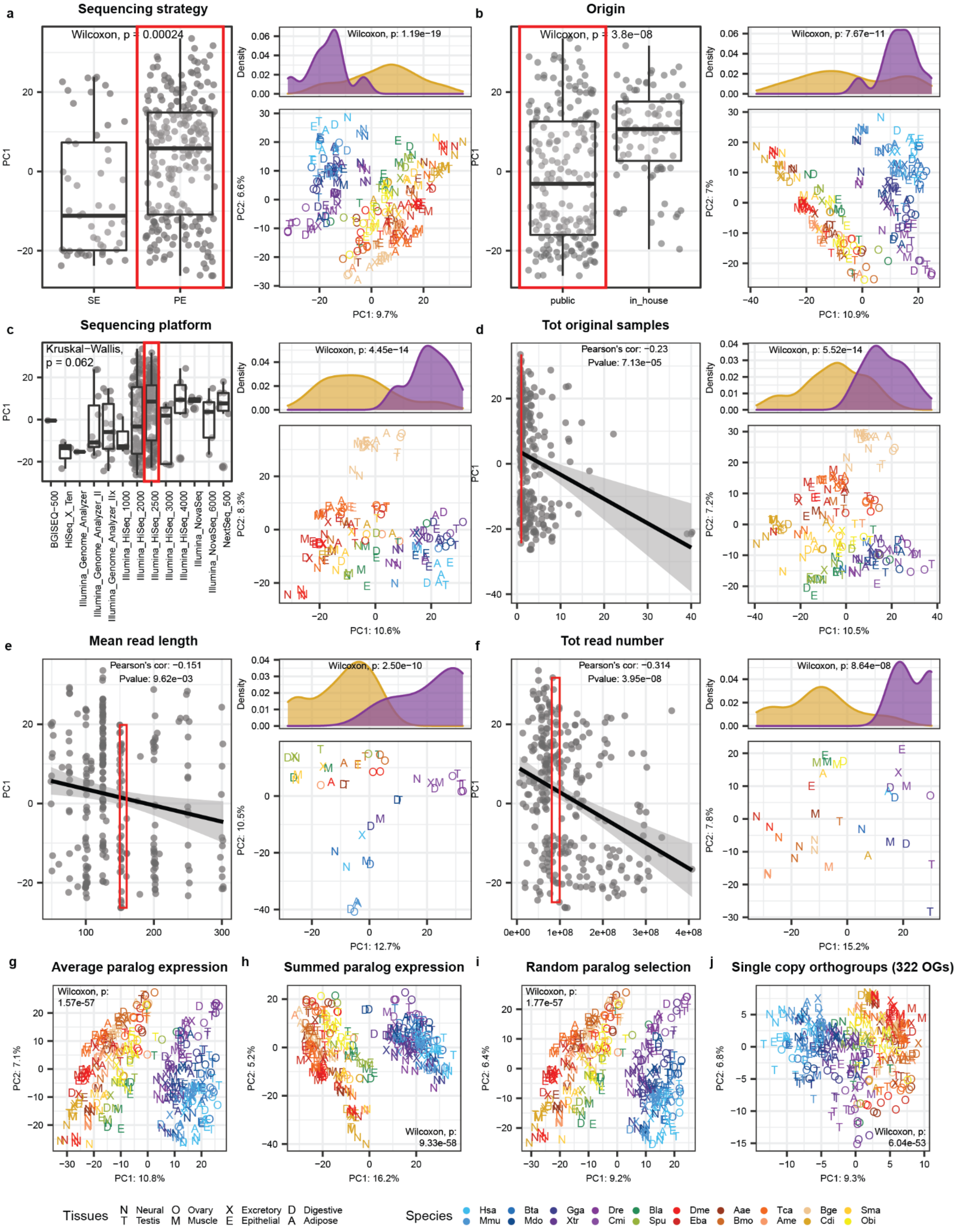


**Supplementary Fig. 2: a-f. Left**: Association between meta-samples' coordinates on the first component of the PCA shown in Fig. 1b (y axis) and covariates potentially linked to batch effects (x axis). The coordinates are plotted after dividing the meta-samples according to their covariate status for the categorical covariates (a,b,c) or in function of the covariate value for continuous covariates (d,e,f). The p-value from a Wilcoxon/Kruskal Wallis test or the coefficient and p-value from a Pearson's correlation test are reported for the categorical and continuous covariates, respectively. The regression line from a linear regression test is added for the continuous covariates. The red rectangle in each plot indicates the category/interval from which meta-samples are selected as input for the PCA shown in the relative right-bottom panel. See **Methods** for details about the meta-samples' selection procedure. **Right-bottom:** PCAs as in Fig. 1b but exclusively performed on the selected meta-samples (red rectangle on the left panel). Legends in common with panels g-j. **Right-top:** distributions of the coordinates of vertebrate (purple) and non-vertebrate (gold) meta-samples along the first component of the underlying PCA. The p-value of a Wilcoxon rank-sum test between the two distributions is reported. **g-j.** PCAs as in Fig. 1b but performed considering as expression measure for each species in each orthogroup (g) the average expression among all its paralogs, (h) the summed expression among all its paralogs, (i) the expression of a randomly selected paralogs, or (j) when limiting the analysis to single-copy orthogroups. Abbreviations: SE: single-end, PE: paired-end.

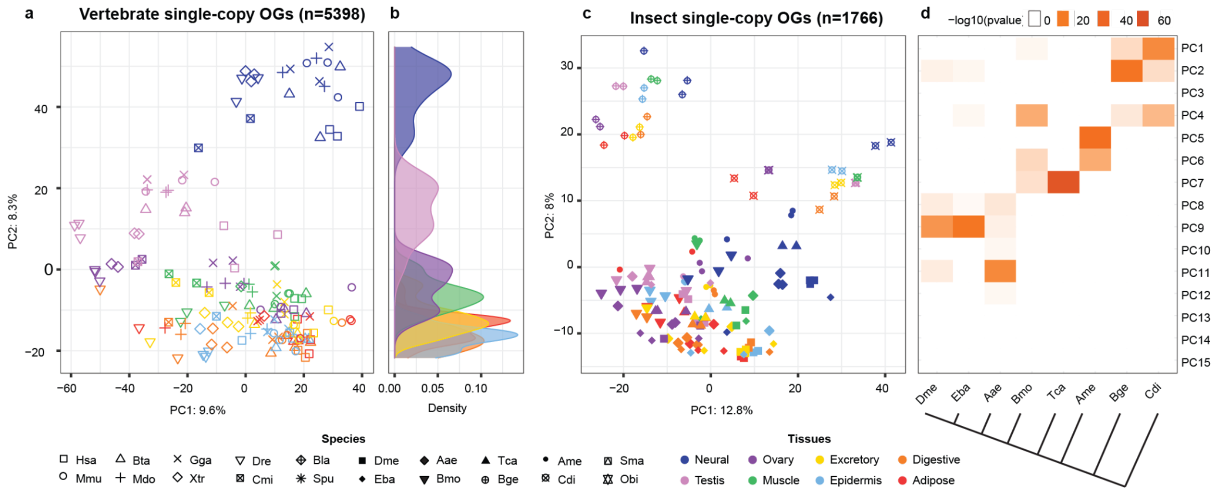


**Supplementary Fig. 3**: **a,c.** PCAs as in Fig. 1b but performed on the single-copy orthogroups (OGs) conserved in all vertebrate (a) or insect (c) species derived from the relative clade-specific gene orthologies (see **Methods**). Shapes and colors respectively represent the species and the tissue of the corresponding meta-sample. **b.** Distributions of the coordinates of each tissue's meta-samples (represented by color) on the second principal component (y axis) of the PCA shown in panel a. **d.** -log10(p-value) of ANOVA tests comparing the coordinates of meta-samples of different species along the top fifteen principal components from the PCA shown in panel c. Each test evaluates if there is a significant difference between any query species (i.e., column) versus all other collapsed species. All tests are performed with the *aov* function in R, and p-values were Bonferroni corrected. The phylogenetic tree on the bottom highlights the phylogenetic relationships between all insect species.

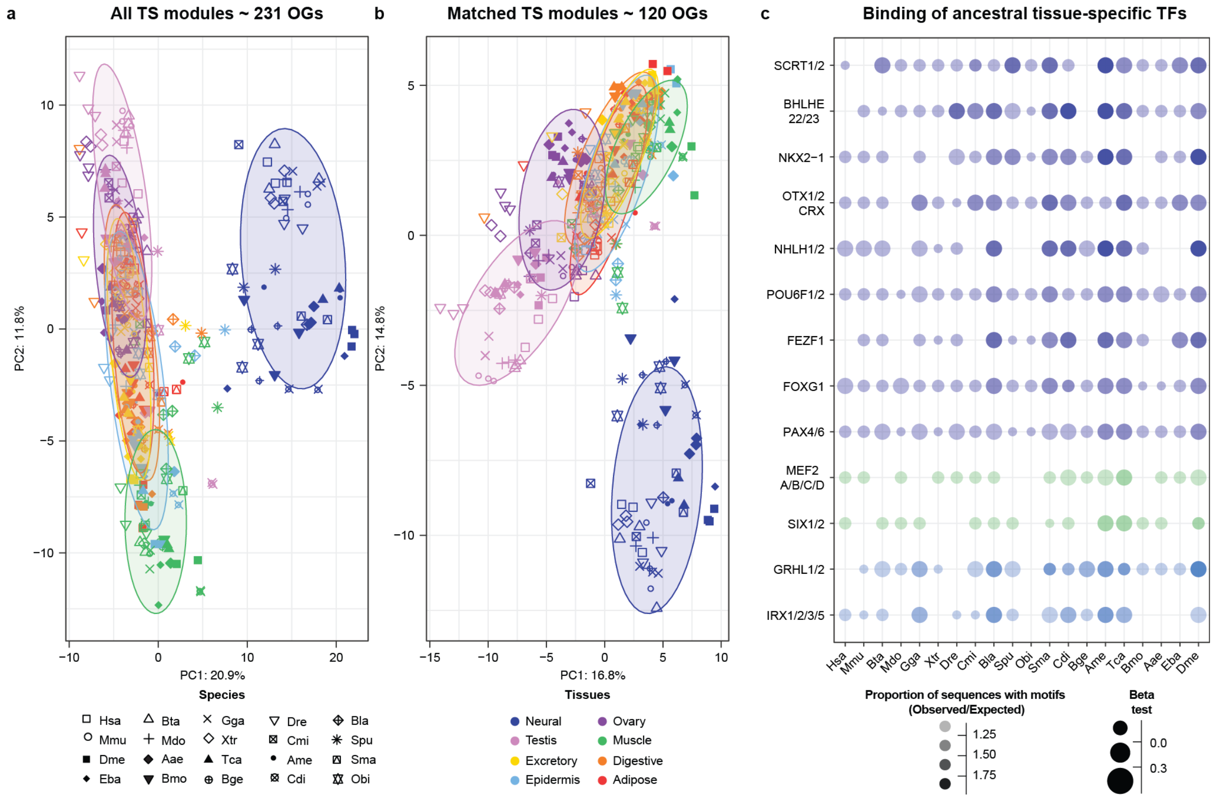


**Supplementary Fig. 4**: **a,b**. PCAs as in Fig. 1b but performed exclusively considering (a) the gene orthogroups in all bilaterian tissue-specific modules (see **Fig. 2b**) or (b) a matched number of gene orthogroups from each tissue's modules (maximum of 20 random orthogroups per module). Only the 2,436 gene orthogroups conserved in all species were considered. The ellipses encompass 80% of the meta-samples of each tissue (*stat_ellipse* function from *ggplot2*). **d.** Enrichment of binding sites across species for the TFs reported in **Fig. 2j**. The size of the dots represents the beta from a regression test checking if the number of binding sites per sequence is increased among the genes in the tested module compared to the species background (i.e., all species bilaterian-conserved genes). The transparency of the dots reflects the ratio between the proportion of sequences with at least one predicted binding site in the tested module (observed) and the proportion in the background (expected). Only cases for which the observed/expected ratio is greater than one are represented.

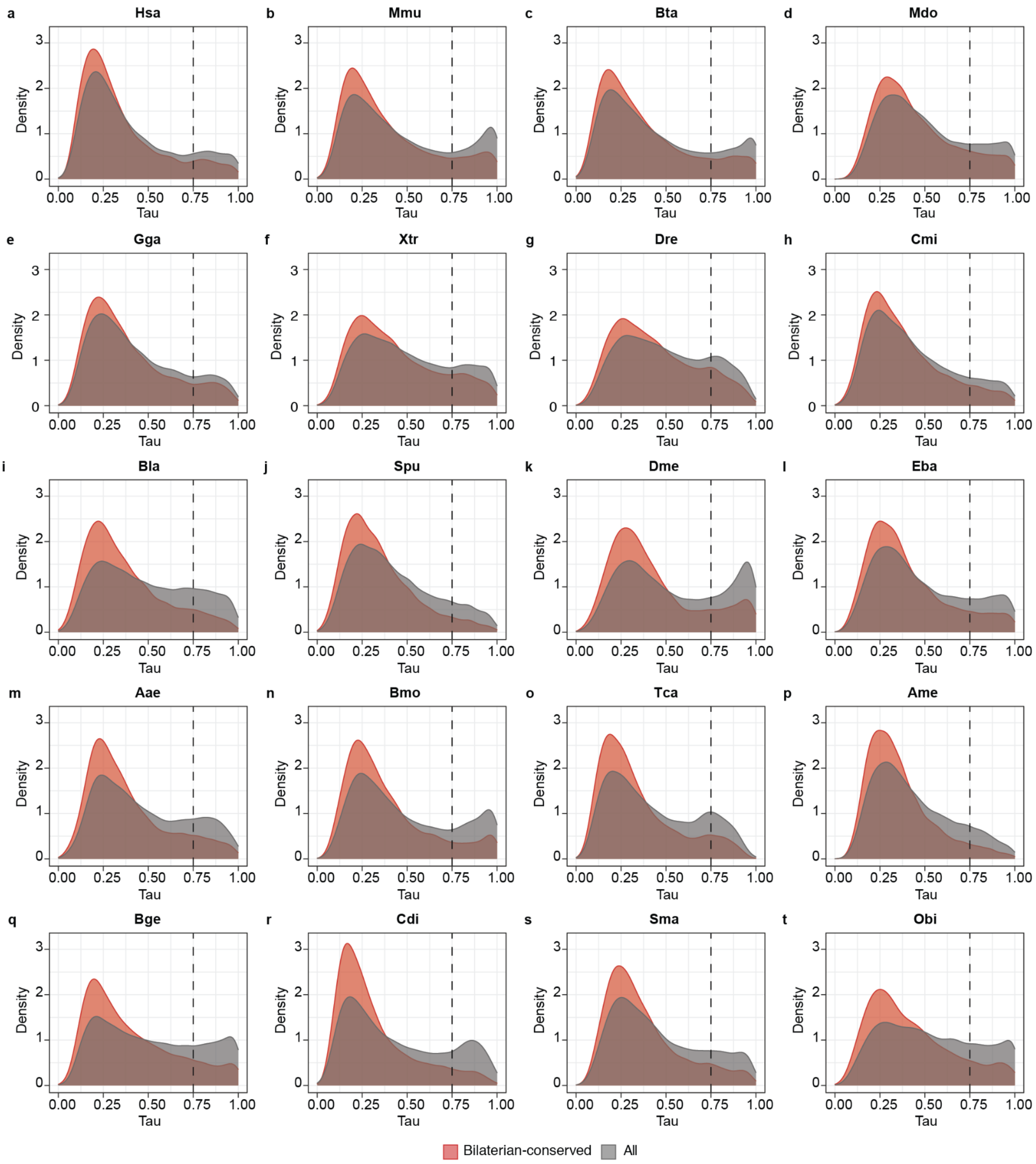


**Supplementary Fig. 5: a-t.** Tau distributions of bilaterian-conserved (red) or all (gray) protein-coding genes passing the expression cutoff in each species. Dashed line marks the selected cutoff for tissue-specificity (Tau=0.75).

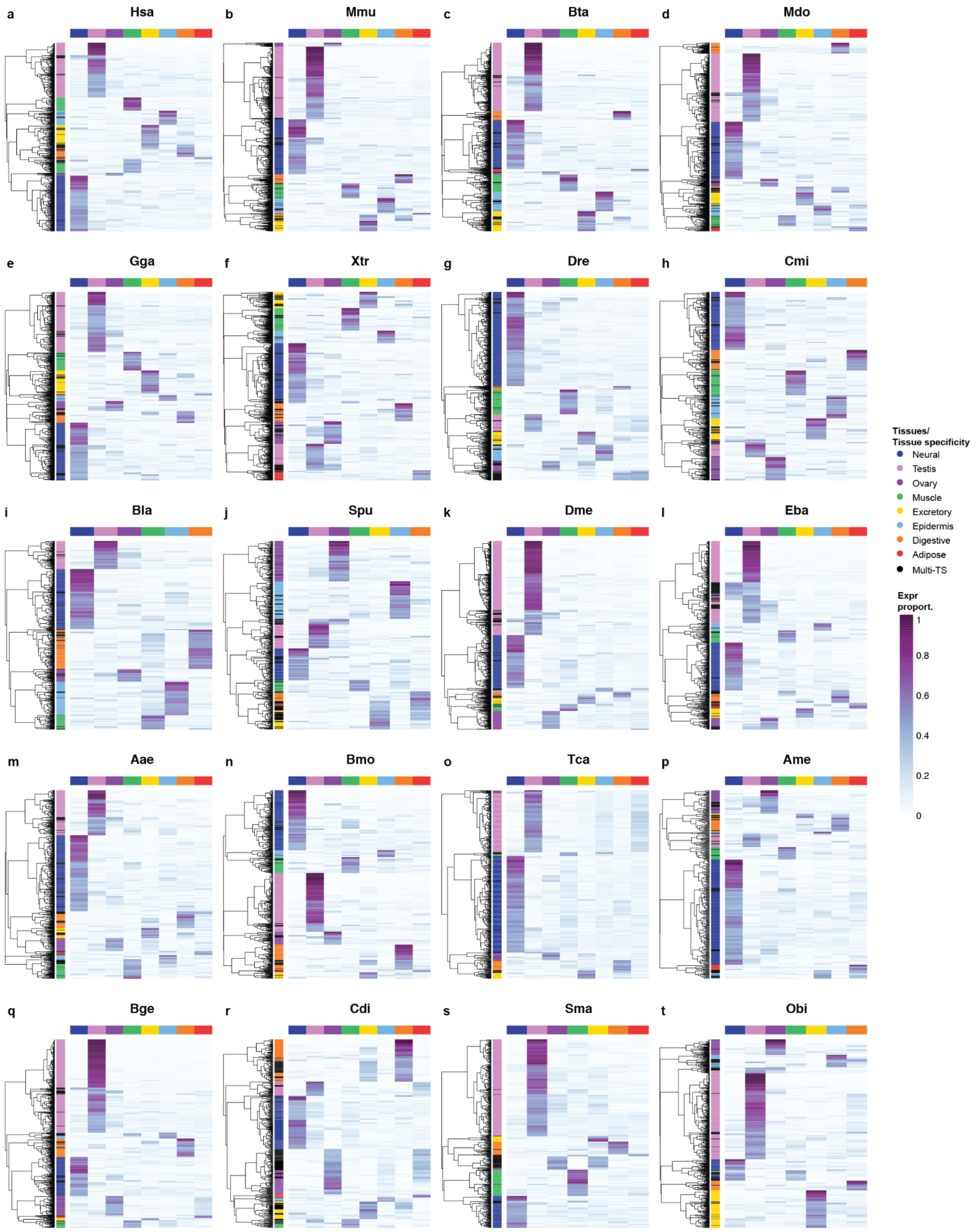


**Supplementary Fig. 6: a-t.** Heatmaps showing the clustering of bilaterian-conserved, tissue-specific genes (rows) based on their expression proportion (tissue_expr / all_tissue_expr) across tissues (columns) in each species. The heatmaps were generated by the *pheatmap* function in R with default clustering parameters.

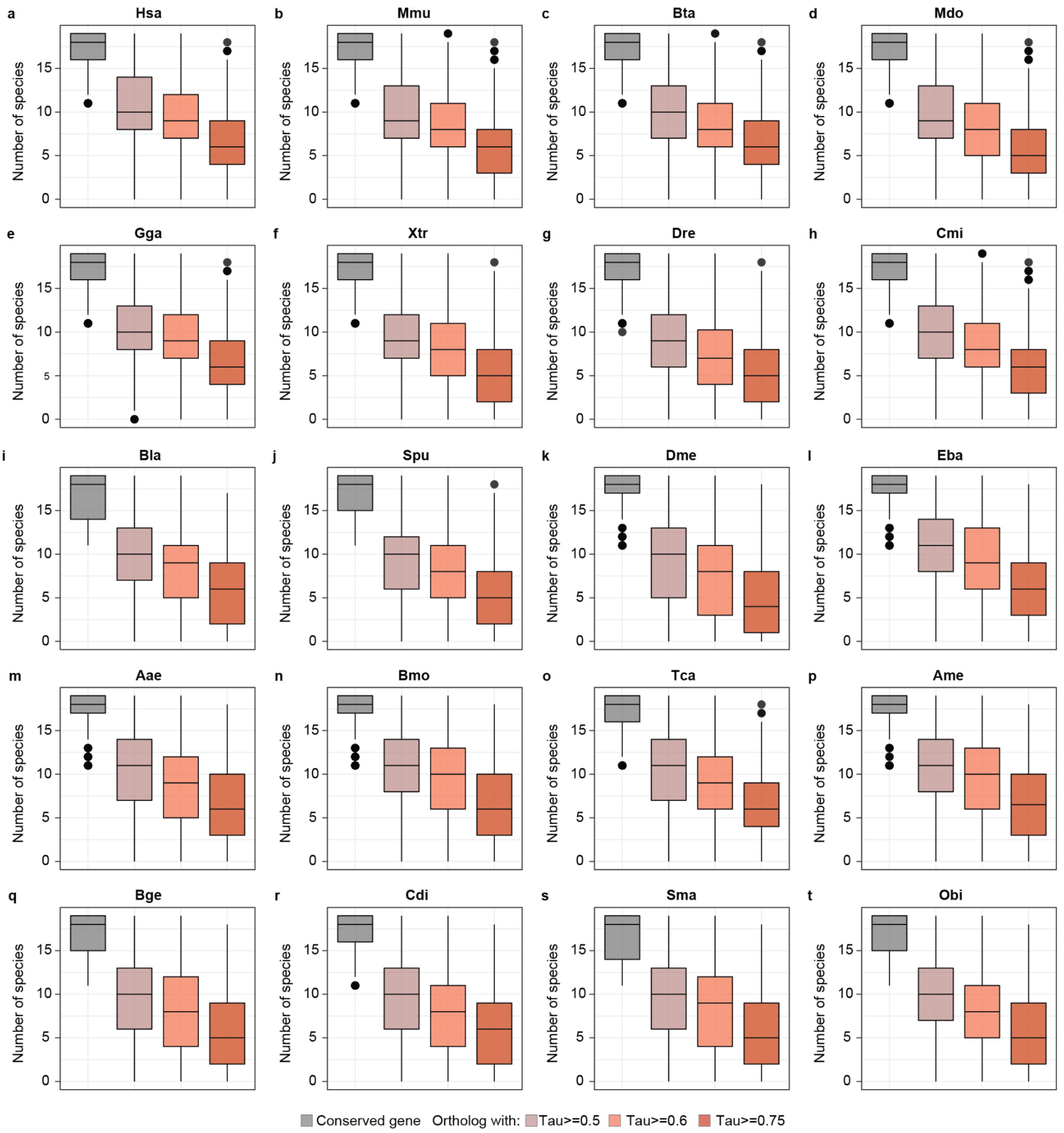


**Supplementary Fig. 7: a-t.** Distribution of the number of species in which each species' bilaterian-conserved, tissue-specific genes have at least one ortholog (gray) and this ortholog(s) has a Tau value higher than the specific cutoff (0.5, 0.6 and 0.75; other shades). The complete bilaterian-conserved orthogroups were considered in this analysis.

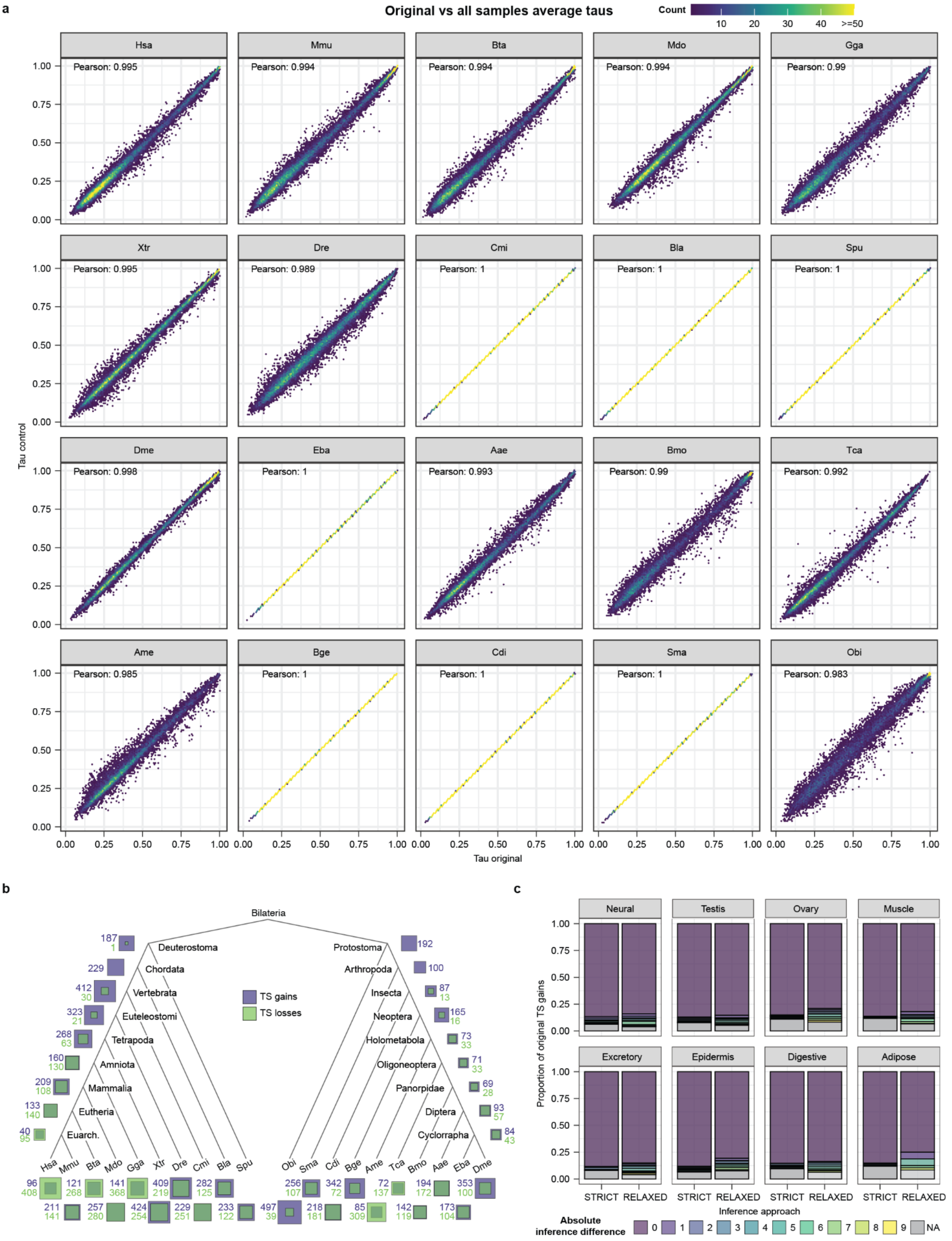

**Supplementary Fig. 8: a.** Comparison between original (x axis) and alternative (y axis) Tau values. Original Taus are computed by considering for each tissue the average expression across meta-samples, while alternative Taus consider the average expression across all samples (see **Methods**). The Pearson's correlation coefficient between original and alternative Taus for each species is reported in the relative panel. **b.** Total numbers of tissue-specificity gains and losses across all nodes and species inferred based on the alternative Taus. **c.** Concordance between the original inferences of tissue-specificity gains and the same inference based on alternative Taus. Original tissue-specificity gains are binned based on the absolute difference in phylogenetic positions with the corresponding alternative inferences. NA values indicate that the original inference does not have any corresponding inferences in the control dataset.

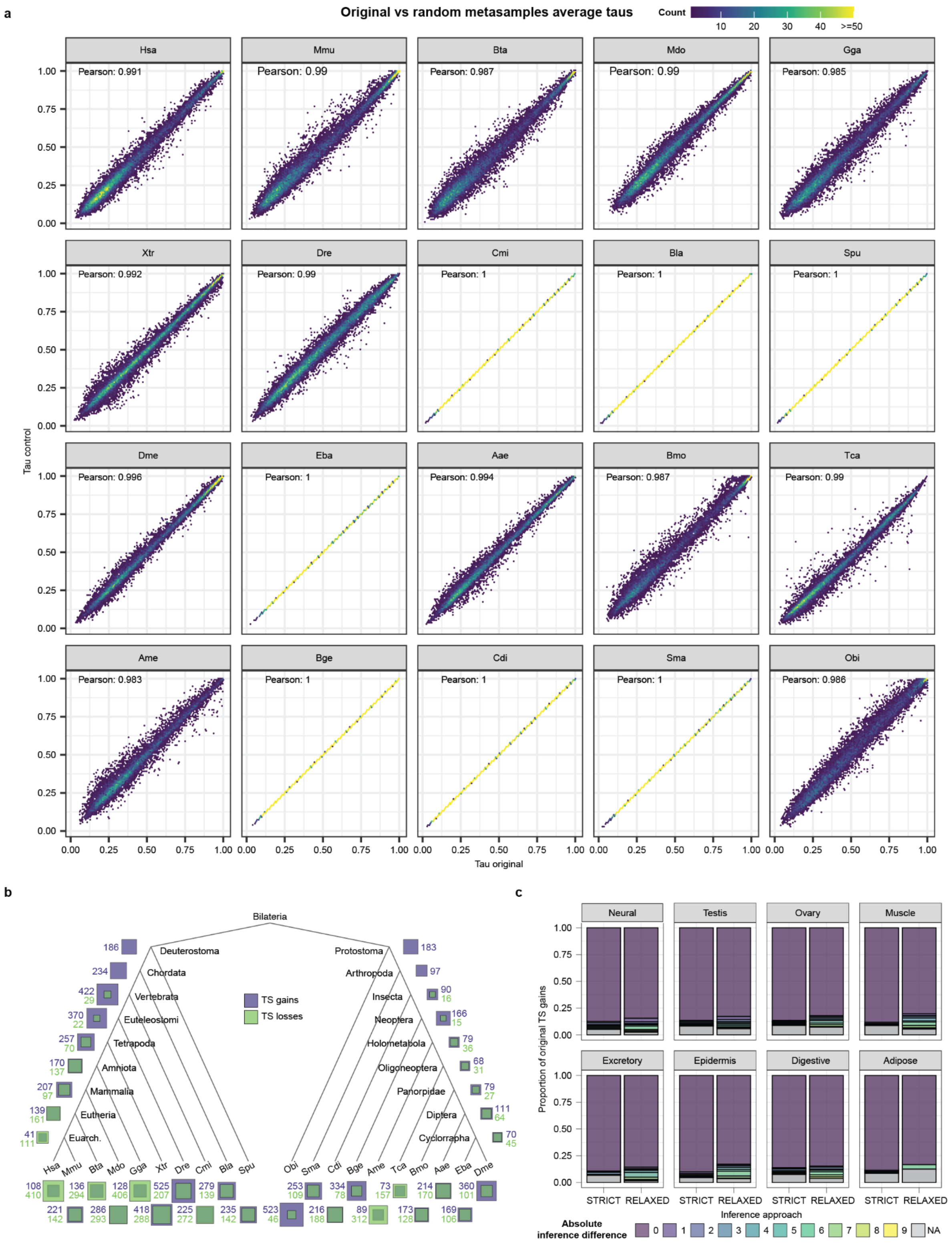


**Supplementary Fig. 9**: **a.** Comparison between original (x axis) and alternative (y axis) Tau values. Original Taus are computed by considering for each tissue the average expression across meta-samples, while alternative Taus are calculated after randomizing the sample to meta-sample correspondences within each tissue (see **Methods**). The Pearson's correlation coefficient between original and alternative Taus for each species is reported in the relative panel. **b.** Total numbers of tissue-specificity gains and losses across all nodes and species inferred based on the alternative Taus. **c.** Concordance between the original inferences of tissue-specificity gains and the same inference based on alternative Taus. Original tissue-specificity gains are binned based on the absolute difference in phylogenetic positions with the corresponding alternative inferences. NA values indicate that the original inference does not have any corresponding inferences in the control dataset.

**a**

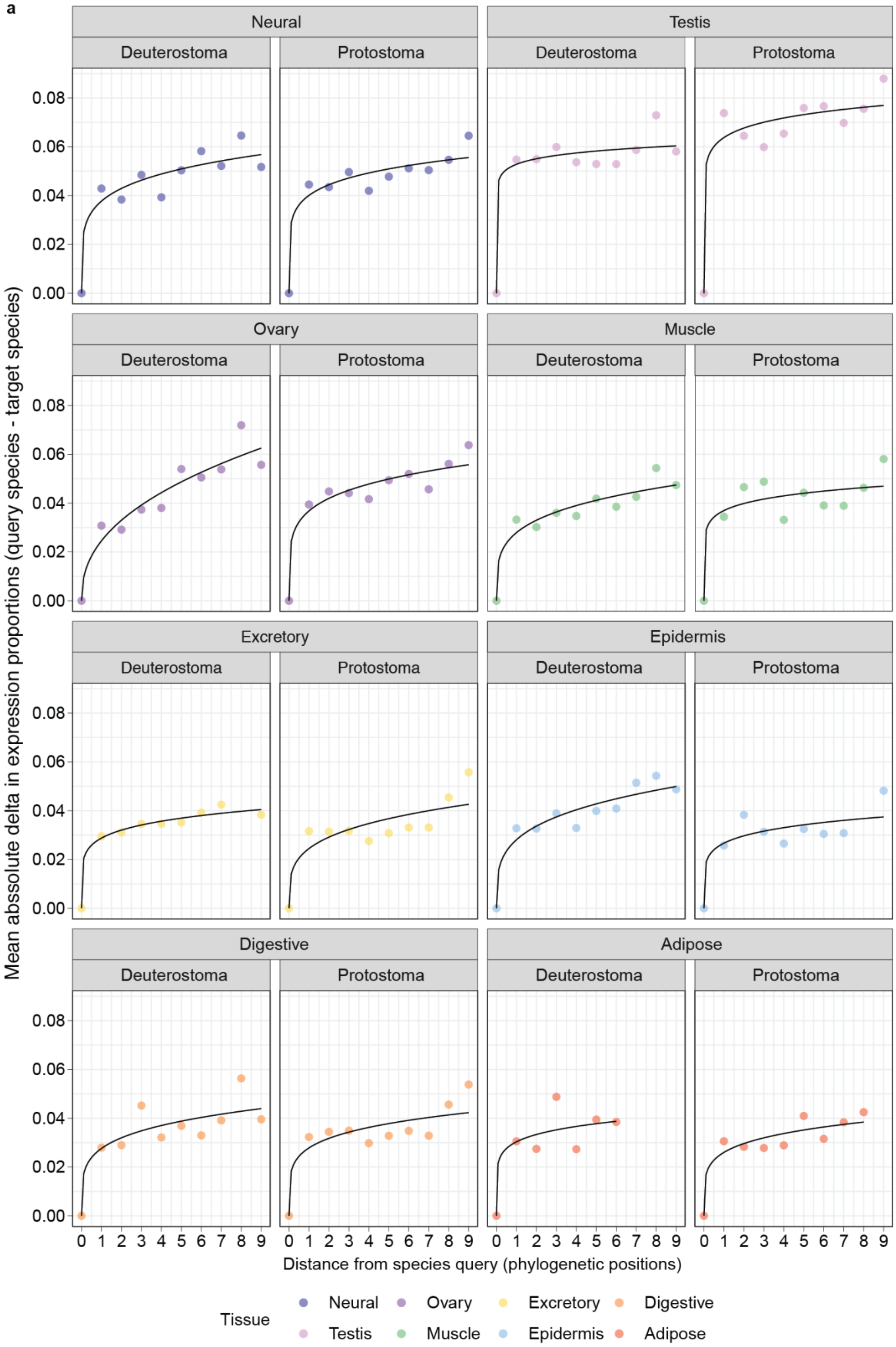


**Supplementary Fig. 10: a.** Expression divergence of the query species on each phylogenetic branch (Hsa for the deuterostomes, Dme for the protostomes) from all other species on the same branch in function of the number of phylogenetic positions that separate them (e.g., 1 corresponds to Mmu/Eba, 2 to Bta/Aae, etc). Expression divergence in each tissue is represented by the absolute deltas of expression proportions between the two compared species, averaged among all best-ancestral orthogroups. The nonlinear regression with formula $y = ax^k$ was fitted as in [20].

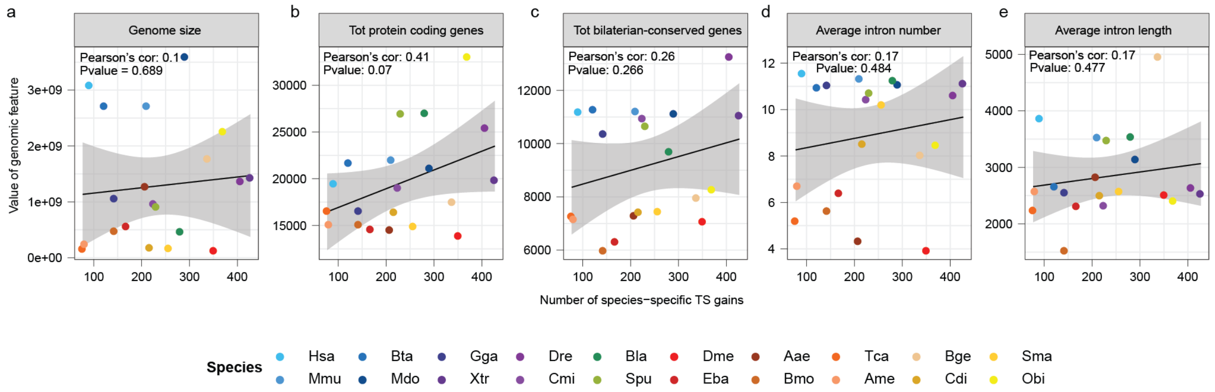


**Supplementary Fig. 11**: **a-e.** Correlation of total number of species-specific, tissue-specific gains with different genomic features. Values for all these genomic features are reported in **Supplementary Table 2**.

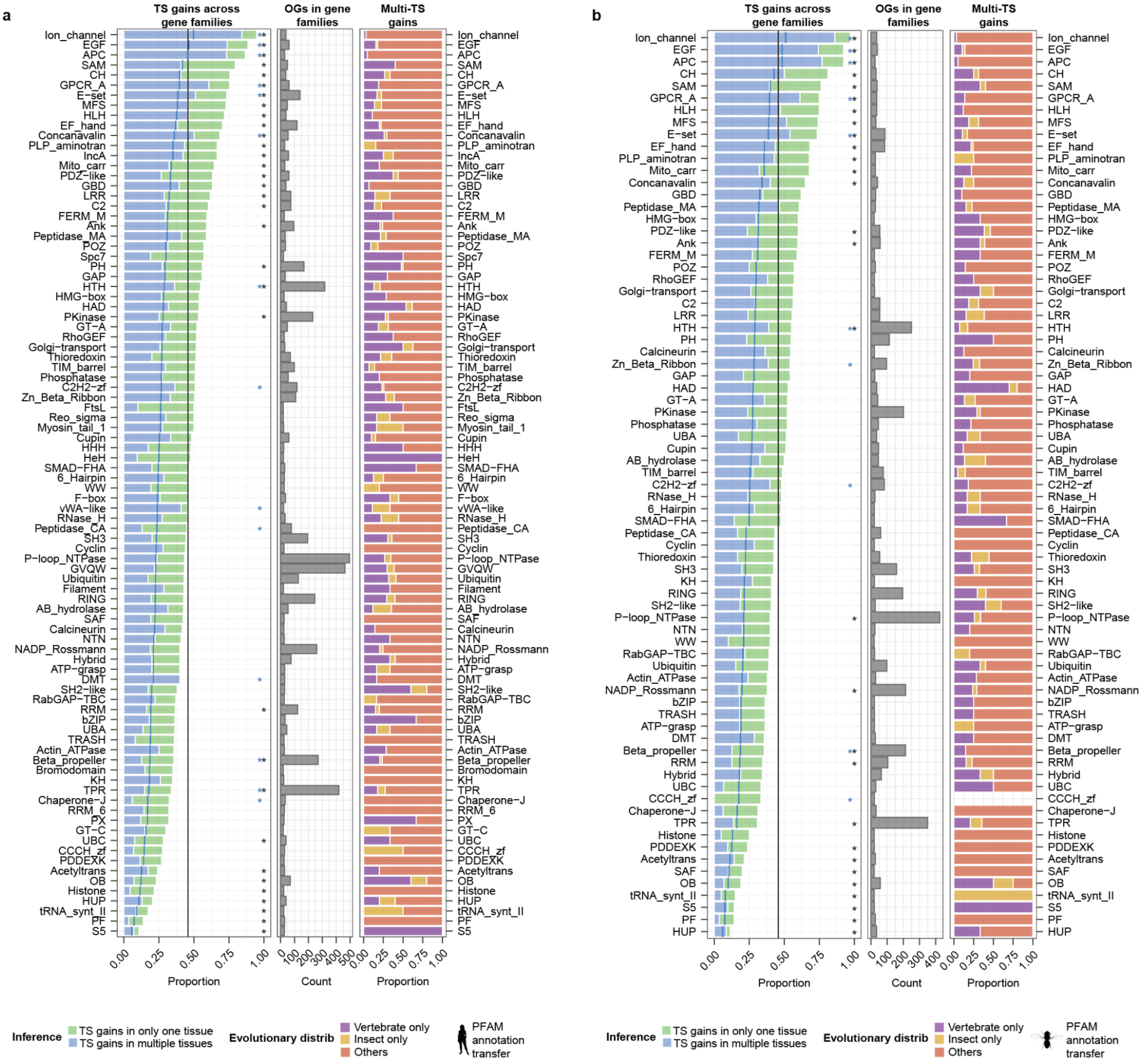


**Supplementary Fig. 12**: **a. Left:** proportions of orthogroups with tissue-specificity gains in only one tissue (single-TS; green) or in multiple tissues (multi-TS; blue) by gene family, ordered by the total proportion of orthogroups with tissue-specificity gains. The gene family classification derives from the human PFAM annotation (see **Methods**). The vertical lines represent the expected proportions of orthogroups with any tissue-specificity gain (black) or multi-TS gains (blue) in each gene family, corresponding to (1) the proportions of all bilaterian-conserved orthogroups with any tissue-specificity gains and (2) the proportion of these orthogroups where the gains are multi-TS. Black and blue asterisks indicate significant binomial tests (either enrichment or depletion; p-value ≤ 0.05) compared to the corresponding background for all and multi-TS gains, respectively. **Middle**: Number of gene orthogroups across gene families. **Right**. Proportions of multi-TS gains in each gene family based on their phylogenetic distribution. Vertebrate/Insect only: the multi-TS gains only occur in vertebrate/insect species or Vertebrata/Insecta and more recent ancestral nodes. Others: any other phylogenetic combination. **b.** Same as panel a, but with gene family classification derived from the fruit fly PFAM annotation.

**Supplementary Tables captions**

**Supplementary Table 1:** Annotation corrections. Catalog of corrected broken genes, resolved and unresolved chimeric genes, together with the correspondences between original and new GeneIDs.

**Supplementary Table 2:** Gene orthogroups statistics. Statistics of protein coding genes from each species included in the gene orthogroups before and after correction and filtering, together with the statistics of bilaterian-conserved orthogroups. The 2R-ohnolog groups from [6] used for one of the correction steps are reported. Original and corrected gene orthogroups files are provided in Supplementary Dataset, together with separate files for bilaterian-conserved orthogroups.

**Supplementary Table 3:** RNA-seq metadata information. For each species and tissue, it includes a list of all samples, sample to meta-sample correspondence, sample origin (i.e., public or in-house), SRA identifier, BioProject identifier, read number, read length, sequencing strategy (i.e. paired or single end), sequencing technology and mapping statistics.

**Supplementary Table 4:** Ancestral bilaterian tissue-specific modules. It provides a list of orthogroups IDs, human and fruit fly best-hits gene symbols for all the orthogroups included in each module.

**Supplementary Table 5:** Neural and testis phenotypes in human, mouse and fruit fly associated with, respectively, neural and testis ancestral bilaterian tissue-specific modules. Neural phenotypes were defined as anything matching "neuro", "behavior", "brain", "glia", "CNS" (case insensitive), while testis phenotypes were defined as anything matching "sperm", "infert", "sterile", "testis" (case insensitive).

**Supplementary Table 6:** GO enrichments for each bilaterian ancestral tissue-specific module as provided by gprofiler2 and using the GO transfers derived from the human GO annotation. All p-values were FDR corrected.

**Supplementary Table 7:** GO enrichments for each bilaterian ancestral tissue-specific module as provided by gprofiler2 and using the GO transfers derived from the fruit fly GO annotation. All p-values were FDR corrected.

**Supplementary Table 8:** GO enrichments for non-tissue-specific orthogroups as provided by gprofiler2. All p-values were FDR corrected.

**Supplementary Table 9:** Tissue specificity gains and losses throughout the phylogeny. For each tissue, it provides a list of inferences that include the orthogroup ID, the relative node/species, the inference type (i.e., gain or loss) and the inference criteria (i.e., strict, relaxed or merged).

**Supplementary Table 10:** GO enrichments for orthogroups with tissue-specific gains in each node and species, as provided by gprofiler2 and using the GO transfers derived from the human GO annotation. We grouped GO enrichment coming from tissue-specific gains in the same tissue, adding a column reporting the relative node/species. All p-values were FDR corrected.

**Supplementary Table 11:** GO enrichments for orthogroups with tissue-specific gains in Vertebrata or more recent nodes and vertebrate species, as provided by gprofiler2 and using the GO transfers derived from the vertebrate-specific GO annotation. We grouped GO enrichment coming from tissue-specific gains in the same tissue, adding a column reporting the relative node/species. All p-values were FDR corrected.

**Supplementary Table 12:** GO enrichments for orthogroups with tissue-specific gains in Insecta or more recent nodes and insect species, as provided by gprofiler2 and using the GO transfers derived from the insect-specific GO annotation. We grouped GO enrichment coming from tissue-specific gains in the same tissue, adding a column reporting the relative node/species.All p-values were FDR corrected.

**Supplementary Table 13:** GO enrichments for orthogroups with tissue-specific gains (as provided by gprofiler2 and using the GO transfers derived from the human GO annotation) uniquely enriched in each node and species compared to all other gains in the same tissue. We grouped GO enrichment coming from tissue-specific gains in the same tissue, adding a column reporting the relative node/species. All p-values were FDR corrected.

**Supplementary Dataset content**

The Supplementary Dataset associated with this manuscript is available at https://data.mendeley.com/datasets/22m3dwhzk6/2.

**Samples_Quality_Control**

- **FastQC_output.zip:** zip archive containing the fastQC html reports for all the RNA-seq samples generated in the context of this study.
- **All_species-mapping_stats.txt:** files containing mapping statistics for all samples in all species, as returned by *Kallisto* [13].

**Gene_Orthology_Files**

- **All_orthogroups.tab:** gene orthogroups returned by a *Broccoli* [4] run on the twenty bilaterian species.
- **All_orthogroups-corrected_and_filtered.tab:** gene orthogroups returned by a *Broccoli* [4] run on the twenty bilaterian species, corrected and filtered as described in the Supplementary Methods (see “Post-orthology genome annotation and gene orthogroups refinements”).
- **Bilateria_conserved_orthogroups-EXPR_genes.tab:** all bilaterian-conserved orthogroups filtered for expressed genes (see “Post-orthology genome annotation and gene orthogroups refinements” in Supplementary Methods).
- **Bilateria_conserved_orthogroups-Best_Ancestral_OGs.tab:** bilaterian-conserved orthogroups filtered only for the best-ancestral orthologs.
- **Insecta_all_conserved_orthogroups-single_copies.tab:** gene orthogroups returned by a *Broccoli* [4] run on the eight insect species. Only filtered single-copy orthogroups conserved in all species are reported (see Methods).
- **Vertebrata_all_conserved_orthogroups-single_copies.tab:** gene orthogroups returned by a *Broccoli* [4] run on the eight vertebrate species. Only filtered single-copy orthogroups conserved in all species are reported (see Methods).
- **Bilateria_conserved_orthogroups-All_genes-associated_tissues.tab:** bilaterian-conserved orthogroups where each gene is associated with information relative to its Tau, associated tissue(s), top tissue(s) and expression cutoff status.
- **Bilateria_conserved_orthogroups-Best_TS_OGs-Neural.tab:** bilaterian-conserved orthogroups with selected best-TS orthologs for Neural.
- **Bilateria_conserved_orthogroups-Best_TS_OGs-Testis.tab:** bilaterian-conserved orthogroups with selected best-TS orthologs for Testis.
- **Bilateria_conserved_orthogroups-Best_TS_OGs-Ovary.tab:** bilaterian-conserved orthogroups with selected best-TS orthologs for Ovary.
- **Bilateria_conserved_orthogroups-Best_TS_OGs-Muscle.tab:** bilaterian-conserved orthogroups with selected best-TS orthologs for Muscle.
- **Bilateria_conserved_orthogroups-Best_TS_OGs-Excretory.tab:** bilaterian-conserved orthogroups with selected best-TS orthologs for Excretory.
- **Bilateria_conserved_orthogroups-Best_TS_OGs-Digestive.tab:** bilaterian-conserved orthogroups with selected best-TS orthologs for Digestive
- **Bilateria_conserved_orthogroups-Best_TS_OGs-Epidermis.tab:** bilaterian-conserved orthogroups with selected best-TS orthologs for Epidermis.
- **Bilateria_conserved_orthogroups-Best_TS_OGs-Adipose.tab:** bilaterian-conserved orthogroups with selected best-TS orthologs for Adipose.

**Expression_Matrices**

- **Bilateria_conserved_orthogroups-metasample_expr-Best_Ancestral-NORM.tab:** normalized expression matrix for all best-ancestral orthogroups (meta-sample level).

- **Bilateria_conserved_orthogroups-metasample_expr-paralogs_average-NORM.tab:** normalized expression matrix where the expression measure for each species in each orthogroup derives from the average expression of all its paralogs (meta-sample level).
- **Bilateria_conserved_orthogroups-metasample_expr-paralogs_summed-NORM.tab:** normalized expression matrix where the expression measure for each species in each orthogroup derives from the summed expression of all its paralogs (meta-sample level).
- **Bilateria_conserved_orthogroups-metasample_expr-paralogs_random-NORM.tab:** normalized expression matrix where the expression measure for each species in each orthogroup corresponds to the expression of a random paralog (meta-sample level).
- **Bilateria_conserved_orthogroups-metasample_expr-single_copies-NORM.tab:** normalized expression matrix where only single-copy orthogroups are included (meta-sample level).
- **Insecta_ALL_conserved_orthogroups-metasample_expr-single_copies-NORM.tab:** normalized expression matrix containing insect single-copy orthogroups conserved in all species (meta-sample level).
- **Vertebrata_ALL_conserved_orthogroups-metasample_expr-single_copies-NORM.tab:** normalized expression matrix containing vertebrate single-copy orthogroups conserved in all species (meta-sample level).
- **Bilateria_conserved_orthogroups-tissue_expr-Best_Ancestral-ZSCORED.tab:** z-scored expression matrix for all best-ancestral orthogroups (tissue level).
- **Bilateria_conserved_orthogroups-tissue_expr-paralogs_average-ZSCORE.tab:** z-scored expression matrix where the expression measure for each species in each orthogroup derives from the average expression of all its paralogs (tissue level).
- **Bilateria_conserved_orthogroups-tissue_expr-paralogs_summed-ZSCORE.tab:** z-scored expression matrix where the expression measure for each species in each orthogroup derives from the summed expression of all its paralogs (tissue level).
- **Bilateria_conserved_orthogroups-tissue_expr-paralogs_random-ZSCORE.tab:** z-scored expression matrix where the expression measure for each species in each orthogroup corresponds to the expression of a random paralog (tissue level).

**GO_Files**
- **GO_transfer_from_Hsa-Bilateria_conserved_OGs.txt:** GO annotation of bilaterian-conserved orthogroups as transferred from the human GO annotation.
- **GO_transfer_from_Dme-Bilateria_conserved_OGs.txt:** GO annotation of bilaterian-conserved orthogroups as transferred from the fruit fly GO annotation.
- **GO_transfer_Insecta_specific-Bilateria_conserved_OGs.txt:** GO annotation of bilaterian-conserved orthogroups as transferred from the insect-specific GO annotation.
- **GO_transfer_Vertebrata_specific-Bilateria_conserved_OGs.txt:** GO annotation of bilaterian-conserved orthogroups as transferred from the vertebrate-specific GO annotation.

**GTFs:**
- **"species"_annot-B-brochi.gtf:** Corrected GTF file for the relative species.

**Motif_files:**
- **Clustered_motifs_annotation_to_OG.tab:** annotation of bilaterian-conserved orthogroups and relative domains.
- **Clustered_motifs_filtered.pfm:** positional frequency matrix of clustered motifs.

**Phenotypic_info:**
- **all_species-Bilateria_conserved-conservation_labels_AND_phenotypic_info.txt:** functional annotation of bilaterian-conserved orthogroups derived from the human, mouse and fruit fly phenotypic annotations.

**Specialization_supporting_plots:**

- **all_ancestors_gains-expression_between_TS_and_nonTS_species.pdf:** expression of best-TS orthologs between species with and without tissue-specificity in corresponding ancestral gains.
- **all_species_gains-expression_between_TS_and_nonTS_species.pdf:** expression of best-TS orthologs between species with and without tissue-specificity in corresponding species-specific gains.

### References


1. Kim, D., Paggi, J. M., Park, C., Bennett, C. & Salzberg, S. L. Graph-based genome alignment and genotyping with HISAT2 and HISAT-genotype. *Nat. Biotechnol.* **37**, 907–915 (2019).
2. Pertea, M. *et al.* StringTie enables improved reconstruction of a transcriptome from RNA-seq reads. *Nat. Biotechnol.* **33**, 290–295 (2015).
3. Haas, B., Papanicolaou, A. & Others. TransDecoder (find coding regions within transcripts). *Google Scholar* (2016).
4. Derelle, R., Philippe, H. & Colbourne, J. K. Broccoli: Combining Phylogenetic and Network Analyses for Orthology Assignment. *Mol. Biol. Evol.* **37**, 3389–3396 (2020).
5. Katoh, K. & Standley, D. M. MAFFT multiple sequence alignment software version 7: improvements in performance and usability. *Mol. Biol. Evol.* **30**, 772–780 (2013).
6. Touceda-Suárez, M. *et al.* Ancient Genomic Regulatory Blocks Are a Source for Regulatory Gene Deserts in Vertebrates after Whole-Genome Duplications. *Mol. Biol. Evol.* **37**, 2857–2864 (2020).
7. Bertram, J. *et al.* CAGEE: computational analysis of gene expression evolution. *Mol. Biol. Evol.* **40**, (2023).
8. Chen, J. *et al.* A quantitative framework for characterizing the evolutionary history of mammalian gene expression. *Genome Res.* **29**, 53–63 (2019).
9. Bedford, T. & Hartl, D. L. Optimization of gene expression by natural selection. *Proceedings of the National Academy of Sciences* **106**, 1133–1138 (2009).
10. Brawand, D. *et al.* The evolution of gene expression levels in mammalian organs. *Nature* **478**, 343–348 (2011).
11. Fukushima, K. & Pollock, D. D. Amalgamated cross-species transcriptomes reveal organ-specific propensity in gene expression evolution. *Nat. Commun.* **11**, 4459 (2020).
12. Khabbazian, M., Kriebel, R., Rohe, K. & Ané, C. Fast and accurate detection of evolutionary shifts in Ornstein–Uhlenbeck models. *Methods Ecol. Evol.* **7**, 811–824 (2016).
13. Bray, N. L., Pimentel, H., Melsted, P. & Pachter, L. Near-optimal probabilistic RNA-seq quantification. *Nat. Biotechnol.* **34**, 525–527 (2016).
